# Architecture and Function of Holocentric CENP-A-Independent Kinetochores

**DOI:** 10.1101/2025.07.14.664805

**Authors:** Christine Yu, Sundar Ram Sankaranarayanan, Gaetan Cornilleau, Anna Howes, Caleigh Azumaya, Inna Zilberleyb, Bobby Brillantes, Tommy K. Cheung, Leonie Dec, Damarys Loew, Phong Tran, Christopher M. Rose, Ines Anna Drinnenberg, Claudio Ciferri, Stanislau Yatskevich

**Author notes:** Corresponding authors: Ines Anna Drinnenberg, Claudio Ciferri, Stanislau Yatskevich. These authors jointly supervised this work (Ines Anna Drinnenberg, Stanislau Yatskevich).

## Abstract

Kinetochores are essential macromolecular complexes that anchor chromosomes to the mitotic spindle to ensure faithful cell division^1^. Despite their critical role, the structural organization of kinetochores assembled on centromeres with vastly distinct architectures across diverse species remains poorly understood^2,3^. To address this question, we determined the cryo-EM structures of the inner kinetochore (CCAN) from the silkmoth *Bombyx mori*, an insect that lacks the canonical centromere-specifying histone variant CENP-A and exhibits chromosome-wide centromeric activity (holocentric). Our analysis reveals that *B. mori* CCAN assembles via atypical histone-fold protein dimerization into a self-contained, head-to-head dimer that topologically entraps and loops DNA, creating a point-centromere-like architecture. This structure also incorporates four previously uncharacterized Centromeric Subunit proteins that are evolutionarily repurposed from the outer kinetochore Dam1/DASH complex. Our work establishes this self-contained CCAN dimer as a key structural unit that forms the basis of a holocentric organization and suggests that large-scale centromere architectures can emerge from the modular arrangement of such discrete kinetochore units.

## Introduction

Centromeres are specialized chromosomal regions responsible for kinetochore assembly and spindle attachment during chromosome segregation^1,4^. In most species, centromeres incorporate a specialized histone H3 variant called CENP-A which assembles into a nucleosome (CENP-A^Nuc^)^5^. The CENP-A^Nuc^ lays the foundation for kinetochore assembly by recruiting the essential inner kinetochore known as the Constitutive Centromere-Associated Network (CCAN)^6–8^. In the case of point centromeres found in a few unicellular yeast species, a single CENP-A^Nuc^ is recruited to the centromere in a DNA-sequence-specific manner and recognized directly and specifically by CCAN and its associated proteins^9–14^. In the case of regional centromeres present in many other eukaryotes, including humans, many copies of CENP-A^Nuc^ are distributed over several hundred kilobases of DNA to recruit numerous CCAN complexes^15,16^. The CENP-C protein is the main scaffolding component that directly decodes the CENP-A^Nuc^ and recruits the rest of the CCAN to the centromere^17–24^. Human CCAN primarily recognizes and binds the linker DNA between nucleosomes via a central CENP-LN DNA-binding channel^19^. Across eukaryotes, CCAN recruits and directly binds to the Knl1-Mis12-Ndc80 (KMN) network, an outer kinetochore complex that mediates microtubule attachment^25^.

Centromere organization and kinetochore composition exhibit considerable diversity across eukaryotes, highlighting their evolutionary plasticity despite their universally essential role in chromosome segregation^3,26–28^. This diversity even extends to the loss of components such as CENP-A and CENP-C, which are thought to act as cornerstones for kinetochore assembly. These findings raise an interesting question: how can kinetochores assemble and function without these key factors that anchor them at most centromeres, including those in humans^29,30^? Despite the loss of CENP-A/C, many of these species, including several insect orders, retain sequence-divergent CCAN subunits, suggesting the existence of alternative, yet functionally robust, pathways to sustain kinetochore-spindle interactions. Intriguingly, CENP-A/C-lacking insects also evolved a holocentric chromosome organization, where centromeric activity and kinetochore assembly are dispersed along the chromosomal length^30^. In the holocentric silkmoth *Bombyx mori* (Lepidoptera order), previous studies of inner (CENP-T) and outer (Dsn1) kinetochore components in asynchronous cell lines revealed that kinetochores assemble over large chromosomal domains scattered along entire chromosomes^31^. In mitosis, microscopy analyses showed that kinetochore staining appears as longitudinal bands along *B. mori* chromosomes consistent with the holocentric architecture^31^. The presence of large centromeric domains permissive to kinetochore formation, along with the band-like staining pattern along mitotic chromosomes, raises important questions about the molecular nature of holocentromeres. Are higher-order centromeres built using functionally segregated kinetochore units, or do kinetochores form an extended, cooperative structure spanning chromatin to coordinate its activity? These questions are also relevant for regional centromeres where genomic analyses have shown that CENP-A and other kinetochore components are enriched over several hundred kilobases of DNA^4,15^. Lastly, and more broadly, how can the essential and conserved process of chromosome segregation be accomplished using seemingly so many different types of centromere architectures?

To address these questions, and following up on previous work, we functionally, biochemically, and structurally characterized the *B. mori* kinetochore ^29–31^. Our work reveals how evolutionarily repurposed, novel, and conserved *B. mori* CCAN subunits assemble into a unique dimeric structure that folds DNA into a configuration poised for chromosome segregation. Remarkably, this dimeric arrangement mirrors point-centromere architecture, suggesting a conserved evolutionary principle that persists independently of CENP-A. These findings illuminate the kinetochore’s remarkable adaptability and provide a new framework for understanding how self-contained kinetochore “module units” can drive diverse centromere organization strategies across eukaryotes, enabling plasticity and evolutionary innovation.

### Conserved and divergent structural features of *Bombyx mori* CCAN

We recombinantly expressed and fully purified individual *B. mori* CCAN (bmCCAN) modules, including the evolutionarily diverged bmCENP-LN and bmCENP-HIKM modules, a unique bmCENP-T protein with an altered histone-fold domain (HFD)^32^, and a module we term bmCENP-OP, predicted to contain an RWD domain similar to its human (hsCENP-OP) and yeast (scCENP-OP) counterparts (Extended Data Fig. 1a)^29^. However, reconstitution experiments using purified components yielded only partial bmCCAN assembly (Extended Data Fig. 1b). To investigate whether incomplete assembly resulted from missing components or absent post-translational modifications, we leveraged the high sequence homology between *Spodoptera frugiperda,* another Lepidopteran insect, and *B. mori* CCAN proteins. Using a hybrid strategy, we overexpressed each *B. mori* CCAN module in *S. frugiperda* cells for semi-native purification (Extended Data Fig. 1 C). Mass spectrometry (MS) confirmed component exchange between species. This approach yielded a more stoichiometric bmCCAN assembly, with a few additional bands visible on the SDS-PAGE gel (Extended Data Fig. 1d, e).

Using cryogenic electron microscopy (cryoEM), we determined three distinct structures from this sample: (i) bmCENP-LN:HIKM:T sub-complex (bmCCAN^LN:HIKM:T^), (ii) bmCENP-LN:HIKM:T:OP complex (bmCCAN^LN:HIKM:T:OP^), and (iii) bmCENP-LN:HIKM:T:OP complex with additional pentameric coiled-coil density bound across bmCENP-LN:HIKM:OP interfaces (apo-bmCCAN) (Fig. 1a-c, Extended Data Fig. 2 and 3, Extended Data Table 1).

**Fig. 1.**
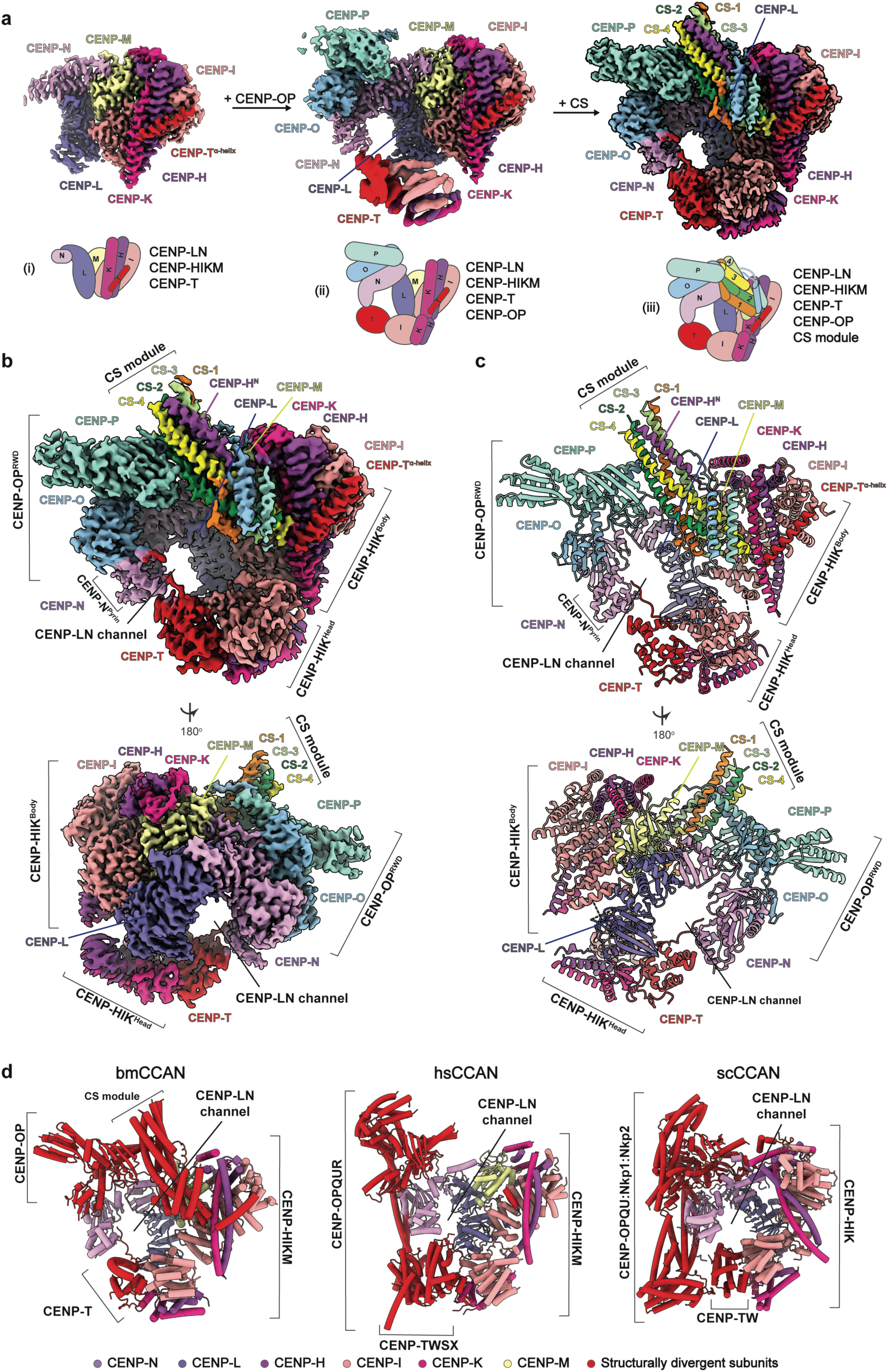
Structure of the *Bombyx mori* inner kinetochore/CCAN complex. **a,** CryoEM reconstructions of the three assembly intermediates observed in the bmCCAN dataset corresponding to (i) bmCENP-LN-HIKM-T subcomplex, (ii) bmCENP-LN-HIKM-T-OP subcomplex and (iii) bmCENP-LN-HIKM-T-OP-CS complex. A cartoon schematic of the subcomplex assembly is shown under each reconstruction. CENP-T^α-helix^ – N-terminal α-helix of CENP-T protein. **b,** Composite cryoEM reconstruction of the complete apo-bmCCAN complex composed of two combined cryoEM maps: locally refined bmCENP-HIK^Head^ map and the rest of the bmCCAN map (Supplementary Fig. 2). CENP-OP^RWD^- RWD domains of CENP-OP proteins. CENP-N^Pyrin^ – pyrin domain of CENP-N protein. **c,** A molecular model of the complete apo-bmCCAN complex. **d,** A comparison between apo-bmCCAN determined in this study with hsCCAN (PDB ID: 7R5S) and scCCAN (PDB ID: 8OVW) ^10,19^. CENP-LN and CENP-HIKM subunits are colored by subunit with the indicated color scheme, while structurally and sequence divergent subunits are colored in red for illustrative purposes.

The global architecture of apo-bmCCAN mirrors the homologous complexes in human and yeast, despite limited sequence conservation and distinct predicted folds between the homologous subunits (Fig. 1d). Apo-bmCCAN forms a ring-like structure, with the conserved CENP-LN channel serving as the central architectural feature (Fig. 1b-d). The bmCENP-HIKM module reinforces and stabilizes bmCENP-L, similar to human and yeast complexes. In contrast, the bmCENP-OP RWD domains (bmCENP-OP^RWD^) adopt a unique configuration, binding directly atop bmCENP-N and adding bulk density to the central channel (Fig. 1b-d). This channel is sealed by the altered HFD of bmCENP-T (bmCENP-T^HFD^) that connects bmCENP-HIK^Head^ with the bmCENP-N^Pyrin^ (head group and pyrin domains of bmCENP-HIK and bmCENP-N, respectively, Fig. 1b, c). The bmCENP-T^HFD^, structurally distinct from canonical histone-fold domains, likely compensates for the absence of the CENP-W protein found in most other species that encode CENP-T (Fig. 1d)^28,32^. BmCENP-T also makes an additional unique interaction with bmCENP-HIK^Body^ using an α-helix (bmCENP-T^α-helix^) that precedes the bmCENP-T^HFD^ (Fig. 1a-c).

In the apo-bmCCAN structure, the bmCENP-N:T interaction is poorly resolved, suggesting a dynamic interface that may serve as the primary DNA-entry gate into the central binding channel. Consistent with this, bmCENP-LN channel is fully open in the bmCCAN^LN:HIKM:T^ subcomplex reconstruction (Fig. 1a). Lastly, in the DNA-bound structure described later, the bmCENP-N:T interaction is rigidified, and an extended density is observed for the CENP-T N-terminal extension that folds along bmCENP-N^Pyrin^.

### Centromeric Subunit proteins form a central bmCCAN coiled-coil

The salient feature in our reconstruction is the presence of a pentameric coiled-coil nestled across most bmCCAN subunits, as this coiled-coil did not belong to any over-expressed components and must have been co-purified from the native source (Fig. 1b, c, Fig. 2a, b). The high resolution of our reconstruction permitted the sequencing of these proteins directly from the cryoEM maps using ModelAngelo (Methods, Extended Data Fig. 4a)^33^. One helix of the pentameric coiled-coils was predicted to belong to the N-terminus of the bmCENP-H protein (bmCENP-H^N^). Consistent with this assignment, continuous density was observed between bmCENP-H protein and bmCENP-H^N^ coiled-coil helix, validating our method (Fig. 1b, c). Sequences for the other four proteins of the heteromeric coiled-coil belonged to previously uncharacterized short proteins, which we termed Centromeric Subunit (CS) proteins 1-4 (Methods, Fig. 1b,c Fig. 2a, b). AlphaFold2 prediction of the complex between all identified CS proteins resulted in a high-confidence model that globally fitted the experimental cryoEM map (Extended Data Fig. 4b, c). Modeling these sequences into the experimental cryoEM map resulted in a good side-chain fit for all CS 1-4 proteins (Extended Data Fig. 4d).

**Fig. 2.**
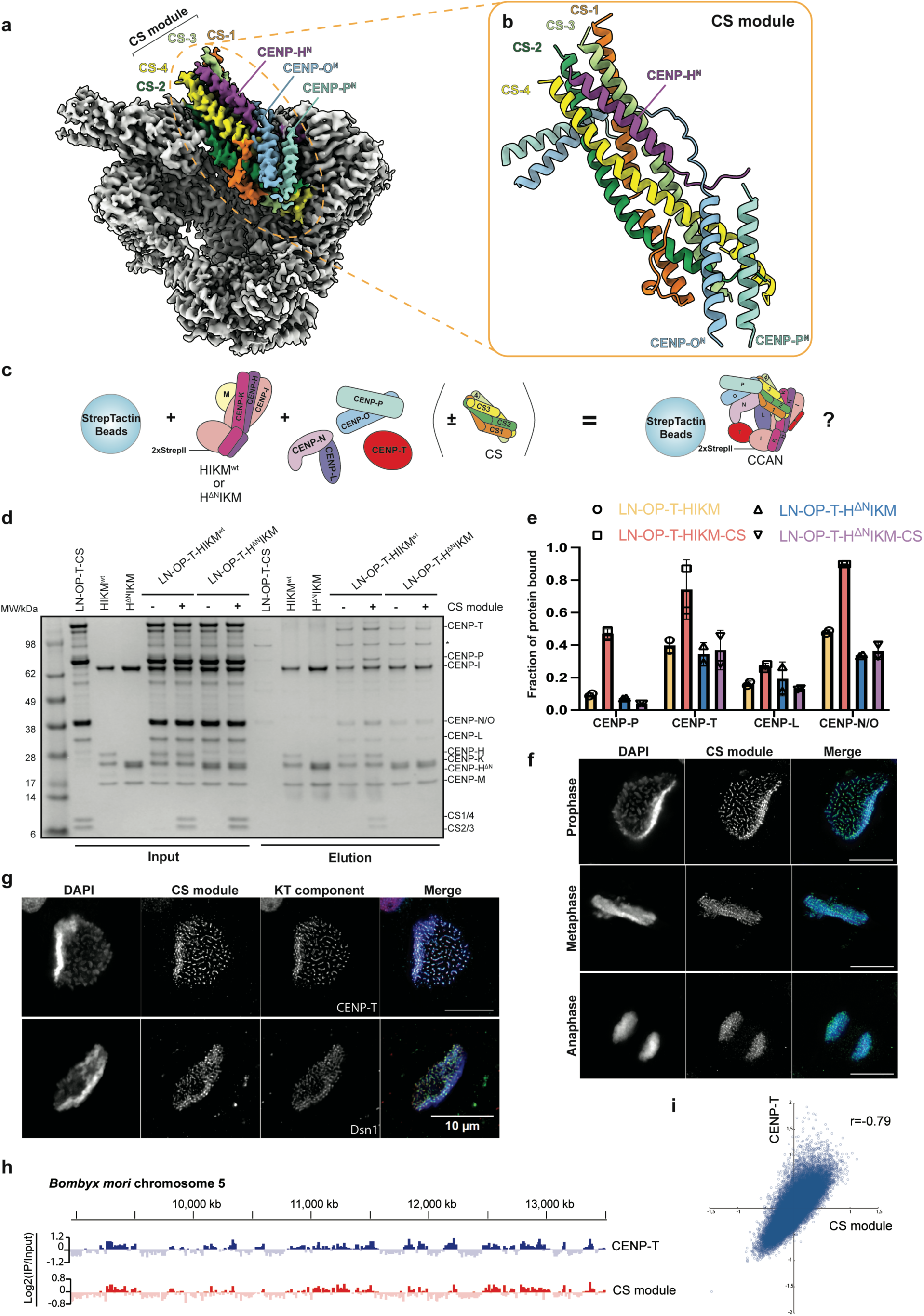
The CS module is an integral component of the bmCCAN. **a,** CryoEM reconstruction of the complete apo-bmCCAN with the CS proteins colored by subunit. The N-termini of bmCENP-H, bmCENP-O, and bmCENP-P proteins that directly support CS module binding are also colored and annotated. **b,** A molecular model of the CS module, with supporting structures from the bmCENP-H, bmCENP-O, and bmCENP-P also shown. **c,** A schematic of the bmCCAN pull-down assay. BmCENP-HIKM wild-type or bmCENP-H^Δ^IKM mutant (where 1-33 amino acids in bmCENP-H protein are deleted) were immobilized on the StrepTactin resin via 2xStrep-tag II at the N-terminus of bmCENP-I protein. Recombinantly purified bmCENP-LN, bmCENP-OP, bmCENP-T and the CS module were added in different combinations to assess bmCCAN assembly. **d,** A representative result of the bmCCAN StrepTactin resin pull-down assay. The components added to each reaction are shown above the gel. CS module addition is indicated with + and -, with no CS module added to other reactions. * indicates a co-purifying contaminant observed during bmCENP-T purification. The assay was repeated in technical duplicates. **e,** Quantification of the bmCCAN StrepTactin resin pull-down assay. Band intensity for CENP-P, CENP-T, CENP-L and overlapping CENP-N/O protein bands was measured in the input and elution reactions in raw gel images shown in (d) to derive the fraction of the protein bound. The data point for each repeat is shown. Column height represents mean value and error bars represent standard deviation. **f,** DAPI-stained mitotic chromosomes (blue) of different stages, immunostained for the CS module (cyan). Scale bar: 5 μm. **g,** DAPI-stained mitotic chromosomes (blue), immunostained for the CS module (red) and either bmCENP-T (cyan, top) or bmDsn1 (cyan, bottom). Scale bar: 10μm. **h,** A representative portion of chromosome 5 showing ChIP-seq signal for bmCENP-T and the CS module. ChIP-seq signals are represented as average log2 ratios of IP/Input in genome-wide 10 kb windows. **i,** Genome-wide correlation plot of CS and BmCENP-T occupancy. The correlation coefficient (r) is indicated.

The newly discovered CS 1-4 subunits, together with bmCENP-H^N^, form the central pentameric coiled-coil that binds along bmCENP-OP^RWD^. The N-terminal extensions of bmCENP-O (bmCENP-O^N^) and bmCENP-P (bmCENP-P^N^) wrap around the CS module, securing its position next to the bmCENP-N and bmCENP-M subunits (Fig. 1, Fig. 2a, b). In the fully assembled structure, bmCENP-H^N^ threads beneath the bmCENP-O^N^ and bmCENP-P^N^ extensions (Fig. 2b). Additionally, CS-4 inserts its C-terminus at the pocket formed by the bmCENP-IK interface (Extended Data Fig. 4e). Notably, CS-4:bmCENP-IK interaction is identical to the human hsCENP-O:hsCENP-IK interaction (Extended Data Fig. 4e)^19–21^. At key contact sites, CS-4 and hsCENP-O share chemically similar amino acids, suggesting that these unrelated proteins accomplish a conserved function.

### The CS module completes and augments bmCCAN architecture

We used the *S. frugiperda* CS module sequences to identify, clone and reconstitute *B. mori* CS 1-4 orthologs. CS 1-4 proteins formed a single soluble complex in solution with 1:1:1:1 stoichiometry as determined by size-exclusion chromatography (SEC) coupled with multi-angle light scattering (MALS) (Extended Data Fig. 5a,b). Using an on-column reconstitution approach, we observed that the CS module stably associates only with the bmCENP-HIKM module (Extended Data Fig. 5c-f). We reasoned that this is likely because bmCENP-H^N^ contributes an α-helix to the pentameric coiled-coil structure (Fig. 2a,b). Consistent with this hypothesis, the interaction between the CS module and the bmCENP-HIKM module was abolished when bmCENP-H^N^ was deleted (bmCENP-H^ΔN^IKM) (Extended Data Fig. 6a). Based on our observation that bmCENP-O^N^ and bmCENP-P^N^ extensions wrap around the CS module and that bmCENP-OP module was sub-stoichiometric in our initial bmCCAN purification (Extended Data Fig. 1d), we reasoned that the CS module might be an integral component of the bmCCAN, which helps to link and stabilize the bmCCAN structure, and in particular important in recruiting the bmCENP-OP. Consistent with our hypothesis, all distinct bmCCAN components were enriched on bmCENP-HIKM-immobilized beads specifically in the presence of the CS module, with the most dramatic enhancement observed for the bmCENP-OP module (Fig. 2 c-e). Deletion of the bmCENP-H^N^ abolished CS module recruitment and CS-mediated bmCCAN stabilization effects, confirming that specific interactions are necessary for correct bmCCAN assembly (Fig. 2d, e). We also observed that the CS module readily became incorporated into bmCCAN during the on-column reconstitutions, and improved the stoichiometry of all other components in the main bmCCAN peak (Extended Data Fig. 6b). Lastly, including the CS module in parallel expression and purification of bmCCAN notably improved the stoichiometry of the bmCENP-OP module (Extended Data Fig. 1, 6c). Collectively, our structural and biochemical analyses suggests that the newly discovered CS module is an important component of the bmCCAN architecture that links multiple modules together and appears to be necessary for bmCENP-OP recruitment.

Unexpectedly, bmCENP-OP associated only weakly with the bmCCAN in the absence of the CS module despite bmCENP-OP sharing a large binding interface with the bmCENP-N protein (Fig. 1a-c). This is different for the human or yeast systems, where CENP-N:CENP-O interactions are mediated primarily by an extension in the CENP-N protein that folds along CENP-O (Extended Data Fig. 7a)^9,26^. The difference is likely due to the presence of CENP-QU in human and yeast that pull CENP-OP away from the CENP-LN module, reducing CENP-N:CENP-O interaction. BmCENP-O is moved 25 Å towards bmCENP-LN and bmCENP-P is rotated by 60° compared to the human hsCENP-O/P (Extended Data Fig. 7b)^19^. A bmCENP-O^ΔN^ mutant, lacking the first 76 amino acids that interact with the CS module (Fig. 2a, b), and a bmCENP-O^CENP-Nm^ mutant (L141A, Q190A, F192A, L195A, E201A substitutions disrupting bmCENP-N interaction, Extended Data Fig. 7c) both failed to associate with bmCCAN in pull-down assays, even when the CS module was present. This suggests multiple interactions are required for complete bmCENP-OP association with the bmCCAN (Extended Data Fig. 7d, e). This is similar to the hsCCAN assembly principles in the human system, where disrupting individual subunit interfaces was sufficient to abolish the formation of the entire hsCCAN^19,34^.

### The CS module is a *bona fide* inner kinetochore component in cells

To understand if the CS module is a kinetochore component in cells, we performed immunoprecipitation (IP) in Sf9 cells experiments targeting CS-1, CS-2, CENP-O, and CENP-P to profile their interactions by MS. MS revealed that CS-1/2 proteins could pull down all other known components (Extended Data 1). We could also detect CS proteins in reciprocal CENP-O/P pull-down experiments (Extended Data 1). Additionally, we used MS to analyze over-expressed and Heparin-purified bmCCAN (Extended Data Fig. 1d, Extended Data 2). Using these and previously published results, we identified other proteins that are also found at human centromeres such as FACT, topoisomerases, cohesin and condensin complexes to be enriched at the bmCCAN ^8,29,31,35–37^. This suggests that despite the holocentric architecture and the absence of CENP-A and CENP-C, similar factors likely govern centromere function across evolutionarily distant organisms.

We targeted the CS module in immunofluorescence (IF) microscopy experiments using a custom-made antibody against the complex. The CS module localized to mitotic chromosomes through all stages of mitosis, similar to other known bmCCAN components (Fig. 2f) ^29^. Furthermore, the CS module also co-localized with bmCENP-T and bmDsn1 proteins, established markers of inner and outer kinetochore localization in Lepidoptera, further supporting its integration into functional kinetochore complexes (Fig. 2g).

To assign CS module localization at the chromatin level we performed chromatin immunoprecipitation and sequencing (ChIP-Seq) targeting the CS module in unsynchronized *B. mori* cells (Fig. 2h, i). We observed that the CS module profile correlated strongly with previously established bmCENP-T profiles, with CS assembling over large domains found along the entire length of the *B. mori* chromosome^31^. The similarity between both CS and bmCENP-T ChIP-seq profiles is consistent with these two modules forming part of the same complex on chromatin in cells.

We took advantage of our specific antibodies to gain insights into localization dependencies between different modules of the *B. mori* CCAN. We found that CS module chromatin localization in cells was dependent on bmCENP-T, bmCENP-I, and bmCENP-H, components essential for bmCCAN assembly in cells (Fig. 3a, b)^29^. In contrast, depletion of the CS module did not affect bmCENP-T localization, in line with the CS module being recruited downstream of bmCENP-T (Extended Data Fig. 8a, b). Depletion of the bmCENP-OP complex had minor effects on CS module levels on chromatin, while depletion of CS abolished bmCENP-OP complex recruitment (Fig. 3c, d). These data are consistent with the CS module’s role in recruiting bmCENP-OP but not *vice versa*. Likewise, depletion of the bmCENP-OP complex did not affect chromatin levels of bmCENP-T, while depleting bmCENP-K significantly reduced bmCENP-T localization to chromosomes, consistent with previous localization dependencies between bmCENP-T and the bmCENP-HIKM complex (Extended Data Fig. 8a, b)^29^. Together with previously published analysis of bmCENP-LN module localization dependencies, our work suggests that bmCENP-LN, bmCENP-HIKM, and bmCENP-T are interdependent and required for minimal bmCCAN assembly on chromatin. BmCENP-LN-HIKM-T then recruits the CS module, likely via bmCENP-H^N^, and the CS module in turn recruits bmCENP-OP (Fig. 3e).

**Fig. 3.**
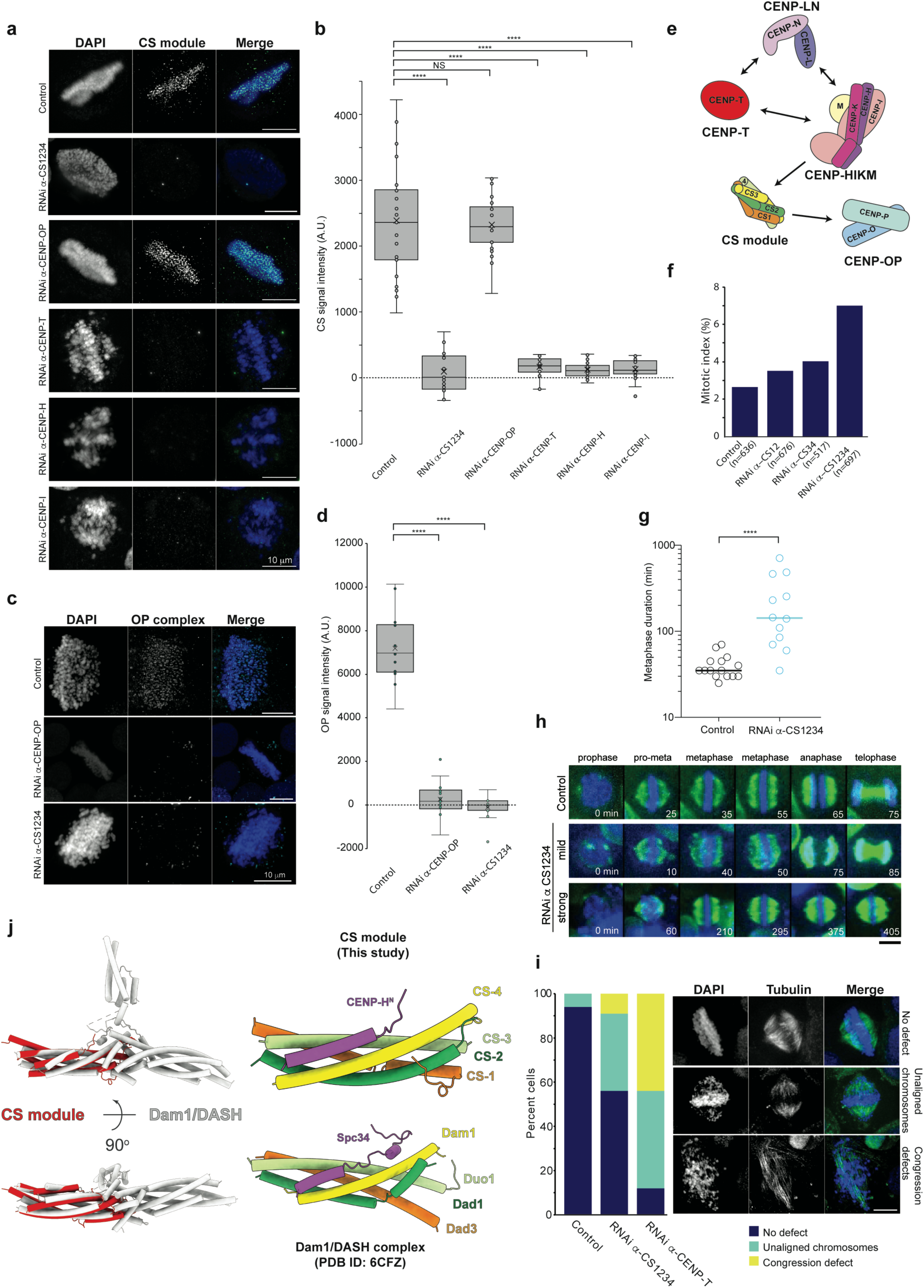
Depletion of CS results in mitotic defects and abolished BmCENP-OP recruitment. **a,** Representative images of mitotic *B. mori* cells showing the levels of the endogenous CS module with and without depletion of the CS module, bmCENP-OP, bmCENP-H, bmCENP-T, and bmCENP-I kinetochore components. Scale bar: 10 µm. **b,** Quantification of mean fluorescence intensity for the CS module signals in the control sample or upon kinetochore depletions. Statistical significance was tested using a two-tailed students’s t-test with unequal variance (p ≤ 0.0001 for four stars, p ≤ 0.001 for three stars, p ≤ 0.01 for two stars and p ≤ 0.05 for one star). **c,** Representative images of mitotic *B. mori* cells showing the levels of endogenous bmCENP-OP with and without depletion of bmCENP-OP and the CS module. Scale bar: 10 µm. **d,** Quantification of mean fluorescence intensity for bmCENP-OP signals in the control or upon bmCENP-OP or the CS module depletions. Statistical significance was tested using a two-tailed students’s t-test with unequal variance (p ≤ 0.0001 for four stars, p ≤ 0.001 for three stars, p ≤ 0.01 for two stars and p ≤ 0.05 for one star). **e,** The assembly hierarchy of the apo-bmCCAN complex based on depletion results determined in this study and previous results ^29^. **f,** Graph showing the percentage of mitotic cells (H3S10ph positive cells) seven days in control or after RNAi-mediated depletions of different CS components (n= number of cells). **g,** Log-scale plot of metaphase duration of control and CS-depleted cells. From available time-lapsed movies, metaphase duration was defined as the time from pro-metaphase (bipolar spindle formation and chromosome aggregation at the mid-spindle) through to telophase (complete chromosome segregation and start of cytokinesis). *P*-value was calculated using a non-parametric *t*-test (p≤ 0.0001 for four stars). **h,** Max-projection time-lapse images of the spindle (green) and chromosome (blue) dynamics for control and CS-depleted cells from mitotic stages prophase through telophase. Time is indicated in minutes. Scale bar, 10 µm. For control, note the symmetrical “square” shaped spindle and strong chromosome compaction and alignment at the metaphase plate. For the CS-depletion, note the spectrum (weak and strong) of mal-shaped spindles and chromosome misalignment during metaphase. Strong phenotypes include long delay in mitosis duration and lagging chromosomes at anaphase. Spurious chromosome puncta not inside the spindle proper are staining artifacts. **i,** Percentage of cells showing no defects (blue), unaligned chromosomes (cyan) or congression defects (yellow) for control, CS or BmCENP-T depletion experiments. Representative images of mitotic cells stained with anti-tubulin (green) used to classify mitotic defects observed. Scale bar: 10 µm. **j,** Structural comparison between the Dam1/DASH complex ^38^ and the CS module in complex with BmCENP-H.

RNAi-mediated depletion of the CS module in cells resulted in an increased number of cells delayed or arrested in mitosis (Fig. 3f), although to a lesser extent than the depletion of other essential CCAN components, such as bmCENP-T^29^. To understand the cause of the mitotic arrest, we performed live cell imaging on cells depleted of the CS module (Fig. 3g, h and Extended Data Fig. 8c). We observed that cells arrested specifically in metaphase, a defect consistent with impairment of kinetochore function. In addition, quantification of the defects upon CS depletion revealed an increased number of mitotic cells with unaligned chromosomes and congression defects (Fig. 3i). Although the observed phenotypes were less severe than the depletion of bmCENP-T, which, in contrast to the depletion of CS (Extended Data Fig. 8d, e), completely abolishes the outer kinetochore recruitment^29^, our results show that the CS module plays an important role in high-fidelity chromosome segregation and correct cell cycle progression.

### The CS module is evolutionarily related to the Dam1/DASH complex

We used sequence alignment tools to understand the evolutionary origin and conservation of the CS module. These analyses revealed a distant homology between three CS proteins (CS-2, CS-3, and CS-4) and the subunits of the Dam1/DASH complex, an established component of the outer kinetochore in certain yeast species (Fig. 3j, Extended Data Fig. 9). The structural comparison confirmed a close similarity between the CS module and one-half of Dam1/DASH (Fig. 3j)^38,39^. Specifically, we found that CS-1 protein corresponds to Dad3, CS-2 protein to Dad1, CS-3 protein to Duo1, CS-4 protein to Dam1 in the structure. The assignment is consistent with phylogenetic clustering of the CS-2, CS-3 and CS-4 with Dad1, Duo1 and Dam1 clades rather than their paralogs described previously^40^. In addition, the N-terminus of bmCENP-H structurally corresponds to Spc34, a component of the Dam1/DASH complex involved in Ndc80 binding^39^ but sequence alignment does not detect any similarities.

The Dam1/DASH complex is a decamer that further oligomerizes to form a ring-like structure around microtubules^38,39^. The proper function of the Dam1/DASH complex is dependent on oligomerization and its association with the Ndc80 complex and microtubules. To mediate these processes, parts of the Dam1/DASH complex, including the C-termini of Dam1, Duo1, Spc19 and Spc34 which collectively interact with the Ndc80 complex and microtubules, are target sites of the error correction pathway that ensures proper kinetochore-microtubule attachment^41^. Alignment of the CS protein sequences to the corresponding Dam1/DASH homologs shows that the homology is restricted to the N-terminal helices that form the core of the Dam1/DASH complex, and none of the aforementioned functionally relevant sites are present in CS module components (Extended Data Fig. 10). Additionally, the CS module in the bmCCAN structure is capped at the bottom by the rest of the bmCCAN, blocking further oligomerization that is seen in the Dam1/DASH complex. SEC-MALS data further show that the CS module is monomeric in solution (Fig. 1, Extended Data Fig. 5a, b), which is consistent with our IP-MS analyses in Lepidopteran cell lines targeting CS-1 and CS-2 proteins that did not reveal additional putative homologs of the Dam1/DASH complex (Extended Data 1).

### Head-to-head bmCCAN dimer bends the DNA into a loop

To understand how bmCCAN interacts with DNA, we determined the bmCCAN structure bound to linear double-stranded DNA (Fig. 4, Extended Data Fig. 11, 12, Extended Table 1, Supplementary Movie 1). We used the 147 base-pair (bp) Widom 601 DNA sequence for our structural work, which has an AT percentage comparable to the *B. mori* genome^42^. We obtained several reconstructions from this sample: apo-bmCCAN structure similar to the one described above; 2.8 Å reconstruction of monomeric bmCCAN bound to ∼30 bp of DNA (66% of bmCCAN particles, Fig. 4a-c); and 3.9-4.5 Å multi-body reconstructions of a head-to-head bmCCAN dimer bound to ∼55 bp of DNA (19.2% of CCAN particles, Fig. 4d, e).

**Fig. 4.**
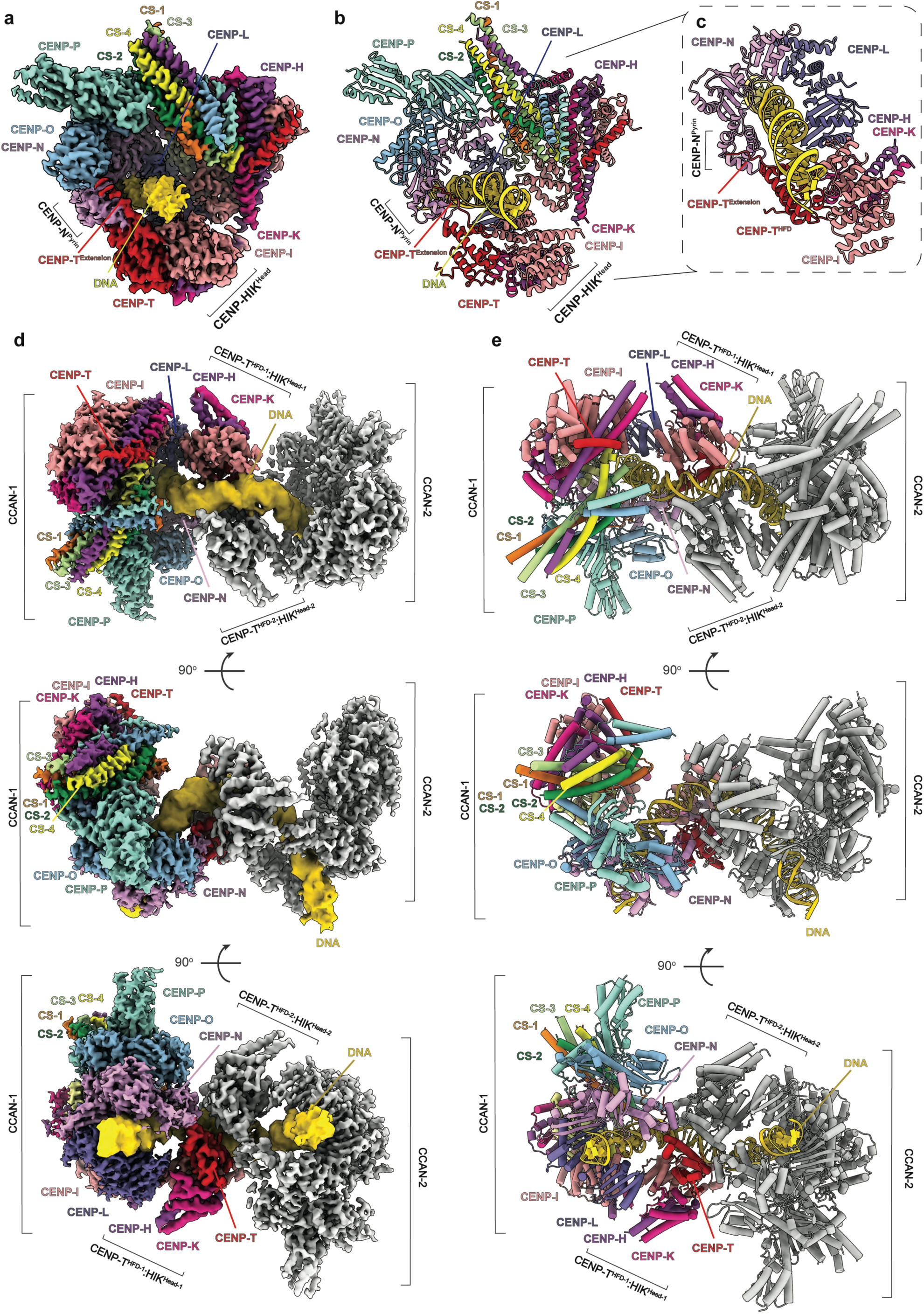
Structure of the bmCCAN bound to DNA. **a,** Composite cryoEM reconstruction of the monomeric bmCCAN bound to 35 bp of double-stranded DNA. The composite map corresponds to the combined locally refined bmCENP-T-HIK^Head^ map and the rest of the bmCCAN:DNA map (Extended Data Fig. 12). **b,** Molecular model of the monomeric bmCCAN:DNA complex structure. **c,** A close-up view of the bmCENP-LN-HIK-T interfaces that line the DNA binding channel in the monomeric bmCCAN:DNA structure. **d,** Composite cryoEM reconstruction of the dimeric bmCCAN bound to approximately 50 bp of double-stranded DNA. A composite map is generated from multibody refinement, dividing the complex into three rigid bodies: CCAN-1, CENP-T^HFD^:HIK^Head^ dimerization interface, and CCAN-2 multibody maps (Extended Data Fig. 12). CCAN-1 is colored by subunit while CCAN-2 is colored in light grey. **e,** Molecular model of the dimeric bmCCAN:DNA complex.

In the monomeric bmCCAN-DNA structure, the DNA is fully topologically entrapped by bmCENP-LN:T:HIK^Head^ modules that form a closed chamber via significant protein-protein interactions to augment the central bmCENP-LN DNA-binding channel (Fig. 4c). The DNA in this state bends slightly at the bmCENP-T^HFD^:HIK^Head^ interface but generally assumes a straight conformation. Overall, the DNA-binding mode via the CENP-LN channel is similar to the ones reported for hsCCAN and scCCAN^9,10,19,21^. However, in the absence of CENP-QU proteins, the topological DNA-binding chamber is closed directly by prominent bmCENP-N:T interactions involving both bmCENP-T^HFD^ and bmCENP-T N-terminal extension (bmCENP-T^Extension^) that interacts with the bmCENP-N^Pyrin^ domain surface (Fig. 4a-c).

In the dimeric state, two CCANs associate in a head-to-head manner (Fig. 4d, e). The unique bmCCAN dimer conformation is achieved by dimerization between bmCENP-N^Pyrin^:T^HFD^:HIK^Head^ groups of bmCCAN-1 and bmCCAN-2 that form a composite interface to bind the DNA duplex, with DNA itself further linking and stabilizing the dimer state. The DNA is first fully topologically entrapped by bmCENP-LN:T:HIK^Head^ modules of bmCCAN-1, as described for the monomeric CCAN:DNA complex (Fig. 5a,b). The DNA is then wrapped around a deep electropositive groove formed by the dimerized bmCENP-N^Pyrin^:T^HFD^:HIK^Head^ modules before passing through a pseudo-symmetrically positioned bmCENP-LN channel in bmCCAN-2 (Fig. 5a, b). Overall, ∼50 bp of DNA are directly bound by the bmCCAN dimer, and the dimer formation induces an approximately 120° bend in the DNA (Fig. 4c,d and 5c). BmCENP-T^HFD^, having an altered HFD, forms an atypical dimer along the outer surface of the central helix of its HFD (Fig. 5d, e). Interestingly, we observe a distinct density for a metal ion at the bmCENP-T:CENP-T dimer interface tetravalently coordinated by H938 and E934 residues from both bmCCANs (Fig. 5e). The identity of the metal is unknown but the coordination geometry and distances are consistent with Zn^2+^ or other tetravalent metal ions. The DNA bending is likely achieved by the dimer formation itself, considering that DNA assumes a largely straight trajectory in the monomeric bmCCAN-DNA complex (Supplementary Movie 1). Overall, the bmCCAN dimer forms via non-canonical interfaces to tightly grip and bend the DNA into a loop-like structure.

**Fig. 5.**
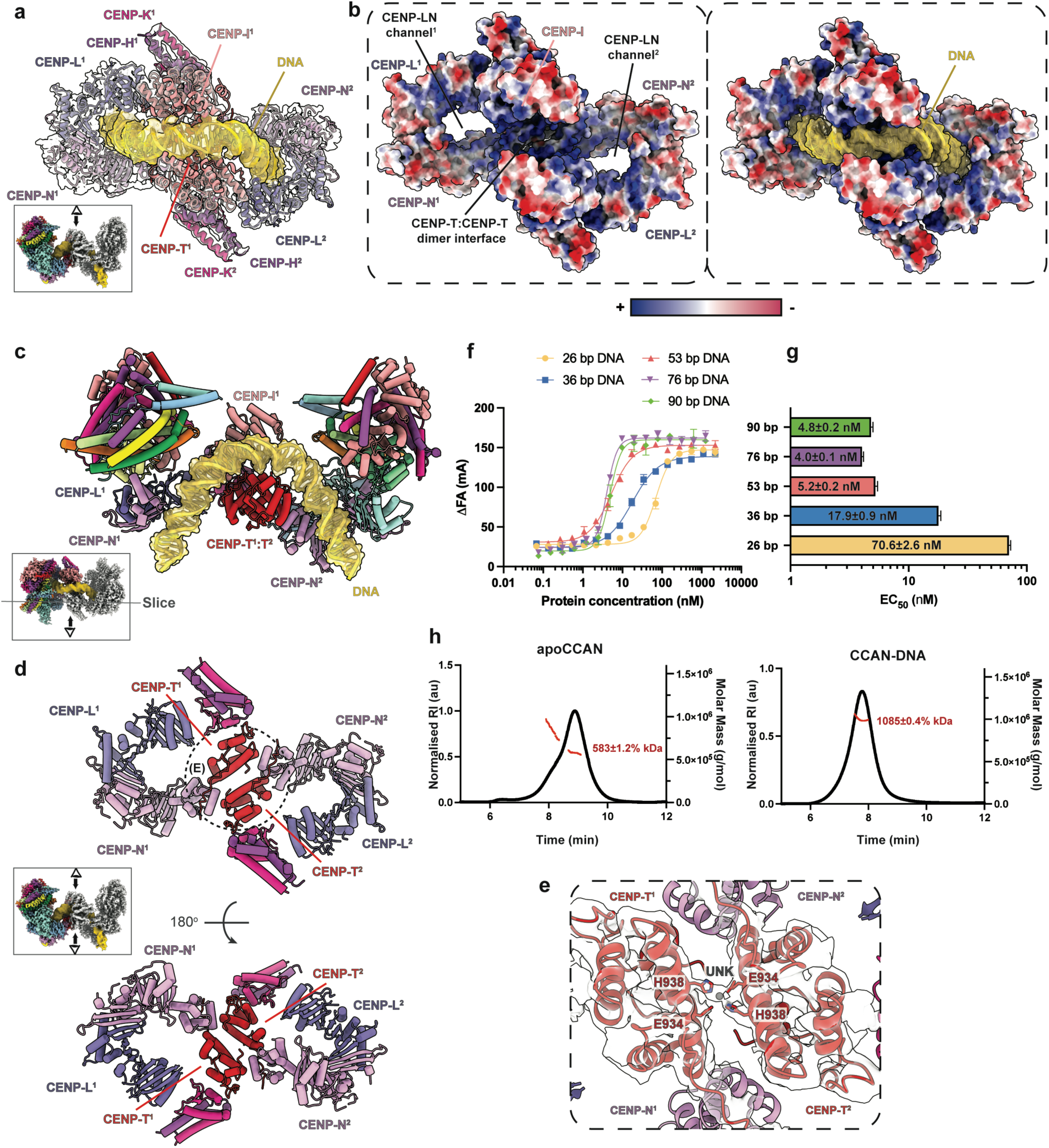
BmCCAN dimer bound to DNA is a self-contained functional unit of the *Bombyx mori* inner kinetochore. **a,** Molecular model of the bmCCAN dimerization interface fit into the cryoEM reconstruction with inset indicating the viewing direction relative to the bmCCAN-DNA dimer complex. The superscript numbering indicates to which CCAN protomer the indicated subunit belongs. **b,** BmCCAN dimerization interface in surface representation colored by electrostatic potential, with blue showing positive electrostatic potential and red showing negative electrostatic potential. The DNA is added in the right panel in transparent surface and cartoon representations to highlight shape and charge complementarity. **c,** A slice through the bmCCAN-DNA dimer molecular model showing 120° bending in the DNA induced by bmCCAN dimerization. The inset indicates the location of the slice and the viewing direction. The DNA is shown in surface and cartoon representations. **d,** A molecular model of the bmCCAN dimerization interface with the DNA model hidden, highlighting CENP-T^1^:CENP-T^2^ and CENP-T^1^:CENP-N^2^ interfaces. **e,** Molecular model of the CENP-T dimerization interface fit into the transparent cryoEM density map with unknown (UNK) metal ion placed into the density map. The H938 distances to the UNK metal ion are between 1.8-2.3 Å. The E934 distances to the UNK metal ion are between 1.9-2.7 Å. **f,** Results of the fluorescence polarization (FP) assay. BmCCAN was added in increasing amounts to FAM-labeled double-stranded DNA of indicated lengths. The difference in fluorescence anisotropy (ΔFA) is plotted against bmCCAN protein concentration. The assay was repeated at least in technical triplicates with mean value and standard deviation values shown. The best-fit regression analysis curve is fitted through the data points to calculate half-maximal effective concentration (EC_50_) values. **g,** EC_50_ values calculated from FP assay shown in **(f)** are plotted with bar height indicating the mean value and the error bars indicating the standard error of the mean (SEM). EC_50_ with SEM values for each DNA length are also stated in the figure. **h,** SEC-MALS results of the apo-bmCCAN and CCAN-DNA complex plotted with normalized refractive index (RI) and molecular mass on Y-axes against elution time of complexes on the X-axis. The estimated molecular weights are: apo-bmCCAN is 530,817 Da; DNA is 90,716 Da; monomeric CCAN-DNA complex is 621,533 Da; dimeric CCAN-DNA complex is 1,152,350 Da. The measured molecular weight for apo-bmCCAN is 583.0 ± 1.175% kDa with 1.1 ± 1.7% polydispersity (Mw/Mn) value. The measured molecular weight for the bmCCAN-DNA complex is 1085 ± 0.4% kDa with 1.0 ± 0.6% polydispersity (Mw/Mn) value. The analysis was repeated in technical duplicates.

### Dimeric bmCCAN is the functional unit of *Bombyx mori* inner kinetochore

To correlate our structural observations with the behavior of the bmCCAN in solution, we measured the affinity between bmCCAN and DNA constructs of different lengths using a fluorescence polarization (FP) assay (Fig. 5f, g). We observed that the affinity of bmCCAN for DNA increased as DNA length increased from 26 bp to 53 bp, with affinity of bmCCAN for DNA of 53 bp, 76 bp, and 90 bp being very similar. This is consistent with our structural observations where ∼50bp of DNA are engaged by bmCCAN directly (Fig. 5c). We also observed that mutating either bmCENP-LN channel or bmCENP-T residues involved in DNA binding (in both monomeric and dimeric states) did not affect bmCCAN assembly but significantly reduced bmCCAN binding to the DNA, consistent with the structure (Extended Data Fig. 13a-f).

We also evaluated the role of bmCCAN assembly intermediates in DNA binding. We could detect weak DNA binding by the bmCENP-HIKM:T modules together but not by bmCENP-LN:T or bmCENP-HIKM:LN modules (Extended Data Fig. 13g, h). Combining the bmCENP-LN:T:HIKM modules resulted in significantly increased binding, consistent with these modules containing all DNA-interacting residues. Interestingly, adding bmCENP-OP or the CS module separately to the bmCENP-LN:T:HIKM assembly only modestly changed the affinity for DNA, but combining all modules into a complete bmCCAN resulted in tight DNA binding. This is consistent with our observations that the CS module is required to recruit bmCENP-OP to bmCCAN and that the bmCENP-OP module further stabilizes the bmCENP-LN structurally into a closed DNA-binding channel conformation (Fig. 1a).

In the dimeric bmCCAN-DNA structure, DNA appears to act as a glue, threading through two bmCCANs and linking them in a head-to-head configuration (Fig. 4d, e). We used SEC-MALS to measure bmCCAN molecular weight in the absence (apo-bmCCAN) or the presence of 147 bp of DNA. We observed that the addition of DNA converted bmCCAN from a monomeric state to an exclusively dimeric state, with no higher oligomeric assemblies observed (Fig. 5h). This is consistent with cryoEM observations where monomeric apo-bmCCAN was only observed in the absence of DNA (Fig. 1) and the dimeric state was detected when DNA was present (Fig. 4d, e).

We designed mutations at the bmCCAN dimerization interface in bmCENP-T protein to understand the functional relevance of the bmCCAN dimer (Extended Data Fig. 14a, b). We observed that these mutations reduced bmCCAN affinity for DNA *in vitro*, suggesting that dimerization strengthens bmCCAN interaction with DNA (Extended Data Fig. 14c, d).

## Discussion

Our characterization of the *B. mori* kinetochore reveals conserved and novel principles of kinetochore organization. Our structural analysis shows how the evolutionarily divergent bmCCAN topologically entraps DNA using the CENP-LN channel, which emerges as the conserved cornerstone feature of all CCAN complexes described so far irrespective of centromere type (Fig. 1d). We identify previously uncharacterized CS1-4 proteins and show that they are evolutionarily related to the Dam1/DASH complex, an outer kinetochore component that stabilizes and regulates spindle microtubule attachments in various fungal species^39,43,44^. The presence of the CS module as part of the *B. mori* CCAN suggests a functional repurposing of these proteins in stabilizing inner kinetochore formation instead of contributing to outer kinetochore function. This new function might have become particularly relevant in *B. mori* and other Lepidoptera to compensate for the absence of CENP-C and/or CENP-QU proteins, which play similar roles in other eukaryotes^18,45,46^. In addition, the presence of distant homologs of Dam1/DASH complex components in other eukaryotes raises the possibility that these homologs may also have been repurposed to function at the inner kinetochore rather than their well-established role at the outer kinetochore^40^. Our findings also suggest that the lack of CENP-C and/or CENP-QU in some eukaryotes, but the presence of most other CCAN genes, could indicate that the CS module or other functionally similar components might be present at the inner kinetochores in these species^47^.

Beyond the innovations at the inner kinetochore, our study uncovers a striking structural feature of the *B. mori* kinetochore-DNA complex. BmCCAN bound to DNA assembles into a self-contained dimer, arranged in a head-to-head configuration that induces a 120° bend in the DNA. Intriguingly, the bmCCAN dimer is globally reminiscent of the complete inner kinetochore structure of the *S. cerevisiae* point-centromere (Extended Data Fig. 15a, b). At the point centromere of *S. cerevisiae*, two scCCAN dimers bind the unwrapped DNA ends of scCENP-A^Nuc^ in a head-to-head arrangement from either side to form a complete kinetochore together with a few auxiliary and Saccharomycotina-specific factors ^10^. Though the scCCAN is asymmetric, contains different components, and assembles via completely different interfaces, our results suggest that the bmCCAN dimer effectively forms an alternative point-centromere-like architecture. This architecture might have evolved as a cause or a consequence of the loss of CENP-A. Instead of being mediated via the CENP-A^Nuc^, the atypical interaction along the two CENP-T^HFD^ brings two bmCCAN complexes together. In addition, bmCCAN also wraps DNA across the dimerization interface, akin to canonical nucleosomes and the human histone-fold CENP-TWSX module. The bmCCAN dimer is an intrinsically self-contained unit incompatible with further oligomerization unless the DNA trajectory is drastically altered. Supporting this, our SEC-MALS and reconstitution experiments consistently failed to detect larger bmCCAN oligomers, even when longer DNA substrates or excess bmCCAN were provided. These findings suggest that, in Lepidoptera, holocentric kinetochore assembly is likely achieved by the foundational “unit module” bmCCAN dimers, strikingly related to the point-kinetochore of *S. cerevisiae*, along chromatin at discrete genomic loci (Fig. 6a-d). This organization may extend to mitosis, where rather than forming continuous higher-order structures, centromeres in Lepidoptera consist of multiple, point-centromere-like kinetochore units that individually recapitulate some or all fundamental functions of kinetochores, such as robust microtubule attachment, error-correction pathway, and biorientation. However, we cannot exclude other conformations of the bmCCAN that could be driven by mitotic conditions and factors associated with it, nor can we exclude additional proteins that could link multiple bmCCAN dimers together.

**Fig. 6.**
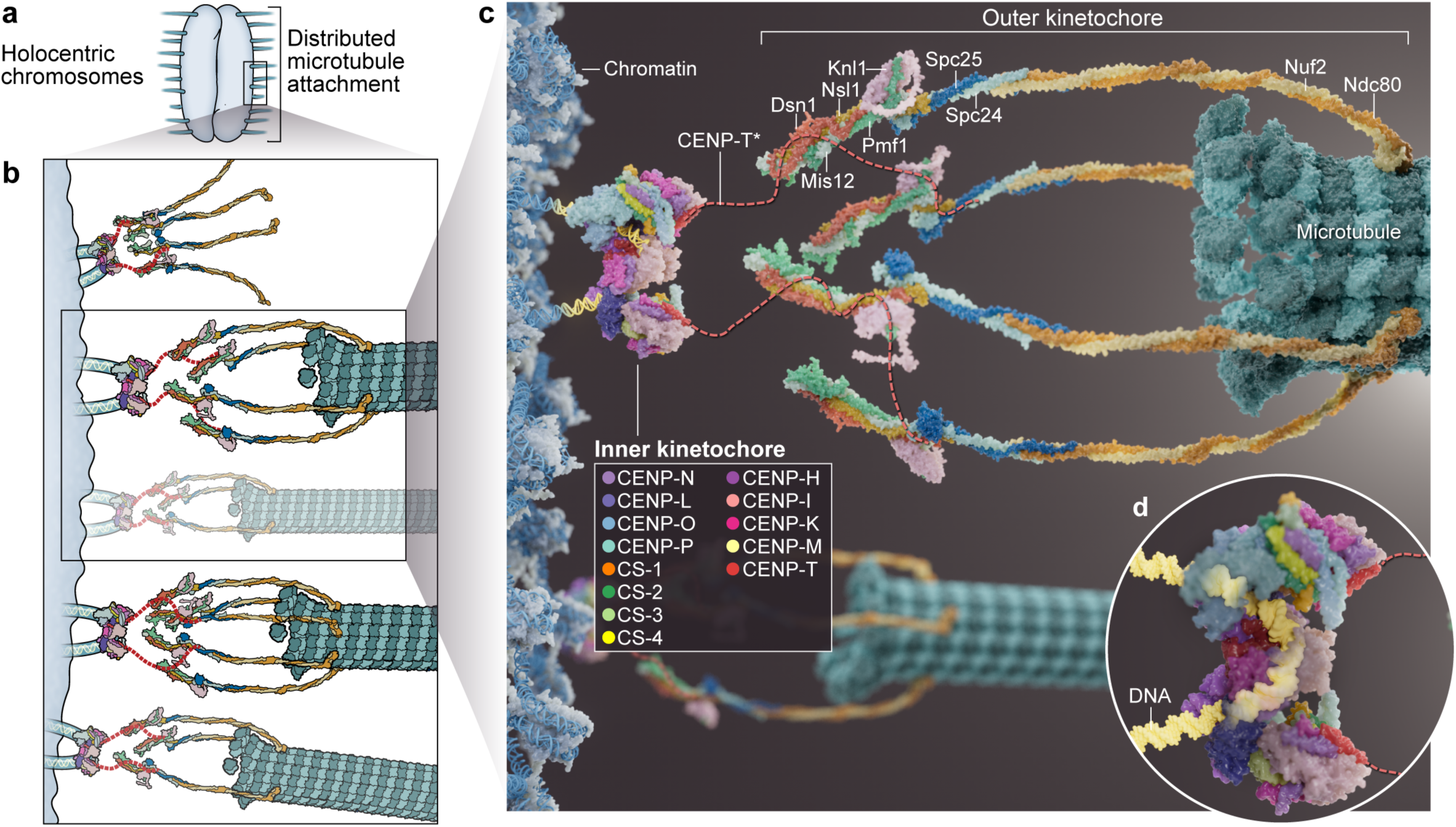
Molecular model of the *Bombyx mori* holocentric kinetochore architecture. **a,** A cartoon illustration of the holocentric compacted chromosomes of *Bombyx mori* during mitosis. **b,** BmCCAN assembles as functional dimers at kinetochore-permissive chromosome loci, recruiting the outer kinetochore KMN network via disordered CENP-T N-terminal tails shown in dotted red lines. The KMN network might have variable stoichiometry numbers, and not all kinetochores might be microtubule bound. **c,** Molecular model of the complete *Bombyx mori* holokinetochore bound to the microtubule (Methods). The experimentally determined bmCCAN dimer bound to DNA is attached via its two CENP-T N-terminal tails (CENP-T*) to four copies of the KMN network predicted using AlphaFold2. The stoichiometry of the KMN network and its interaction with CENP-T are not experimentally validated and represent a model. **d,** A surface representation of the bmCCAN dimer structure bound to the extended DNA loop.

Our molecular model is consistent with a historic model proposed by Franz Schrader^48^, which describes holocentromeres as polycentromeres of multiple discrete centromeric units dispersed along the chromosomes. In *B. mori*, these units are randomly positioned within large chromosomal regions permissive for kinetochore assembly ^31^, an arrangement similar to centromeric domains described in *Caenorhabditis elegans*, another holocentric species^49–51^. In the context of monocentric chromosome organizations, our model is consistent with suggestions that the “unit module” of hsCCAN is a foundational repeating unit at the human regional centromeres^18–22,52^ and agrees with *in situ* cryoET observations that 20-25 nm nucleosome-associated complexes are randomly distributed at the human centromere^53^. Nevertheless, we hypothesize that larger hsCCAN complexes also exist. Given that both bmCCAN and scCCAN form dimers, and that dimerization of hsCENP-C—a key hsCCAN component—is essential for complete human inner kinetochore assembly^22^, it is plausible that the foundational unit of the human kinetochore is also a dimer. Among available high-resolution CCAN structures, hsCCAN:hsCENP-A^Nuc^ stands out as the only monomeric structure (Extended Data Fig. 15c). Structural modeling of an hsCCAN dimer based on the bmCCAN structure predicts a 22 nm particle with no steric clashes (Extended Data Fig. 15d), suggesting that alternative human CCAN structures are at least possible. Alternatively, hsCCAN may also dimerize across hsCENP-A^Nuc^, similar to scCCAN:scCENP-A^Nuc^ complex.

We also observe that the bmCCAN dimer formation induces a 120° turn in the DNA. We propose that this bent DNA structure improves mechanical resistance against spindle forces, reducing the likelihood of premature chromosome detachment. The DNA has a relatively low persistence length and, therefore, likely bends and buckles when the mitotic spindle applies forces during chromosome segregation (Extended Data Fig. 15e). BmCCAN preforms DNA into a loop-like structure poised for chromosome segregation that would resist buckling and bending when spindle forces are applied (Extended Data Fig. 15a, e). Additionally, DNA motors such as cohesin and condensin are thought to require two proximal DNA strands for their function to extrude loops, and bmCCAN bound to DNA might provide a platform for loading of these SMC complexes^54^. Consistent with this, we detect enriched levels of both cohesin and condensin subunits in our MS analysis of multiple bmCCAN samples. Beyond this study, we speculate that bending DNA into loop-like structures is likely a conserved and fundamental inner kinetochore function. For example, only partial DNA wrapping/bending around hsCENP-TWSX module was observed, but biochemical evidence suggests that hsCENP-TWSX could wrap a longer DNA around itself, generating a DNA arrangement similar to the one we observe in bmCCAN:DNA structure (Extended Data Fig. 15f)^19,55^.

Overall, our results reveal that kinetochore assembly is governed by conserved, modular principles, regardless of centromere type. Our observations also suggest how the fundamental and essential process of chromosome segregation is achieved in organisms with diverse centromeric organizations: the same conserved foundational kinetochore unit, related to the kinetochore architecture of the point-centromere, can be used and repeated as many times as needed to drive chromosome segregation in species with drastically different centromere organizations, including holocentrics.

## Supporting information

Methods_ExtendedDataFigures

## Acknowledgments

We thank Genentech colleagues in the BioMolecular Resources and Structural Biology departments, especially James Kiefer and Peter Hsu for helpful discussions and advice, and all members of the Drinnenberg lab for their support. We also thank Zhen Zeng for the mass spectrometry analysis of the gel bands.

## Author contributions

Conceptualization: IAD and SY

Methodology: IAD, CC and SY

Investigation: CY, SRS, GC, AH, CA, IZ, BB, TKC, CMR, LD, DL, PT, IAD, CC, SY

Visualization: IAD and SY

Funding acquisition: IAD

Project administration: IAD, CC, and SY

Supervision: IAD, CC, and SY

Writing – original draft: IAD, CC and SY

Writing – review and editing CY, SRS, GC, AH, CA, IZ, BB, TKC, CMR, LD, DL, PT, IAD, CC, SY

## Competing interests

CA, IZ, BB, CMR, and CC are Genentech/Roche employees and own shares in the Genentech/Roche group. The other authors declare no competing interests.

## Funding

Labex DEEP ANR-11-LABX-0044 part of the IDEX Idex PSL ANR-10-IDEX-0001-02 PSL (IAD)

Institut Curie ATIP-AVENIR Research grant (IAD)

FRM, Fondation Schlumberger: FSER202202015420 (IAD)

European Research Council CENEVO-758757 (IAD)

Salary support from the French National Centre for Scientific Research, CNRS (IAD)

EMBO LTF 505-2021 (SRS)

## Data availability

All proteomics data have been deposited with either PRIDE (PXD063391) or MassIVE (MSV000097156) repositories. All models have been deposited with Protein Data Bank (PDB ID: 9OKL, 9OKK, 9OKE, 9OKD, 9OKB). All cryoEM maps have been deposited with Electron Microscopy Data Bank (EMDB ID: 70568, 70567, 70561, 70560, 70558).

## Materials & Correspondence

Requests for materials should be addressed to Ines Anna Drinnenberg, Claudio Ciferri, or Stanislau Yatskevich.

## Additional Information

Methods.

Extended Data Figs. 1 to 15.

Extended Data Table 1.

Supplementary Movie 1.

Supplementary Data 1 and 2.

## Notes

### Competing Interest Statement

CY, CA, IZ, BB, TKC, CMR, and CC are Genentech/Roche employees and own shares in the Genentech/Roche group. The other authors declare no competing interests.

## References

1 McAinsh, A. D. & Marston, A. L. The Four Causes: The Functional Architecture of Centromeres and Kinetochores. Annu Rev Genet 56, 279–314 (2022). 10.1146/annurev-genet-072820-034559

2 Mellone, B. G. & Fachinetti, D. Diverse mechanisms of centromere specification. Curr Biol 31, R1491–R1504 (2021). 10.1016/j.cub.2021.09.083

3 Drinnenberg, I. A. & Akiyoshi, B. Evolutionary Lessons from Species with Unique Kinetochores. Prog Mol Subcell Biol 56, 111–138 (2017). 10.1007/978-3-319-58592-5_5

4 Sundararajan, K. & Straight, A. F. Centromere Identity and the Regulation of Chromosome Segregation. Front Cell Dev Biol 10, 914249 (2022). 10.3389/fcell.2022.914249

5 Kixmoeller, K., Allu, P. K. & Black, B. E. The centromere comes into focus: from CENP-A nucleosomes to kinetochore connections with the spindle. Open Biol 10, 200051 (2020). 10.1098/rsob.200051

6 McKinley, K. L., et al. The CENP-L-N Complex Forms a Critical Node in an Integrated Meshwork of Interactions at the Centromere-Kinetochore Interface. Mol Cell 60, 886–898 (2015). 10.1016/j.molcel.2015.10.027

7 Okada, M., et al. The CENP-H-I complex is required for the efficient incorporation of newly synthesized CENP-A into centromeres. Nature cell biology 8, 446–457 (2006). 10.1038/ncb1396

8 Foltz, D. R., et al. The human CENP-A centromeric nucleosome-associated complex. Nature cell biology 8, 458–469 (2006). 10.1038/ncb1397

9 Yan, K., et al. Structure of the inner kinetochore CCAN complex assembled onto a centromeric nucleosome. Nature 574, 278–282 (2019). 10.1038/s41586-019-1609-1

10 Dendooven, T., et al. Cryo-EM structure of the complete inner kinetochore of the budding yeast point centromere. Sci Adv 9, eadg7480 (2023). 10.1126/sciadv.adg7480

11 Hinshaw, S. M. & Harrison, S. C. The structure of the Ctf19c/CCAN from budding yeast. Elife 8 (2019). 10.7554/eLife.44239

12 Henikoff, S. & Henikoff, J. G. “Point” centromeres of Saccharomyces harbor single centromere-specific nucleosomes. Genetics 190, 1575–1577 (2012). 10.1534/genetics.111.137711

13 Akiyoshi, B., et al. Tension directly stabilizes reconstituted kinetochore-microtubule attachments. Nature 468, 576–579 (2010). 10.1038/nature09594

14 Furuyama, S. & Biggins, S. Centromere identity is specified by a single centromeric nucleosome in budding yeast. Proc Natl Acad Sci U S A 104, 14706–14711 (2007). 10.1073/pnas.0706985104

15 Altemose, N., et al. Complete genomic and epigenetic maps of human centromeres. Science 376, eabl4178 (2022). 10.1126/science.abl4178

16 Logsdon, G. A., et al. The variation and evolution of complete human centromeres. Nature 629, 136–145 (2024). 10.1038/s41586-024-07278-3

17 Kato, H., et al. A conserved mechanism for centromeric nucleosome recognition by centromere protein CENP-C. Science 340, 1110–1113 (2013). 10.1126/science.1235532

18 Klare, K., et al. CENP-C is a blueprint for constitutive centromere-associated network assembly within human kinetochores. J Cell Biol 210, 11–22 (2015). 10.1083/jcb.201412028

19 Yatskevich, S., et al. Structure of the human inner kinetochore bound to a centromeric CENP-A nucleosome. Science 376, 844–852 (2022). 10.1126/science.abn3810

20 Pesenti, M. E., et al. Structure of the human inner kinetochore CCAN complex and its significance for human centromere organization. Mol Cell 82, 2113–2131 e2118 (2022). 10.1016/j.molcel.2022.04.027

21 Tian, T., et al. Structural insights into human CCAN complex assembled onto DNA. Cell Discov 8, 90 (2022). 10.1038/s41421-022-00439-6

22 Walstein, K., et al. Assembly principles and stoichiometry of a complete human kinetochore module. Sci Adv 7 (2021). 10.1126/sciadv.abg1037

23 Ali-Ahmad, A., Bilokapic, S., Schafer, I. B., Halic, M. & Sekulic, N. CENP-C unwraps the human CENP-A nucleosome through the H2A C-terminal tail. EMBO Rep 20, e48913 (2019). 10.15252/embr.201948913

24 Carroll, C. W., Milks, K. J. & Straight, A. F. Dual recognition of CENP-A nucleosomes is required for centromere assembly. J Cell Biol 189, 1143–1155 (2010). 10.1083/jcb.201001013

25 Sridhar, S. & Fukagawa, T. Kinetochore Architecture Employs Diverse Linker Strategies Across Evolution. Front Cell Dev Biol 10, 862637 (2022). 10.3389/fcell.2022.862637

26 Yatskevich, S., Barford, D. & Muir, K. W. Conserved and divergent mechanisms of inner kinetochore assembly onto centromeric chromatin. Curr Opin Struct Biol 81, 102638 (2023). 10.1016/j.sbi.2023.102638

27 Tromer, E. C., van Hooff, J. J. E., Kops, G. & Snel, B. Mosaic origin of the eukaryotic kinetochore. Proc Natl Acad Sci U S A 116, 12873–12882 (2019). 10.1073/pnas.1821945116

28 van Hooff, J. J., Tromer, E., van Wijk, L. M., Snel, B. & Kops, G. J. Evolutionary dynamics of the kinetochore network in eukaryotes as revealed by comparative genomics. EMBO Rep 18, 1559–1571 (2017). 10.15252/embr.201744102

29 Cortes-Silva, N., et al. CenH3-Independent Kinetochore Assembly in Lepidoptera Requires CCAN, Including CENP-T. Curr Biol 30, 561–572 e510 (2020). 10.1016/j.cub.2019.12.014

30 Drinnenberg, I. A., deYoung, D., Henikoff, S. & Malik, H. S. Recurrent loss of CenH3 is associated with independent transitions to holocentricity in insects. Elife 3 (2014). 10.7554/eLife.03676

31 Senaratne, A. P., et al. Formation of the CenH3-Deficient Holocentromere in Lepidoptera Avoids Active Chromatin. Curr Biol 31, 173–181 e177 (2021). 10.1016/j.cub.2020.09.078

32 Sankaranarayanan, S. R., et al. Insects evolved a monomeric histone-fold domain in the CENP-T protein family Sundar. bioRxiv 11 (2024). 10.1101/2024.11.15.623767

33 Jamali, K., et al. Automated model building and protein identification in cryo-EM maps. Nature 628, 450–457 (2024). 10.1038/s41586-024-07215-4

34 Pesenti, M. E., et al. Reconstitution of a 26-Subunit Human Kinetochore Reveals Cooperative Microtubule Binding by CENP-OPQUR and NDC80. Mol Cell 71, 923–939 e910 (2018). 10.1016/j.molcel.2018.07.038

35 Izuta, H., et al. Comprehensive analysis of the ICEN (Interphase Centromere Complex) components enriched in the CENP-A chromatin of human cells. Genes Cells 11, 673–684 (2006). 10.1111/j.1365-2443.2006.00969.x

36 Obuse, C., et al. Proteomics analysis of the centromere complex from HeLa interphase cells: UV-damaged DNA binding protein 1 (DDB-1) is a component of the CEN-complex, while BMI-1 is transiently co-localized with the centromeric region in interphase. Genes Cells 9, 105–120 (2004). 10.1111/j.1365-2443.2004.00705.x

37 Schweighofer, J., et al. Interactions with multiple inner kinetochore proteins determine mitotic localization of FACT. J Cell Biol 224 (2025). 10.1083/jcb.202412042

38 Jenni, S. & Harrison, S. C. Structure of the DASH/Dam1 complex shows its role at the yeast kinetochore-microtubule interface. Science 360, 552–558 (2018). 10.1126/science.aar6436

39 Muir, K. W., et al. Structural mechanism of outer kinetochore Dam1-Ndc80 complex assembly on microtubules. Science 382, 1184–1190 (2023). 10.1126/science.adj8736

40 van Rooijen, L. E., Tromer, E. C., van Hooff, J. J. E., Kops, G. & Snel, B. Increased Sampling and Intracomplex Homologies Favor Vertical Over Horizontal Inheritance of the Dam1 Complex. Genome Biol Evol 15 (2023). 10.1093/gbe/evad017

41 Tanaka, T. U. & Zhang, T. SWAP, SWITCH, and STABILIZE: Mechanisms of Kinetochore-Microtubule Error Correction. Cells 11 (2022). 10.3390/cells11091462

42 Kawamoto, M., et al. High-quality genome assembly of the silkworm, Bombyx mori. Insect Biochem Mol Biol 107, 53–62 (2019). 10.1016/j.ibmb.2019.02.002

43 Westermann, S., et al. Formation of a dynamic kinetochore-microtubule interface through assembly of the Dam1 ring complex. Mol Cell 17, 277–290 (2005). 10.1016/j.molcel.2004.12.019

44 Cheeseman, I. M., Enquist-Newman, M., Muller-Reichert, T., Drubin, D. G. & Barnes, G. Mitotic spindle integrity and kinetochore function linked by the Duo1p/Dam1p complex. J Cell Biol 152, 197–212 (2001). 10.1083/jcb.152.1.197

45 Kagawa, N., et al. The CENP-O complex requirement varies among different cell types. Chromosome Res 22, 293–303 (2014). 10.1007/s10577-014-9404-1

46 Hori, T., Okada, M., Maenaka, K. & Fukagawa, T. CENP-O class proteins form a stable complex and are required for proper kinetochore function. Mol Biol Cell 19, 843–854 (2008). 10.1091/mbc.e07-06-0556

47 Shah, H., et al. Life-cycle-coupled evolution of mitosis in close relatives of animals. Nature 630, 116–122 (2024). 10.1038/s41586-024-07430-z

48 Schrader, F. The Role of the Kinetochore in the Chromosomal Evolution of the Heteroptera and Homoptera. Evolution 1, 134–142 (1947). 10.2307/2405489

49 Steiner, F. A. & Henikoff, S. Holocentromeres are dispersed point centromeres localized at transcription factor hotspots. Elife 3, e02025 (2014). 10.7554/eLife.02025

50 Gassmann, R. Cell Division: Chromatin Dynamics Shape Insect Holocentromeres. Curr Biol 31, R34–R37 (2021). 10.1016/j.cub.2020.10.049

51 Gassmann, R., et al. An inverse relationship to germline transcription defines centromeric chromatin in C. elegans. Nature 484, 534–537 (2012). 10.1038/nature10973

52 Weir, J. R., et al. Insights from biochemical reconstitution into the architecture of human kinetochores. Nature 537, 249–253 (2016). 10.1038/nature19333

53 Kixmoeller, K., Tarasovetc, E. V., Mer, E., Chang, Y. W. & Black, B. E. Centromeric chromatin clearings demarcate the site of kinetochore formation. Cell (2025). 10.1016/j.cell.2024.12.025

54 Kim, E., Barth, R. & Dekker, C. Looping the Genome with SMC Complexes. Annu Rev Biochem 92, 15–41 (2023). 10.1146/annurev-biochem-032620-110506

55 Nishino, T., et al. CENP-T-W-S-X forms a unique centromeric chromatin structure with a histone-like fold. Cell 148, 487–501 (2012). 10.1016/j.cell.2011.11.061

