## Supplementary material for "Architecture and Function of Holocentric CENP-A-Independent Kinetochores": Methods_ExtendedDataFigures

**Cloning of the proteins for purification.**

Genes encoding *Bombyx mori* CENP proteins (CENP-H, CENP-I, CENP-K, CENP-M, CENP-O, CENP-P, CENP-L, CENP-N, CENP-T) including mutants and Centromere Subunits (CS) module were synthesized by GenScript with codon optimization for insect cells. Mutants involved in this study are: CENP-O$^{\text{CENP-Nm}}$ (L141A, Q190A, F192A, L195A, E201A), CENP-O$^{\Delta N}$ (M1-N76 deletion), CENP-H$^{\Delta N}$ (M1-T33 deletion), CENP-LN$^{\text{mut}}$ (K281E, K263E, K285E, K197E mutation in CENP-N, K202E, K204E, R195E, K112E mutations in CENP-L), CENP-T$^{\text{mut}}$ (K890E, K892E, K895E, R896E), CENP-T$^{\text{DimerMut}}$ (H938A, E934A, K931A, E960A, K945A, K944A).

Genes for the CENP-HIKM, CENP-OP, CENP-LN, CS module, and their corresponding mutants as indicated in the text and in the figure legends, were subsequently cloned into pAcGP67-Dual expression vector. Tobacco etch virus (TEV) protease cleavable hexahistidine (His8)-tag followed by SNAP-tag was added to the N-terminus of CENP-T protein and CENP-T mutants. TEV protease cleavable double streptag-II (DS) tags were added to the C-termini of CENP-I. TEV protease cleavable SNAP-tag followed by DS tag was added to the C-terminus of CENP-P. CENP-LN were sub-cloned with C-terminal TEV-cleavable SNAP-DS tags on CENP-N and an N-terminal TEV-cleavable GST-tag on CENP-L.

**Expression and purification of *Bombyx mori* proteins from insect cells.**

The baculoviruses for expression of all protein complexes and mutants were generated using standard protocols based on pAcGP67 vector. All proteins were expressed in Sf9 cells which were not tested for mycoplasma contamination and were not authenticated. Typically, 4L of Sf9 cells were infected 2.5% v/v with the P3 cell culture and the cells were harvested 48-72 h after infection by centrifugation.

<u>*CENP-HIKM*</u>

Cells for CENP-HIKM and CENP-HIKM mutant modules containing Twin-Strep-tag were lysed by sonication in 50 mM Tris-HCl pH 8.0, 500 mM NaCl, 10% glycerol, 1 mM TCEP and 1 mM EDTA supplemented with protease inhibitor cocktail tablets (Roche), PMSF, benzamidine and Benzonase. Cell lysates were clarified by centrifugation at 40K rpm for 1 hour at 4°C. Supernatants were passed through a 1.5 µm syringe filter and applied to three 5ml Strep-Tactin Superflow Plus Cartridge column (Qiagen) pre-equilibrated with lysis buffer. The Strep-tactin column was washed with lysis buffer and the protein was eluted in 50 mM Tris-HCl pH 8.0, 500 mM NaCl, 10% glycerol, 1mM TCEP with 5 mM desthiobiotin (Sigma-Aldrich). The Strep-Tactin eluted fractions containing CENP-HIKM module were diluted to a final salt concentration of 300 mM NaCl and directly loaded onto HiTrap Heparin HP affinity column (Cytiva) pre-equilibrated with 25 mM Hepes pH 8.0, 0.3 M NaCl, 10% glycerol and 1mM TCEP. CENP-HIKM was eluted in 25 mM Hepes pH 8.0, 10% glycerol and 1mM TCEP with a NaCl gradient from 300mM to 2000mM NaCl.

<u>*CENP-LN*</u>

For CENP-LN and CENP-LN mutants, cells were lysed by sonication in 50 mM Tris-HCl pH 8.0, 500 mM NaCl, 10% glycerol, 1mM TCEP and 1mM EDTA supplemented with protease inhibitor cocktail tablets (Roche), PMSF, benzamidine and Benzonase. Cell lysates were clarified by centrifugation at 40K rpm for 1 hour at 4°C. Supernatants were passed through a 1.5 µm syringe filter and applied to three 5ml Strep-Tactin Superflow Plus Cartridge column (Qiagen) pre-equilibrated with lysis buffer. The Strep-Tactin column was washed with lysis buffer and the protein was eluted in 50mM Tris-HCl pH 8.0, 500mM NaCl, 10% glycerol, 1mM TCEP with 5mM desthiobiotin (Sigma-Aldrich). TEV protease was added to the Strep-Tactin eluted fraction for tag removal at 4°C for overnight incubation without rotating. The un-tagged CENP-LN sample was collected in the flow through from Ni-NTA affinity column in the buffer containing 25 mM Hepes pH 8.0, 0.3 M NaCl, 10% Glycerol, 1mM TCEP, 20 mM imidazole and directly loaded onto HiTrap Heparin HP affinity column (Cytiva) pre-equilibrated with 25 mM Hepes pH8.0, 0.3 M NaCl, 10% glycerol and 1 mM TCEP. Un-tagged CENP-LN was eluted in 25 mM Hepes pH8.0, 10% glycerol and 1 mM TCEP with a NaCl gradient from 300 mM to 2000 mM NaCl.

*CENP-OP*

For CENP-OP and CENP-OP mutants, cells were lysed by sonication in 50 mM Tris-HCl pH 8.0, 500 mM NaCl, 10% glycerol, 1 mM TCEP and 1mM EDTA supplemented with protease inhibitor cocktail tablets (Roche), PMSF, benzamidine and Benzonase. Cell lysates were clarified by centrifugation at 40K rpm for 1 hour at 4°C. Filtered lysate was applied to three 5 ml Strep-Tactin Superflow Plus Cartridge column (Qiagen) pre-equilibrated with lysis buffer. The Strep-tactin column was washed with lysis buffer and the protein was eluted in 50mM Tris-HCl pH 8.0, 500 mM NaCl, 10% glycerol, 1 mM TCEP with 5 mM desthiobiotin (Sigma-Aldrich). The Strep-Tactin eluate was dialyzed in 25 mM Hepes pH8.0, 0.3 M NaCl, 10% glycerol and 1 mM TCEP at 4°C overnight. The dialyzed CENP-OP was loaded onto HiTrap Heparin HP affinity column (Cytiva) pre-equilibrated with 25mM Hepes pH 8.0, 0.3 M NaCl, 10% glycerol and 1 mM TCEP. CENP-HIKM was eluted in 25 mM Hepes pH 8.0, 10% glycerol and 1mM TCEP with a NaCl gradient from 300 mM to 2000 mM NaCl.

*CENP-T*

For CENP-T and CENP-T mutants, cell pellets were resuspended in 50 mM Tris-HCl pH 8.0, 0.5 M NaCl, 10% glycerol, 1 mM TCEP without EDTA and supplemented with protease inhibitor cocktail EDTA-free tablets, PMSF, benzamidine and Benzonase. Cells were lysed on ice by sonication and followed by centrifugation for 1 hr at 40k rpm. The supernatant was incubated with nickel-nitriloacetic acid Ni-NTA agarose beads (Qiagen) for 3 hr with gentle rotation, followed by washing beads in 25 mM Tris-HCl pH 8.0, 1 M NaCl, 10% Glycerol, 1 mM TCEP with 20 mM imidazole. CENP-T proteins were eluted from the beads in 25 mM Tris-HCl pH 8.0, 0.3 M NaCl, 10% Glycerol, 1 mM TCEP with 300 mM imidazole. Ni-NTA eluted CENP- T was dialyzed in 25 mM Hepes pH 8.0, 0.3 M NaCl, 10% glycerol and 1 mM TCEP at 4°C overnight. Dialyzed CENP-T was further purified using a 5 ml HiTrap Heparin HP column (Cytiva) pre-equilibrated in 25 mM Hepes pH 8.0, 0.3 M NaCl, 10% glycerol, 1 mM TCEP. CENP-T were eluted with 25 mM Hepes pH 8.0, 10% glycerol, 1 mM TCEP with a NaCl gradient of 300 mM to 2000 mM.

CS module

For CS module, cells were lysed by sonication in lysis buffer containing 50 mM Tris.HCl pH 8.0, 300 mM NaCl, 0.5 mM TCEP and 10% glycerol supplemented with benzamidine, PMSF, EDTA-free protease inhibitor tablets and benzonase. The cell lysate was clarified using 1h ultracentrifuge spin at 40k rpm. Clarified lysate was loaded onto a Strep-Tactin column (Qiagen), immobilized proteins washed with lysis buffer, and the complexes were eluted in a buffer containing 50 mM Tris pH 8.0, 300 mM NaCl, 1 mM TCEP, 10% glycerol with the addition of 5 mM desthiobiotin (Sigma). TEV protease was added to cleave the Twin-Strep tag overnight. Cleaved protein was diluted to 100 mM NaCl and applied to HiTRAP HP Q column (5 ml) equilibrated with 20 mM HEPES pH 8.0, 100 mM NaCl, 10% glycerol, 1 mM TCEP. CS was eluted with a gradient of 100 mM to 600 mM NaCl over 20 cv. The peak fractions were concentrated and applied onto Superdex 200 16/600 column equilibrated in 20 mM HEPES, 300 mM NaCl, 1 mM TCEP and 10% glycerol buffer. Peak fractions were concentrated and flash-frozen in liquid nitrogen.

**Large-scale CCAN assembly for proteomics and structural studies.**

CENP-HIKM, CENP-OP, CENP-LN, with and without CS module, were purified via the Strep-Tactin column (Qiagen) individually, as described above. CENP-T was purified using Ni-NTA agarose beads (Qiagen), as described above. CENP-LN was incubated with TEV protease to remove tags overnight, while CENP-T was dialysed overnight to remove imidazole. CENP-LN was passed through Ni-NTA affinity column to remove tags and TEV protease. All components were combined and salt was adjusted to 250 mM NaCl using 25 mM HEPES pH 8.0 buffer. The complex was loaded onto HiTrap Heparin HP affinity column (Cytiva) pre-equilibrated with 25 mM HEPES pH 8.0, 250 mM NaCl, 10% glycerol and 1 mM TCEP. CCAN complex was eluted in 25 mM HEPES pH 8.0, 10% glycerol and 1mM TCEP with a NaCl gradient from 250 mM to 1000 mM. Peak fractions of CCAN complex from HiTrap Heparin column were concentrated and dialyzed overnight to reduce salt concentration in the buffer containing 20 mM HEPES pH 8.0, 300 mM NaCl, 1 mM TCEP. For CCAN-DNA structural work, 147 bp Widom 601 DNA sequence (EpiCypher) was added at an equimolar ratio to the CCAN complex prior to dialysis. After dialysis, CCAN or CCAN-DNA complex was ultracentrifuged at 50k rpm to remove any larger aggregates. For apo-bmCCAN structure, CCAN complex was on-column cross-linked using Superose 6 Increase 10/300 column (Cytiva) by injecting 0.25% glutaraldehyde (GA) at a flow rate set to 0.3ml/min for 7 ml before injecting CCAN complex. For CCAN-DNA structure, 5 mM of BS3-PEG5 chemical cross-linker was used instead of GA, but an identical procedure was followed. Lastly, the peak apo-bmCCAN or CCAN-DNA fractions were further purified using Superose 6 Increase 3.2/300 column (Cytiva) in buffer containing 20 mM HEPES pH 8.0, 300 mM NaCl, 1 mM TCEP.

**Cryo-EM grid preparation.**

3 µl of apo-bmCCAN sample or CCAN:DNA sample at $A_{280}$=1.0 (as measured by NanoDrop, ThermoFisher Scientific) was applied to Quantifoil 1.2/1.3 (300 Cu mesh) grid previously glow-discharged using Solarus plasma-cleaner for 60 sec. Grids were blotted for 3 seconds at 7 force before being plunge vitrified in liquid ethane using a MarkIV Vitrobot (ThermoFisher). The blotting chamber was maintained at 4°C and 100% humidity during freezing.

**CryoEM data collection.**

Movies were collected using a Titan Krios G3i (ThermoFisher) outfitted with a K3 camera and Bioquantum energy filter (Gatan). The K3 detector was operated in nonCDS counting mode and the energy filter slit width was set to 20 eV. Movies were collected at a nominal magnification of 105,000×, physical pixel size 0.838 Å $pixel^{-1}$, with a 50 µm C2 aperture and 100 µm objective aperture at a dose rate of 15 $e^{-}$ $pixel^{-1}$ $s^{-1}$. A total dose of 60-62 $e^{-}$ $Å^{-2}$ was collected as a 59-frame movie, resulting in a 3 s movie with 1.02-1.05 $e^{-}$ per frame.

**CryoEM data processing.**

Both apo-bmCCAN and CCAN-DNA datasets were pre-processed on-the-fly using cryoSPARC v4.0 (apo-bmCCAN) and v4.2.1 (CCAN-DNA)[56]. Movies were motion corrected using patch motion correction, CTF information was estimated using patch CTF estimation, and micrographs were curated based on CTF resolution fit. Default settings were used for all of the pre-processing steps.

For both datasets, similar initial processing steps were followed in cryoSPARC. Two datasets were processed independently. Initially, blob picking was used to over-pick all micrographs. These picked particles were extracted in a 384-pixel box, and binned to 128 pixels. 2D classification was run with 100 classes, 40 iterations and 400 particles per class to curate the particles. 'Good' (showing clear secondary structural features) class averages were selected and three *ab initio* 3D models were generated using default parameters. The best *ab initio* class was further refined via non-uniform refinement job-type. The 'good' classes were also used as templates to re-pick particles in all the apo-bmCCAN micrographs. Two rounds of heterogenous refinement were used to curate all particles with sorting against one good CCAN volume derived above and four decoy noise volumes, which were generated via *ab initio* jobs using 'bad' (clear noise or fragmented particles from the initial 2D classification jobs) particles from each dataset. The CCAN particles were subsequently non-uniform refined and then re-extracted with a 400-pixel box without binning. Non-uniform refinement was used to generate high-resolution CCAN maps which were used for per-particle motion correction as implemented in cryoSPARC to further polish the particle stack. Subsequently, different steps were used for the two datasets.

For apo-bmCCAN, 3D classification without alignment was used as implemented in cryoSPARC using principle-component analysis (PCA) initialization mode, 20 classes, and limiting resolution to 7 Å. This resulted in several distinct structural classes split over the set of 20 computational classes. The structural classes are bmCENP-LN-HIKM-T class, $bmCCAN^{CENP}$ class, and the full apo-bmCCAN. Each class was non-uniform refined to generate high-resolution maps. To improve the resolution around the mobile $bmCENP\text{-}HIK^{Head}$ domain, mask was generated around the density map in this region to exclude all density outside of the $bmCENP\text{-}HIK^{Head}$ domain. Local refinement as implemented in cryoSPARC was used to improve the density map, using pose/shift Gaussian prior set to 5 degrees and 3 Å for rotations and shifts, respectively. The local resolution map was computationally aligned on the consensus map and then combined (using Phenix 1.20.1[57]) with the consensus map into a single composite map suitable for modeling building.

For monomeric bmCCAN-DNA, 3D classification without alignment was used as implemented in cryoSPARC using simple initialization mode, 10 classes, and limiting resolution to 10 Å. Apo-bmCCAN and flexible CCAN-DNA classes were excluded, while rigid monomeric CCAN-DNA particles were further non-uniform refined. Local refinement with mask around mobile bmCENP-HIK$^{Head}$ domain was used (pose/shift Gaussian prior method with 5 degrees and 3 Å for rotations and shifts, respectively) was used to improve resolution around this region. The bmCENP-HIK$^{Head}$ domain local resolution map was computationally aligned on the consensus map of monomeric bmCCAN-DNA and then combined (using Phenix 1.20.1[57]) with the consensus map into a single composite map suitable for modeling building.

For dimeric bmCCAN-DNA, particles were re-extracted using 768-pixel box and 3x bin. 2D classification with 50 classes was used to detect larger bmCCAN-DNA dimer classes, which were selected and *ab initio* models were generated. Non-uniform refinement was used to improve the dimeric bmCCAN-DNA reconstruction. Subsequently, heterogenous refinement was used to sort all good bmCCAN particles against either monomeric bmCCAN-DNA class, dimeric bmCCA-DNA class and decoy noise volume. Dimeric bmCCAN-DNA particles were then re-extracted in 600-pixel box and binned to 400-pixel box given an effective pixel size of 1.0875 Å. Particles were moved to RELION 5.0[58], where the 3D refinement produced a consensus reconstruction. During these steps, CCAN-1 body was always well resolved while CCAN-2 body appeared to be more flexible. Mask around CCAN-2 was applied and classification without alignment in RELION 5.0 was used to classify bmCCAN-DNA dimers into three classes. One of the classes showed strong features for CCAN-2 body, and that class was further refined giving a consensus bmCCA-DNA dimer map used as final map in figure presentation and data deposition. Masks around CCAN-1, CENP-HIK$^{Head}$-T-N$^{Pyrin}$ dimers and CCAN-2 were used for MultiBody reconstruction with Blush regularization[59] of the composite bmCCAN-DNA dimer structure. CENP-HIK$^{Head}$-T-N$^{Pyrin}$ dimers body was used as the main body around which two CCAN bodies were allowed to move. The three bodies were aligned using bmCCAN-dimer consensus map and then combined using Phenix 1.20.1[57] to generate maps suitable for model building.

**Model building.**

AlphaFold2[60] was used to generate initial models for subcomplexes: CENP-OP, CENP-LN, CENP-HIKM$^{Head}$ and CENP-HIKM$^{Body}$, CS module, and CENP-T$^{HFD}$. The models were rigid body fitted into the cryoEM density maps using Chimera. Subsequently, Coot[61] was used to manually correct and re-build the complete atomic models. The complete models were then refined in Phenix (version 1.20.1)[57] using real space refinement algorithms using standard parameters.

**CS module identification from cryoEM maps.**

3.0 Å map of the apo-bmCCAN was used as input into Model Angelo software[62] in sequence-less mode (build_no_seq) and standard settings were used to generate a homology profile for each of the chains that form the CS module. Homology profile for each CS module chain was then used in HHBlits algorithm[63] to query the UniRef100 database. The top hits were then manually inspected, and the top hit for each CS protein was used in BLAST algorithm against *Spodoptera frugiperda* annotated genome to discover the sfCS proteins. SfCS protein sequences were then used to query the *Bombyx mori* database to discover the bmCS module components.

For example, the top hit from HHblits for CS-4 protein was UniRef100_A0A2A4IU08 Uncharact with P-value of 100.0. This corresponds to uncharacterized protein in *Heliothis virescens* (Tobacco Budworm Moth). The sequence of this protein is: MSALGNMVQSTQQLQQDLQNELTTSNQLLRLISIELQQIKYLTSSGGEFESNLTKNSSLV KTLANMQQVDIENLPTFTRKQIDMSMNLTQNASYEEHNTFRNESEMEESQ. This sequence was used to find *Spodoptera frugiperda* CS-4 protein (original protein in the cryoEM density map) with sequence: MSALGNLSQGNQQQDLQNELTTANQLLRLISLELEQIKNLTNPAGDFAANIKKNAELV KKLANIQQVDLTNLPPFTRKYIDLNENANQNVSYEEHNTFRNESEMDESQ. The sequence identity between the two proteins is 70%.

<u>Bombyx mori CS module sequences discovered in this study.</u>

>CS1 (KWMTBOMO01279)
MAQLQQLQQDLQNELITTNQLLHLIGNELQQIICLTSSGGEIEKNIKKNASLMSILANLQ
HIDIENIPLVTRNQVTSNQNTEELNLQRNVLDETD
>CS2 (KWMTBOMO05467)
MTSVAPSDSESNVSGFVRYSDKSFKHLYTSVLEKLELINQDILQIENNLKNLIQHAGPLE
SQLSAVLHSLPKPNTNTAMETE
>CS3 (KWMTBOMO00944)
MSNNSETSLSQRDKAAELPMELESIRALNACLQAYLEHIRNVKRNLLAMNDNYRDLED
VNRQWQEVVEFRRE
>CS4
MNSTVSDDLTRENSASEISKCISRLSANNRNLSLLHKELLEINLSVKAYVQESKKRLQGL
RLDLAHEARSSHQVVELLTEK

Note that the *B. mori* CS4 protein was not annotated in the current *B. mori* genome assembly but instead inferred from full-length cDNA clones.

**Modeling of the complete Bombyx mori holo-kinetochore.**

We used our experimentally determined bmCCAN structure and AlphaFold2-predicted bmKMN structure[60] to model complete Bombyx mori kinetochore at the molecular level (Fig. 6). Firstly, we had to assign orientation to the bmCCAN dimer with respect to the chromosome and the mitotic spindle. A unique feature of the bmCCAN is that bmCENP-T, apart from binding to the bmCENP-HIK$^{Head}$ via cryptic HFD, also binds to the bmCENP-HIK$^{Body}$ via additional α-helix (CENP-T$^{\alpha\text{-helix}}$). Because of this additional bmCENP-T$^{\alpha\text{-helix}}$, the N-terminus of CENP-T that binds bmKMN faces away from bmCCAN from the top of the loop structure as shown in Figure 6. Thus, the unique bmCENP-T binding mode to the bmCCAN defines polarity of the inner kinetochore. Secondly, we had to connect the inner and the outer kinetochore, and determine the stoichiometry of the bmKMN. Disordered bmCENP-T N-terminus is the only known bmKMN-binding region of bmCCAN, and bmCENP-T is sufficient to recruit outer kinetochore in cells [64]. AlphaFold2, however, could not predict a consistent high-confidence model for the bmCENP-T:KMN interaction, potentially suggesting that multiple CENP-T regions could bind separate bmKMN modules, similar to the human hsCENP-T. This would also be consistent with recent findings that multiple hsKMN molecules are necessary for stable kinetochore-microtubule attachments[65]. Therefore, we modeled four KMN complexes bound per bmCCAN dimer connected to the bmCENP-T in an unknown manner. We also suggest that alternative stoichiometries are possible if more than two bmCENP-T are recruited to the bmCCAN or if other uncharacterized factors contribute to outer kinetochore recruitment (Fig. 6b).

***In vitro* SEC reconstitutions.**

Purified CENP-HIKM, CENP-OP, CENP-LN, CENP-T, and CS module were analyzed in a buffer containing 25 mM HEPES pH 8.0, 300 mM NaCl, 1 mM TCEP using the microAKTA system and the Superose 6 Increase 3.2/300 column (Cytiva). All proteins were used at 2 μM concentrations for each component. Figures and figure legends describe which components were used in the reactions. Elutions were collected in the 96-well plate and similar elution fractions starting from fraction A5 to B10 were analyzed using SDS-PAGE gel (NuPAG Bis-Tris Mini Protein Gels, 4–12%, 1.0–1.5 mm, ThermoFisher Scientific) and stained with Instant Blue Coomassie stain.

**Fluorescence polarization (FP) assay.**

All fluorescence polarization measurements were conducted on an EnVision 2103 Multilabel Reader (Perkin Elmer), with excitation filter at 480 nm and emission filter at 535 nm and carried out in a ProxiPlate 384-well, round, black-bottom plate (Perkin Elmer 6008260). FP assays were performed in binding buffer of 20 mM HEPES pH 8.0, 150 mM NaCl, and 0.01% IgePAL CA-630. Fluorescenin-labeled DNA with different lengths of 15-bp, 26-bp, 36-bp, 46-bp, 53-bp, 76-bp and 96-bp sequences (IDT) were evaluated for the optimal binding of CCAN complex. The CCAN complex was serially diluted in the binding buffer containing 2.5 nM labeled DNA. The binding reaction was analyzed after overnight incubation at 4°C. The data for each condition was collected at least in technical triplicates. All data points were displayed on the graphs with standard deviation (sd) displayed as bars.

**Analysis and quantification of the FP assay.**

GraphPad Prism 10 was used for all FP data analysis and fitting. Non-linear regression analysis of the response curve (change in fluorescence anisotropy) against protein concentration produced the best fit of curves to the data from which mean half effective concentration ($EC_{50}$) values were calculated with standard error of the mean (SEM), as implemented in Prism 10. These values are stated in the text and figures for each condition. The binding equations and curves used to estimate $K_D$ values did not produce perfect fits, probably because of some degree of cooperativity during DNA binding that was difficult to mathematically approximate. Therefore, we state the obtained $EC_{50}$ values for which we have a greater confidence.

**Streptavidin bead pulldown assay.**

Streptavidin agarose beads Strep-Tactin XT 4Flow®resin (IBA Lifesciences) were washed with binding buffer containing 20 mM HEPES pH 7.5, 300 mM NaCl, 10% glycerol, 0.01% IGEPAL CA-630, 0.1 mg/ml BSA. Different CENP protein complexes, as indicated in the figures and figure legends, were incubated with streptavidin beads slurry at 30°C for 1 hr in binding buffer with gentle shaking on a thermomixer (Eppendorf). The final protein concentration for all components was 0.5 µM. The beads were pelleted and washed three times with 20 mM HEPES pH 7.5, 300 mM NaCl, 10% glycerol, 0.01% IGEPAL CA-630 to remove unbound protein, and the immobilized protein was incubated for 10 min at 30°C with elution buffer containing 20 mM HEPES pH 7.5, 300 mM NaCl, 10% glycerol, 0.01% IGEPAL CA-630, 50 mM biotin. The pre-treated and biotin-eluted samples were analyzed by NuPAGE 4-12% Bis-Tris protein gels (Invitrogen) and InstantBlue Coomassie staining. All reactions were performed at least in triplicates for quantification.

**Quantification of the pull-down assays.**

Coomassie-stained NuPAGE 4-12% Bis-Tris protein gels were analyzed using FIJI 2.9.0 software[66]. Standard protocols were used to select each lane on the gel and to quantify the protein band intensities in each lane, obtaining arbitrary intensity values. These values were normalized using band intensity for a given protein in the input lane. For example, CENP-T band intensity in the input and in the elution was estimated using this method, and the fraction of CENP-T protein bound was obtained by dividing CENP-T band intensity in the elution by CENP-T band intensity in the input. Some proteins, such as CENP-O and CENP-N, run at a very similar position on the SDS-PAGE gel. Therefore, a combined values of CENP-O/N intensity was reported, since we could not unambiguously separate these two proteins. The quantification measurements were performed across at least three replicates, and all values were plotted. The columns represent mean values with bars representing standard deviation values.

**Size Exclusion Chromatography with Multi-Angle Light Scattering (SEC-MALS).**

Molar mass analysis was performed using Agilent HPLC 1200/1220/1260/1290 series system connected to DAWN MALS detector (Wyatt Technology) and an Optilab differential refractive index (dRI) detector (Wyatt Technology). The CCAN complex was prepared as described above for large scale purification up until final purification step on SEC. Apo-bmCCAN and CCAN-DNA complexes were cross-linked in solution, centrifuged at 50k for 30 mins and passed through 0.2 µM filter before applying 20-30 µg of protein onto Bio SEC-5, 5µm, 500Å, 4.6 x 300mm column (Agilent Technologies) equilibrated with 20 mM HEPES pH 8.0, 300 mM NaCl and 1 mM TCEP at 0.3ml/min. Only dissociated sub-modules were detected in the absence of cross-linking and no accurate molecular weight measurements could be made. For CS modules, 20 µg of purified CS module was applied onto XBridge Premier Protein SEC column, 500Å, 5µm, 7.8 x 300mm (Waters Corp) in PBS (10mM Na2HPO4, 1.8mM KH2PO4, 137mM NaCl, 2.7mM KCl, pH7.2) flowing at 1 ml/min. Data were collected and analyzed using ASTRA v8.1 (Wyatt Technology). Data was plotted using GraphPad Prism.

**Protein gell band identification and confirmation.**

To identify unknown proteins using 2D LC-MS/MS, we employed a 6545XT AdvanceBio LC/Q-TOF system equipped with an Agilent PLRP-S column (2.1 × 50 mm, 5 µm, 1,000Å). SDS-PAGE gel bands of different BmCCAN complex samples were subjected to in-gel digestion using LysC and Chymotrypsin to extract peptides. The resulting peptides were then analyzed by the 2D LC-MS/MS with an electrospray ionization (ESI) source. Protein identification was performed by comparing the obtained peptide data against a known protein sequence database using Mascot MS/MS Ions Search. Further peptide mapping was conducted with the assistance of Agilent MassHunter Qualitative Analysis Software B.07.

**Insect kinetochore heparin column fractions in solution mass spec analysis.**

Label-free LC-MS/MS analysis was performed by injecting 1 µL of each fraction on an Orbitrap Eclipse mass spectrometer (ThermoFisher) coupled to a Dionex Ultimate 3000 RSLC (ThermoFisher) employing a 25 cm IonOpticks Aurora Series column (IonOpticks, Parkville, Australia) with a gradient of 2% to 30% buffer B (98% ACN, 2% H2O with 0.1% FA, flow rate = 300 nL/min). The samples were analyzed with a total run time of 95 min, FAIMS Pro DUO of -40, -60CV collected FTMS1 scans at 120,000 resolution with an AGC target of 1 x 106 and a maximum injection time of 50 ms. FTMS2 scans on precursors with charge states of 3-6 were collected at 15,000 resolution with CID fragmentation at a normalized collision energy of 30%, an AGC target of 2 x 104, a max injection time of 11 ms.

Offline search was performed using comet v.2019.01 with the following parameters: a Uniprot trichoplusia ni and spodoptera frugiperda with CENPO human sequences database 2025 version; static modification of Cys carbamidomethylation (+57.0215); variable modification of Met oxidation (+15.9949). Peptide FDR was filtered to <1% using linear discriminator algorithm.

RAW spectra files can be found on MassIVE with the dataset identifier: MSV000097156.

**Visualization.**

Chimera-1.16 and ChimeraX-1.5[67] were used for structure visualization in this study. Jalview 2.11.4.1 was used for displaying sequence alignment files. GraphPad Prism 10 was used to generate quantification of pull-down and FP assays figures.

**Lepidopteran cell lines and culture conditions.**

Cultured silkworm ovary-derived BmN4-SID1 cell lines (RRID:CVCL_Z091) [43] were maintained in Sf-900 II SFM medium (Gibco Cat#10902-088) supplemented with 10% fetal bovine serum (Eurobio Cat#CVFSVF0001), antibiotic-antimycotic (Gibco Cat#15240-062) and 2mM L-glutamine (Gibco Cat#25030-024) at 27 °C. Sf9 cells (Gibco Cat#12659017) were maintained in Sf-900 II SFM medium (Gibco Cat#10902-088) supplemented with antibiotic-antimycotic (Gibco Cat#15240-062) and 2mM L-glutamine (Gibco Cat#25030-024) at 27 °C.

**Construction of stable cell lines.**

Around 1-5 µg of plasmid DNA was transfected into $10^6$ BmN4-Sid1 (RRID:CVCL_Z091 [68] or Sf-9 cells using either XtremeGene (Roche) or Cellfectin II (Gibco Cat#10362100) according to the manufactor's instructions. For IF experiments cells were grown on coverslips before transfections. For the generation of stable polyclonal cell lines, antibiotics were added 48 hours after transfection (40 µg/ml Blasticidin (Gibco Cat#R21001)). Selection was continued until no viable untransfected cells were observed.

**Antibody generation.**

For the generation of the antibodies purified complexes of full-length *B. mori* CENP-O and CENP-P, and CS1, CS2, CS3 and CS4 protein were sent to Covalab (Villeurbanne, FR) for generation of antibodies in rabbits or mice, respectively. The specificity of the immunosignal on mitotic chromosomes detected by the antibodies was verified by RNA-mediated depletion experiments targeting the mRNAs encoding for the respective proteins.

**Plasmid construction.**

For the kinetochore IPs, full-length ORFs of *S. frugiperda* CENP-O, CENP-P, CS1 and CS2 fused to 3xFLAG tags were cloned into pIBV5 (Invitrogen Cat#12550018) using Gibson assembly.

**Affinity co-immunoprecipitations.**

Cultures of the Sf9 strains expressing full-length *S. frugiperda* kinetochore proteins fused to a C-terminal 3XFLAG tags or control wild-type Sf9 cells were grown to exponential phase in Sf-900 II SFM medium (Gibco Cat#10902-088). For each strain, 100 ml of 5-6 x $10^9$ cells were harvested by centrifuging for 10 min at 300 *xg*, and the pellets were washed twice in cold PBS. The pellets were resuspended in 3 mL HDG150 Buffer (20 mM HEPES pH 7.0, 150 mM KCl, 10% glycerol, 0.5 mM DTT, 1 tablet cOmplete Protease Inhibitor), and then cells were disrupted with 50 strokes in a dounce homogenizer at 4 °C. The dounced fraction was centrifuged at 1700 *xg* for 10 min at 4 °C, and the nuclei (lower fraction) were gently resuspended with 2 mL HDG150 Buffer and re-centrifuged in the same conditions. The nuclei were resuspended in a final volume of 2 mL using HDG150 Buffer. Nuclear extracts were prepared by passing the nuclear fraction 10 times through a 20G 1 ½" needle (0.9 x 38 mm) and then 5 times through a 25G 3/8" needle (0.5 x 16 mm). The nuclear extracts were centrifuged at 20 000 *xg* for 10 min at 4 °C in microcentrifuge tubes. The pellets were resuspended in HDG150 Buffer and pooled together at a final volume of 2 mL. To prepare the chromatin, the nuclear extracts were digested with ~10 units MNase (Sigma Cat#N3755-500UN) for 1 hour at 4 °C on a roller, 3 mM $CaCl_2$ was added to the digestions. The MNase digestions were stopped by adding 50 µL of 0.2 M EGTA. To solubilize the digested chromatin, 4 mL of HDG400 Buffer (20 mM HEPES pH 7.0, 400 mM KCl, 10% glycerol, 1 mM DTT, 0.05% NP-40, 1 tablet cOmplete Protease Inhibitor) was added to the samples and incubated for 2 h at 4 °C on a roller. The samples were centrifuged at 8000 *xg* for 10 min at 4 °C. The supernatants were saved to bind to the anti-FLAG M2 beads (Sigma Cat#M8823; RRID:AB_2637089). The M2 magnetic beads were prepared according to the manufacturer's recommendations. The digested chromatin samples were incubated with 30 µL anti-FLAG M2 beads. The beads were washed four times with 1 mL HDGN320 Buffer (20 mM HEPES pH 7.0, 320 mM KCl, 10% glycerol, 1 mM DTT, 0.05% NP-40, 1 tablet cOmplete Protease Inhibitor). For proteomic analyses, IP samples were washed trice with 100 µL of 100 mM $NH_4HCO_3$ (ABC) and proteins were trypsin/lysC (0.2 µg, Promega) digested directly on beads in a total volume of 100 µL of 100 mM ABC for one hour at 37 °C while vortexed at 1000 rpm. Samples were then cleaned on homemade C18 StageTips (AttractSPE Disk Bio C18-100.47.20 Affinisep), before being eluted using 40/60 CH3CN/H2O + 0.1% formic acid and vacuum concentrated to dryness.

**Proteomics and Mass Spectrometry Analysis.**

LC-MS/MS analysis: data independent acquisition (DIA) analyses were performed with a RSLCnano system (Ultimate 3000, Thermo Scientific) coupled to an Orbitrap Exploris 480 mass spectrometer (MS), interfaced by a Nanospray Flex ion source (Thermo Scientific). Peptides were trapped on a 2 cm nanoviper Precolumn (75 µm inner diameter x 2 cm, C18 Acclaim PepMapTM 100, Thermo Scientific) at a flow rate of 2.5 µL/min in 98% buffer A (2/98 CH3CN /H2O + 0.1% formic acid) for 4 min to desalt and concentrate the samples. Peptides were then injected onto a C18 column (75 µm inner diameter x 50 cm double nanoViper PepMap Neo, 2µm, 100Å, Thermo Scientific) regulated at a temperature of 50°C, and separated with a linear gradient from 98% buffer A (2/98 CH3CN /H2O + 0.1% formic acid) to 30% buffer B (98/2 CH3CN /H2O + 0.1% formic acid) at a flow rate of 300 nL/min over 91 min. Peptides were analyzed in the MS applying a 2400V spray voltage, funnel RF level at 40% and a heated capillary temperature set to 280°C. MS full scans were recorded in centroid mode for ranges 375-1500 m/z with a resolution of 120,000 at m/z 200, a normalized AGC target set at 300% and a maximum injection time set to auto. The MS2 acquisitions were performed in centroid mode and on an auto range scan mode after fragmentation using HCD with 25% normalized collision energy, a normalized AGC target of 3000% and a resolution of 15,000. A window width of 15 Th with 1 Th overlap, 40 scan events and a maximum injection time set to auto were chosen for DIA on a 400-1000 m/z precursor mass range.

Data processing: For identification, the data were searched against the *Spodoptera frugiperda* corn annotation OGS6.1 database (https://bipaa.genouest.org/sp/spodoptera_frugiperda_pub/) manually curated to include the sequences of *S. frugiperda* CENP-K and CENP-N using Pulsar search engine through Spectronaut v18.7 (Biognosys) by directDIA+ analysis using default search settings. Enzyme specificity was set to trypsin and a maximum of two missed cleavage sites was allowed. N-terminal acetylation and oxidation of methionine were set as variable modifications. The resulting files were further processed using myProMS v3.10. https://github.com/bioinfo-pf-curie/myproms[69].

For protein quantification, MS2 XICs from proteotypic peptides shared between compared conditions (TopN matching) were used with missed cleavages allowed. Median and scale normalization at peptide level was applied on the total signal to correct the XICs for each biological replicate (N =2 or 3). To evaluate the statistical significance of the change in protein abundance, a linear model (adjusted on peptides and biological replicates) was performed, and a two sided T-test was apply on the fold change estimated by the model. The p-values were then adjusted using the Benjamini–Hochberg FDR procedure.

The mass spectrometry proteomics raw data have been deposited to the ProteomeXchange Consortium via the PRIDE[70] partner repository with the dataset identifier PXD063391.

**Analyses MS data.**

The hits for each IP-MS were generated applying the following filters in the dataset- fold change over control: 2 across the replicates, number of unique peptides: >2 in all three replicates. From the individual IP hits, those that are common between any sets of proteins, and (CENP-O, -P, CS1, and CS2: refer to sheet 'consolidated') were identified and listed in the additional sheets in the supplementary table. Description of protein function was added based on either known function at the kinetochore, or based blast searches to the human or *Drosophila melanogaster* proteomes.

**Immunofluorescence.**

Cells were grown on glass coverslips and fixed with 4% PFA (anti-CENP-T, anti-CENP-OP, anti-tubulin), ice cold acetone (anti-Dsn1, anti-CS1-4, anti-SPC24/25) and 4% PFA (anti-tubulin), followed by permeabilization using 0.3% Triton X-100 in PBS and blocked in 3% BSA-PBS. The following antibodies were used: rabbit polyclonal anti-*B. mori* CENP-T[64], mouse polyclonal anti-*B. mori* CENP-T[71] rabbit polyclonal anti-*B. mori* Spc24/25[64], polyclonal rabbit anti-*B. mori* Dsn1[64], and polyclonal mouse anti-*B. mori* CS1-4 and polyclonal rabbit anti-*B. mori* CENP-OP generated by Covalab (Villeurbanne, FR) at the dilution 1:1000, anti-α-tubulin monoclonal Alexa Fluor 488 (Thermo Fisher Scientific Cat#53-4502-80; RRID:AB_1210526) at 1:1000, anti-FLAG M2 mouse monoclonal (Sigma Cat#F1804; RRID:AB_262044) at 1:1000, anti-phospho Histone H3-Ser10 rat monoclonal (Sigma Cat#MABE939) at 1:1000. For fluorescent conjugated secondary antibodies, we used goat anti-rabbit IgG Alexa Fluor 568 (Thermo Fisher Scientific Cat#A-11011; RRID:AB_143157) at 1:1000, goat anti-rat IgG Alexa Fluor 568 (Thermo Fisher Scientific Cat#A-11077; RRID:AB_2534121) at 1:1000, goat anti-rat IgG Alexa Fluor 488 (Thermo Fisher Scientific Cat#A-11006; RRID:AB_2534074) at 1:1000, goat anti-mouse IgG Alexa Fluor 488 (Thermo Fisher Scientific Cat#A-11029; RRID:AB_2534088) at 1:1000, goat anti-mouse IgG Alexa Fluor 568 (Thermo Fisher Scientific Cat#A-11004; RRID:AB_2534072) at 1:1000 and goat anti-rat IgG Alexa Fluor 633 (Thermo Fisher Scientific Cat#A-21094; RRID:AB_2535749) at 1:1000. DNA was stained with DAPI (Sigma Cat#D9542) and samples were mounted in Vectashield Antifade Mounting Medium (Vector Laboratories Cat# H-1000; RRID:AB_2336789).

For anti-tubulin staining cells were fixed three and five days after RNAi-mediated depletion using a protocol for the preservation of the whole cytoskelon [72]. Cells were washed with PBS for 5 minutes, then incubated for 10 min at room temperature in 1 mM dithiobis(succinimidyl propionate, DSP) (Thermo Fisher Scientific Cat#22585) in Hank's balanced salt solution (HBSS) (Gibco Cat#14025050), followed by an incubation for 10 min at room temperature in 1 mM DSP in microtubule-stabilizing buffer (MTSB). Cells were next washed for 5 min in 0.5% Triton X-100 in MTSB and then fixed in 4% PFA in MTSB for 15 min at room temperature. After a 5 min wash in PBS, cells were incubated for 5 min in 100 mM glycine in PBS, then washed again in PBS for 5 minutes and finally nuclei were stained with DAPI (Sigma Cat#D9542) and samples were mounted in Vectashield Antifade Mounting Medium (Vector Laboratories Cat# H-1000; RRID:AB_2336789).

**Microscopy.**

Images for quantification of CS complex localization, co-stained with BmCENP-T and BmDsn1 and upon depletion of other kinetochore subunits, and for BmCENP-OP localization upon depletion of CS complex, were acquired in Leica Thunder microscope using a 63x objective (oil immersion, NA 1.4). The acquisition parameters are as follows: DAPI – 390nm, excitation filter- 375-405 nm; emission filter- 420-450nm; GFP (H3S10p)- 475nm, excitation filter- 462-496 nm; emission filter- 506-532, and Red (CS proteins) – 555nm , excitation filter- 542-566 nm; emission filter- 578-610 nm. Signals were detected using a Leica k8 CMOS black thinned camera

(2048*2048 pixels, pixel size 6.5* 6.5 um), and sensor size (diagonal) of 18.8 mm. Quantification of fluorescence intensity was performed using the Fiji software[66] on unprocessed TIFF images. Mitotic cells (H3S10ph positive) were quantified. For RNAi depleted cells, H3S10p signals were then used as markers to manually select, using the freehand selections tool, the nuclear area. The mean fluorescence intensity of each nucleus was measured and corrected for background. For background correction, the average of mean intensities of three random circular regions of fixed size (30x30 pixels) placed outside nuclear areas was determined and subtracted from the IF signal of each nucleus. For statistical analysis a student's t-test (unpaired, unequal variance) was used. Differences were considered statistically significant at values of P values <0.05.

**RNAi-mediated knock-down.**

BmN4-SID1 cells were grown on coverslips and incubated with 400pg/μl dsRNA for three days. After three days, the medium was change to add another 400pg/μl dsRNA. After five (CENP-T, CENP-H, CENP-I, CENP-K) or eight days (CENP-OP, CS1234), cells were fixed and processed for IF as described. 200-400 bp dsRNAs were generated from DNA templates fused to T7 promoter using MAXIscript T7 Transcription kit (Thermo Fisher Scientific, Cat#AM1312).

**Life-cell microscopy.**

*Bombyx mori* BmN4-Sid1 cells were cultured in a 2 mL FluoroDish (World Precision Instruments, Cat# FD35-100). Cells were treated with Control and CS dsRNAs for 3-5 days. Two hours prior to imaging, SPY-DNA-555nm (Spirochrome Cat# SC201) and SPY-tubulin-650nm (Spirochrome Cat# SC503) dyes were added to the medium at a 10,000X dilution. Cells were imaged using the spinning disk confocal microscope[73]. Briefly, the Nikon Eclipse Ti-E perfect focus inverted microscope was coupled to the Yokogawa CSU-X1 spinning disk confocal unit (Yokogawa). The microscope was enclosed within a thermal box to keep stable temperatures of 27°C. It is equipped with 40X/1.3 N.A. Plan Apo oil immersion objective lens and Mad City Piezo stepper stage, the Photometrics Evolve EM-CCD camera, the Gataca Systems laser unit with 561 nm (100 mW) and 605 nm (100 mW) lines, and controlled by Molecular Devices software MetaMorph 7.8. Movies were made with the following parameters: laser power 10% SPY-DNA, 20% SPY-Tubulin, EM-gain 300, Bin 1X, exposure time 100-200 ms, 13 optical z-sections, 2 μm spacing per 3D stack, 5 min time interval between stacks, 15 hr movies. Mitotic duration for Figure 3 was determined as the time taken for anaphase onset after spindle pole body duplication (min) in control and CS depleted cells.

**Cross-linked ChIPs using in-house protocol.**

Cross-linked (X-ChIP) was performed as previously described[74] with the some modifications. Two confluent T75 flasks (Thermofisher, catalog # 156499) of BmN4 cells were used for each ChIP. Cells were cross-linked in freshly prepared 1% MeOH-free formaldehyde (Thermofisher catalog # 28906) for 10 min at room temperature. Cross-linking was quenched by adding glycine to 125 mM for 2 min at room temperature. Cells were then washed in ice-cold PBS and incubated for 10 min with 150 μl ice-cold lysis buffer (1% SDS, 10 mM EDTA, 50 mM Tris-HCl pH 8.1, 1 X cOmplete Protease Inhibitor Cocktail (Roche catalog # 11697498001)). To the cell lysates, 1350 μl ChIP buffer (1% Triton X-100, 150 mM NaCl, 2 mM EDTA, 20 mM Tris-HCl pH 8.1, 1 X Protease inhibitor cocktail) was added along with 4.5 μl CaCl2 1 M (3 mM final) and then pre-warmed for 2 min at 37 °C. Nuclei were then treated with 1 or 2 units of MNase (Sigma, catalog # N3755-500UN) for 15 min at 37 °C. MNase reaction was stopped by adding a mix of 30 µl EDTA (0.5 M stock) and 60 µl EGTA (0.5 M stock). Each MNase-treated nuclei sample was then sonicated using a Covaris E220 sonicator under the following parameters: 150 sec, Duty 10%, Power 75 W, cycles/burst 200, 7 °C. Sonicated chromatin was centrifuged 3 min at 16000 *g* and clear supernatant containing the solubilized chromatin was saved either as input or for ChIP. anti-CENP-T serum (rabbit polyclonal) and anti-CS1-4 serum (mouse polyclonal) were incubated with Protein A dynabeads (Thermofisher, catalog # 10001D) for 10 min at room temperature to allow for beads-antibody binding. Antibody-bound beads were washed with ChIP buffer and mixed with input chromatin. All samples were incubated overnight at 4 °C. Chromatin-bound beads were collected the next day on a magnetic rack and washed with the following ice-cold buffers: once with low-salt TSE 1 (0.1% SDS, 1% Triton X-100, 150 mM NaCl, 2 mM EDTA, 20 mM Tris-HCl pH 8.1,); four times with high-salt TSE II (0.1% SDS, 1% Triton X-100, 500 mM NaCl, 2 mM EDTA, 20 mM Tris-HCl pH 8.1); and three times with 1x TE. DNA was directly extracted from chromatin-beads or input by adding DNA extraction buffer (20 mM Tris-HCl pH 8.1, 10 mM EDTA, 5 mM EGTA, 300 mM NaCl, 1% SDS) and incubating at 37 °C followed by reversing cross-links by addition of proteinase K (Qiagen, catalog # 19131) and incubation overnight at 65°C. DNA was extracted with Phenol:Chloroform and precipitated with NaOAc and 100% EtOH in the presence of 1 µl 15 mg/ml Glycoblue (Ambion, catalog # AM9515). DNA was finally re-suspended in 1x TE containing RNase A (1µg/µl) (QIAGEN, catalog # 19101) and incubated for 15 min at 37 °C. Nucleosome profiles for Input and ChIP DNA were analyzed using an Agilent 4200 Tapestation with a DNA high sensitivity kit.

**Next-generation sequencing and ChIP-seq data analysis.**

All steps of Illumina library preparation and sequencing were carried out at the Curie Institute's sequencing platform. Adapter trimmed, single-end Illumina reads of 100 bp length were mapped using Bowtie2[75] with default parameters to the *B. mori* genome assembly downloaded from Silkbase: http://silkbase.ab.a.u-tokyo.ac.jp, which was modified to extract only assembled chromosomes 1 to 28. After removal of duplicates using Picard tools (http://broadinstitute.github.io/picard/), Deeptools bamCompare function[76] was as used to generate ChIP-seq signal tracks represented as histograms of the average log2-ratio of RPKM-normalized read counts in IP over Input in genome-wide 1 kb windows that were visualized in IGV [77]. Deeptools multiBigWigSummary function was used to compute average log2-ratio of RPKM-normalized read count in IP over Input in genome-wide 10 kb windows for making scatterplots (Wickham H (2016), https://ggplot2.tidyverse.org) of the correlation between different ChIP-seq targets. Pearson correlation was calculated in Rstudio (Rstudio Team (2016). Rstudio: Integrated Development for R. Rstudio, Inc., Boston, MA URL http://www.rstudio.com/) after filtering out those 10 kb windows with zero mapped reads in both IP and Input.

**Homology predictions, alignments and phylogenetic analyses.**

To test for homology of the newly identified CS proteins to components of the Dam1 complex or to any other protein HHpred version 3.2.0 searches[78] were performed against the PDB_mmCIF70_3_Aug.

The protein sequences of Dam1 complex members published in Rooijen[79] were used for phylogenetic analyses including the newly identified putative homologs. The sequences of each orthologous group were aligned using MAFFT E-INS-I[80] (MAFFT v7). For inferring building the tree, the MSAs of the subunits of Dam1-C were aligned by using MAFFT merge E-INS-i. IQ-TREE[81] was then used to select a substitution model as advised by ModelFinder[82] and to infer the phylogeny using 1000 ultrafast bootstraps. The tree was visualized using iTOL[83].

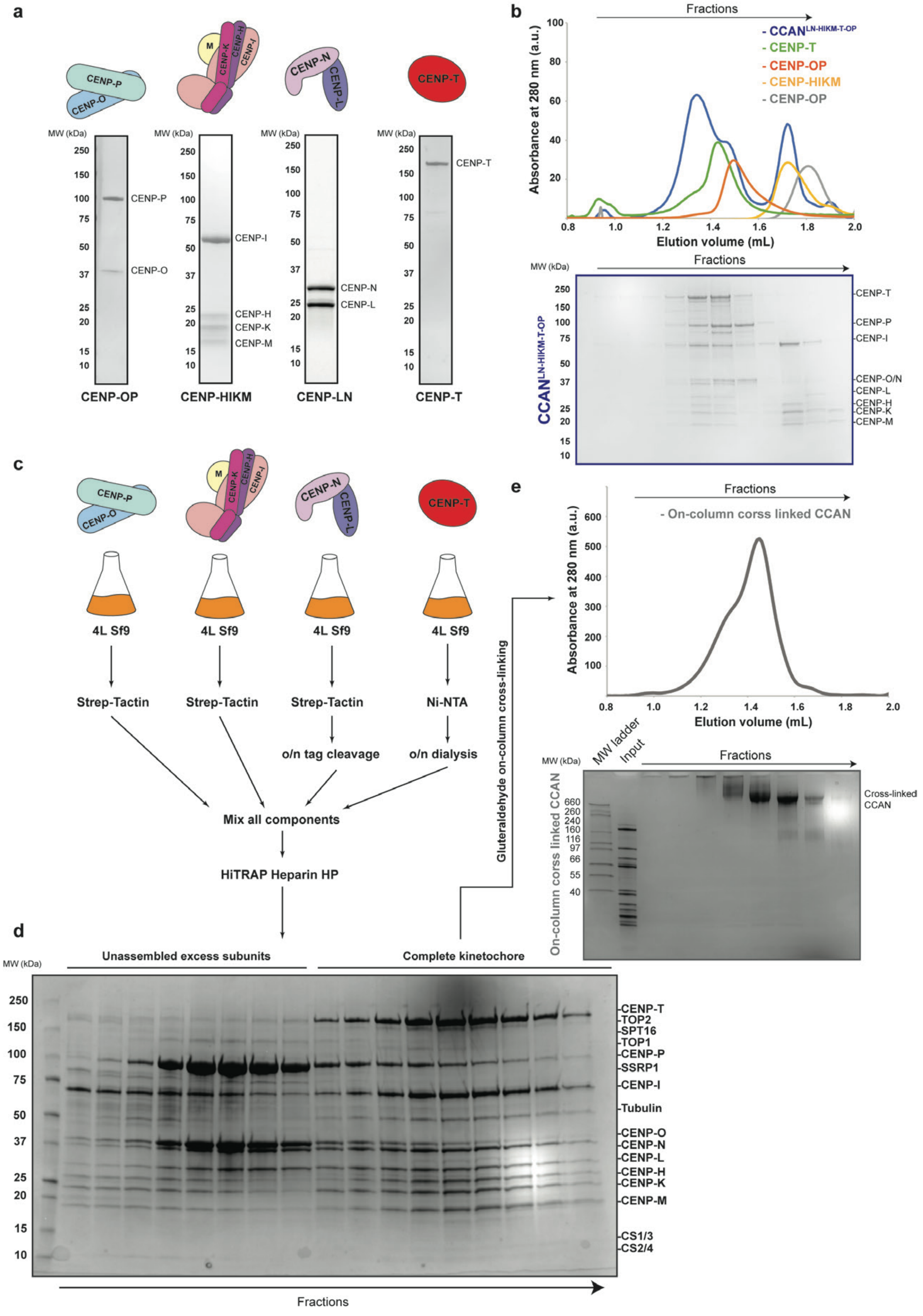
a
CENP-P
CENP-O
M
CENP-K
CENP-H
CENP-I
CENP-N
CENP-L
CENP-T
MW (kDa)
250
150
100
75
50
37
25
20
15
10
CENP-P
CENP-O
CENP-I
CENP-H
CENP-K
CENP-M
CENP-N
CENP-L
CENP-T
CENP-OP
CENP-HIKM
CENP-LN
CENP-T
b
Fractions
- CCAN^LN-HIKM-T-OP
- CENP-T
- CENP-OP
- CENP-HIKM
- CENP-OP
Absorbance at 280 nm (a.u.)
100
80
60
40
20
0.8
1.0
1.2
1.4
1.6
1.8
2.0
Elution volume (mL)
MW (kDa)
Fractions
CCAN^LN-HIKM-T-OP
-CENP-T
-CENP-P
-CENP-I
-CENP-O/N
-CENP-L
-CENP-H
-CENP-K
-CENP-M
c
4L Sf9
Strep-Tactin
Strep-Tactin
Strep-Tactin
Ni-NTA
o/n tag cleavage
o/n dialysis
Mix all components
HiTRAP Heparin HP
Gluteraldehyde on-column cross-linking
e
Fractions
- On-column corss linked CCAN
Absorbance at 280 nm (a.u.)
600
500
400
300
200
100
0.8
1.0
1.2
1.4
1.6
1.8
2.0
Elution volume (mL)
MW (kDa)
MW ladder
Input
Fractions
On-column corss linked CCAN
660
260
240
160
116
97
66
55
40
Cross-linked CCAN
d
MW (kDa)
Unassembled excess subunits
Complete kinetochore
250
150
100
75
50
37
25
20
15
10
-CENP-T
-TOP2
-SPT16
-TOP1
-CENP-P
-SSRP1
-CENP-I
-Tubulin
-CENP-O
-CENP-N
-CENP-L
-CENP-H
-CENP-K
-CENP-M
-CS1/3
-CS2/4
Fractions

2

**Extended Data Fig. 1. BmCCAN purification and reconstitution.**

**a,** SDS-PAGE gel of purified bmCCAN components. **b,** Reconstitution of the apo-bmCCAN using purified components shown in **a,**. The reconstitution was performed using a Superose 6 3.2/300 column, and chromatograms for each module were plotted. SDS-PAGE gel for the full CCAN assembly is shown below, with gel fractions aligned to the chromatogram above. The experiment has been repeated at least in triplicate with similar results. **c,** Schematic for the combined CCAN purification from Sf9 cells. Each module was independently overexpressed in Sf9 cells and purified using the corresponding tag during the first purification step (Methods). The purified components were mixed together and applied to the Heparin HP column. **d,** SDS-PAGE gel of the elution fractions from the Heparin column is shown. Each band has been identified and confirmed using mass-spectrometry analysis (Methods). Certain bands could not be unambiguously identified and so were not annotated. **e,** Heparin fractions that contain complete bmCCAN were used for on-column cross-linking using a Superose 6 3.2/300 column, with the chromatogram for the elution shown. SDS-PAGE gel of the on-column cross-linked sample is shown below.

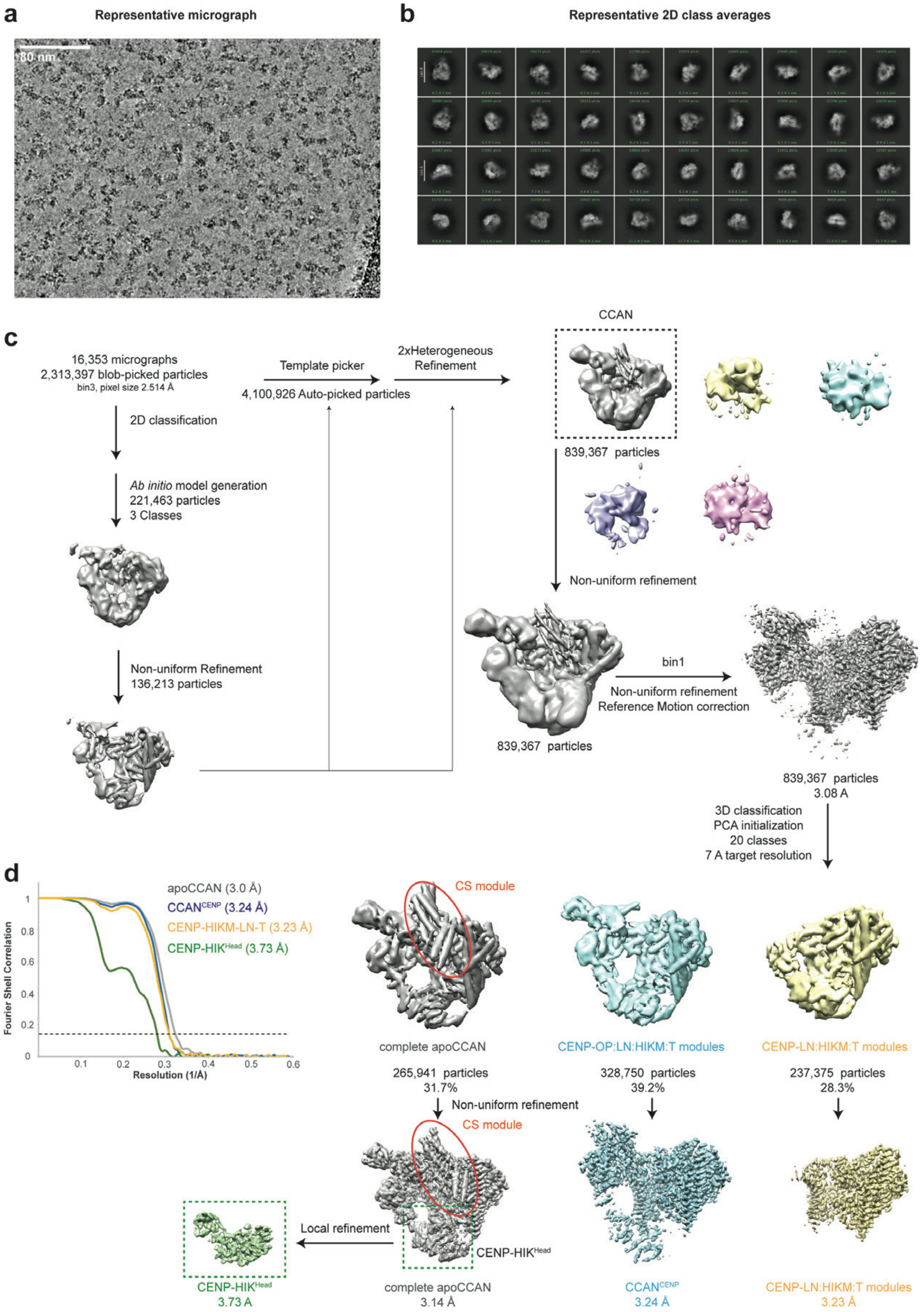
a
Representative micrograph
80 nm
b
Representative 2D class averages
c
16,353 micrographs
2,313,397 blob-picked particles
bin3, pixel size 2.514 Å
2D classification
Ab initio model generation
221,463 particles
3 Classes
Non-uniform Refinement
136,213 particles
Template picker
4,100,926 Auto-picked particles
2xHeterogeneous Refinement
CCAN
839,367 particles
Non-uniform refinement
839,367 particles
bin1
Non-uniform refinement
Reference Motion correction
839,367 particles
3.08 A
3D classification
PCA initialization
20 classes
7 A target resolution
d
apoCCAN (3.0 Å)
CCANCENP (3.24 Å)
CENP-HIKM-LN-T (3.23 Å)
CENP-HIKHead (3.73 Å)
Fourier Shell Correlation
Resolution (1/Å)
CS module
complete apoCCAN
CENP-OP:LN:HIKM:T modules
CENP-LN:HIKM:T modules
265,941 particles
31.7%
328,750 particles
39.2%
237,375 particles
28.3%
Non-uniform refinement
CS module
Local refinement
CENP-HIKHead
CENP-HIKHead
3.73 A
complete apoCCAN
3.14 Å
CCANCENP
3.24 Å
CENP-LN:HIKM:T modules
3.23 Å

**Extended Data Fig. 2. BmCCAN cryoEM data collection and processing.**

**a,** Representative cryoEM micrograph for the on-column cross-linked apo-bmCCAN sample. **b,** Representative 2D class averages of the apo-bmCCAN complex generated using cryoSPARC. **c,** CryoEM data processing workflow for the apo-bmCCAN dataset (Methods). The blob picker, as implemented in cryoSPARC, was used to perform initial particle picking. These particles were 2D classified to select the best particle set with classes that contain clear secondary structural features. The best particles were used to generate an *ab initio* model which was further refined. The 2D classes were also used for template picking to improve particle picking, and heterogeneous refinement was used to separate "true" bmCCAN particles from broken or noise particles. The particles that correspond to the bmCCAN were further refined and classified without alignment to isolate different bmCCAN assembly intermediates as described in the text. Local refinement was used to improve the resolution of the CENP-HIK$^{Head}$ group in the complete apo-bmCCAN reconstruction. **d,** Fourier Shell Correlation (FSC) curves for all reported apo-bmCCAN reconstructions, with the grey dotted line representing 0.143 value.

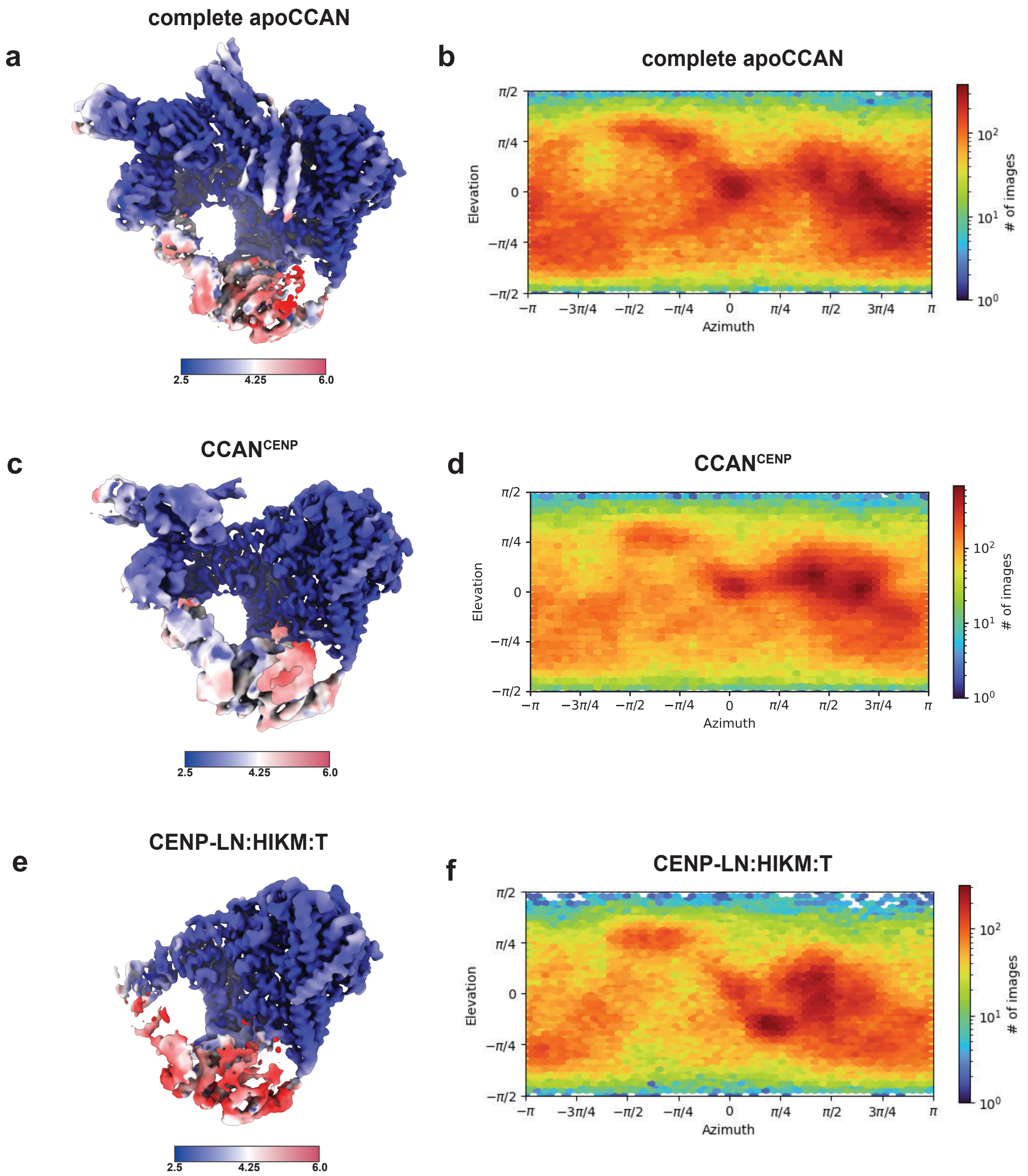


**Extended Data Fig. 3. BmCCAN cryo-EM reconstruction parameters.**
**a,** CryoEM reconstruction of the apo-bmCCAN colored by resolution, with the resolution range shown below. **b,** Angular distribution of views for the apo-bmCCAN reconstruction. **c,** CryoEM reconstruction of the bmCCAN$^{CENP}$ (bmCENP-LN-HIKM-T-OP) colored by resolution, with the resolution range shown below. **d,** Angular distribution of views for the bmCCAN$^{CENP}$ reconstruction. **e,** CryoEM reconstruction of the CENP-LN-HIKM-T subcomplex colored by resolution, with the resolution range shown below. **f,** Angular distribution of views for the CENP-LN-HIKM-T subcomplex reconstruction.

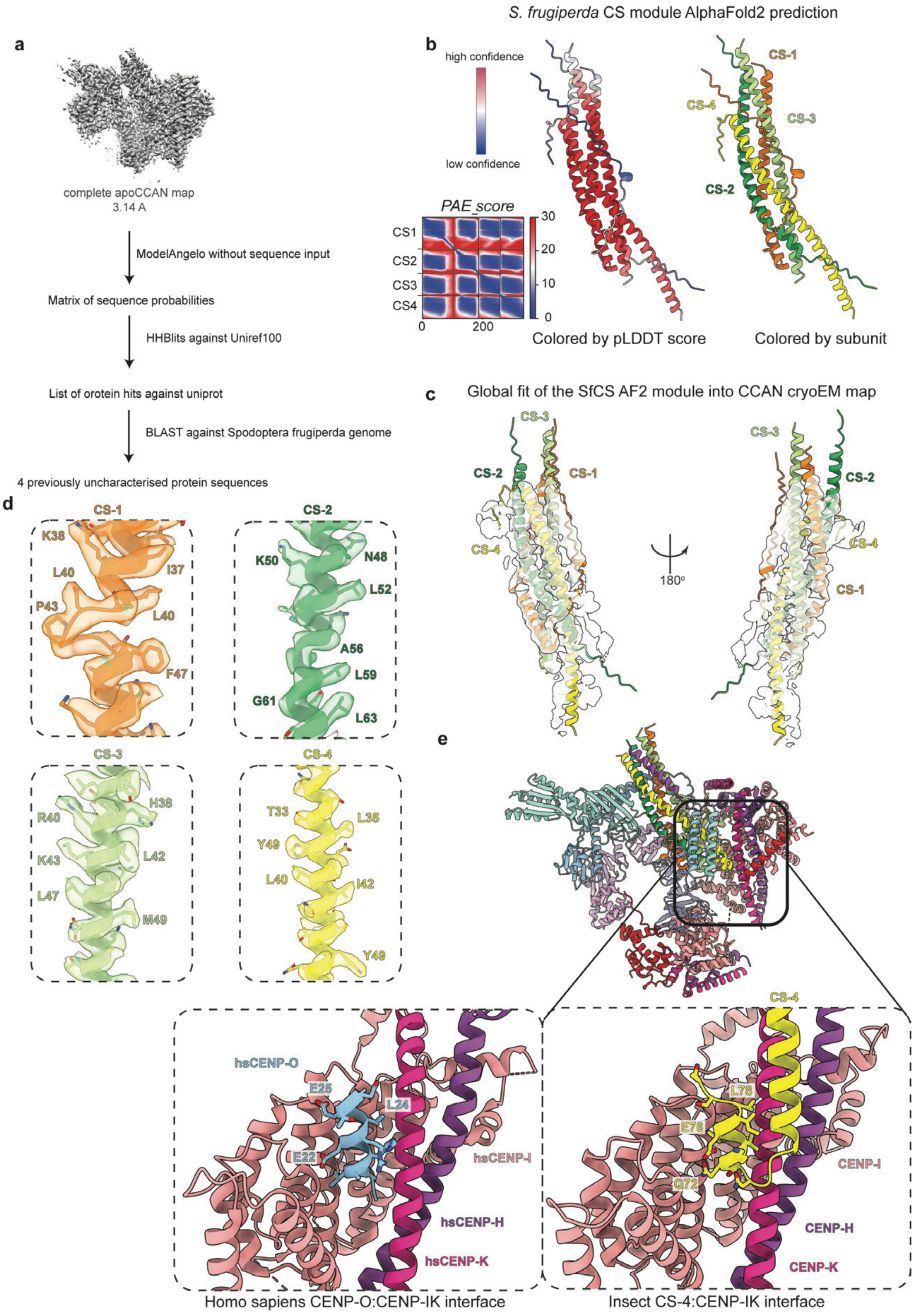

S. frugiperda CS module AlphaFold2 prediction
a
complete apoCCAN map
3.14 A
ModelAngelo without sequence input
Matrix of sequence probabilities
HHBlits against Uniref100
List of orotein hits against uniprot
BLAST against Spodoptera frugiperda genome
4 previously uncharacterised protein sequences
b
high confidence
low confidence
PAE_score
CS1
CS2
CS3
CS4
0
200
30
20
10
0
Colored by pLDDT score
CS-1
CS-4
CS-3
CS-2
Colored by subunit
c
Global fit of the SfCS AF2 module into CCAN cryoEM map
CS-3
CS-2
CS-1
CS-4
180°
d
CS-1
K38
L40
I37
P43
L40
F47
CS-2
K50
N48
L52
A56
L59
G61
L63
CS-3
H38
R40
L42
K43
L47
M49
CS-4
T33
L35
Y49
L40
I42
Y49
e
hsCENP-O
E25
L24
E22
hsCENP-I
hsCENP-H
hsCENP-K
CS-4
L78
E76
Q72
CENP-I
CENP-H
CENP-K
Homo sapiens CENP-O:CENP-IK interface
Insect CS-4:CENP-IK interface


2

**Extended Data Fig. 4. Identification and validation of the CS module subunits.**
**a,** The workflow used to identify CS module protein sequences, described in detail in the Methods section. **b,** AlphaFold2 structure prediction of the *S. frugiperda* CS module using sequences identified in **a,**. The model on the left is colored by the pLDDT score, with PAE values plotted for protein-protein contacts. The model colored by subunit is shown on the right. **c,** AlphaFold2 model of the *S. frugiperda* CS module fit into the apo-bmCCAN cryoEM reconstruction. **d,** CS module side-chain density from the apo-bmCCAN cryoEM reconstruction with the fitted molecular model of the *S. frugiperda* CS module protein sequences identified in **a**. **e,** Comparison of the apo-bmCCAN CS-4:CENP-IK interface on the right with the human CENP-O:CENP-IK interface (PDB ID: 7R5S)[84] on the left.

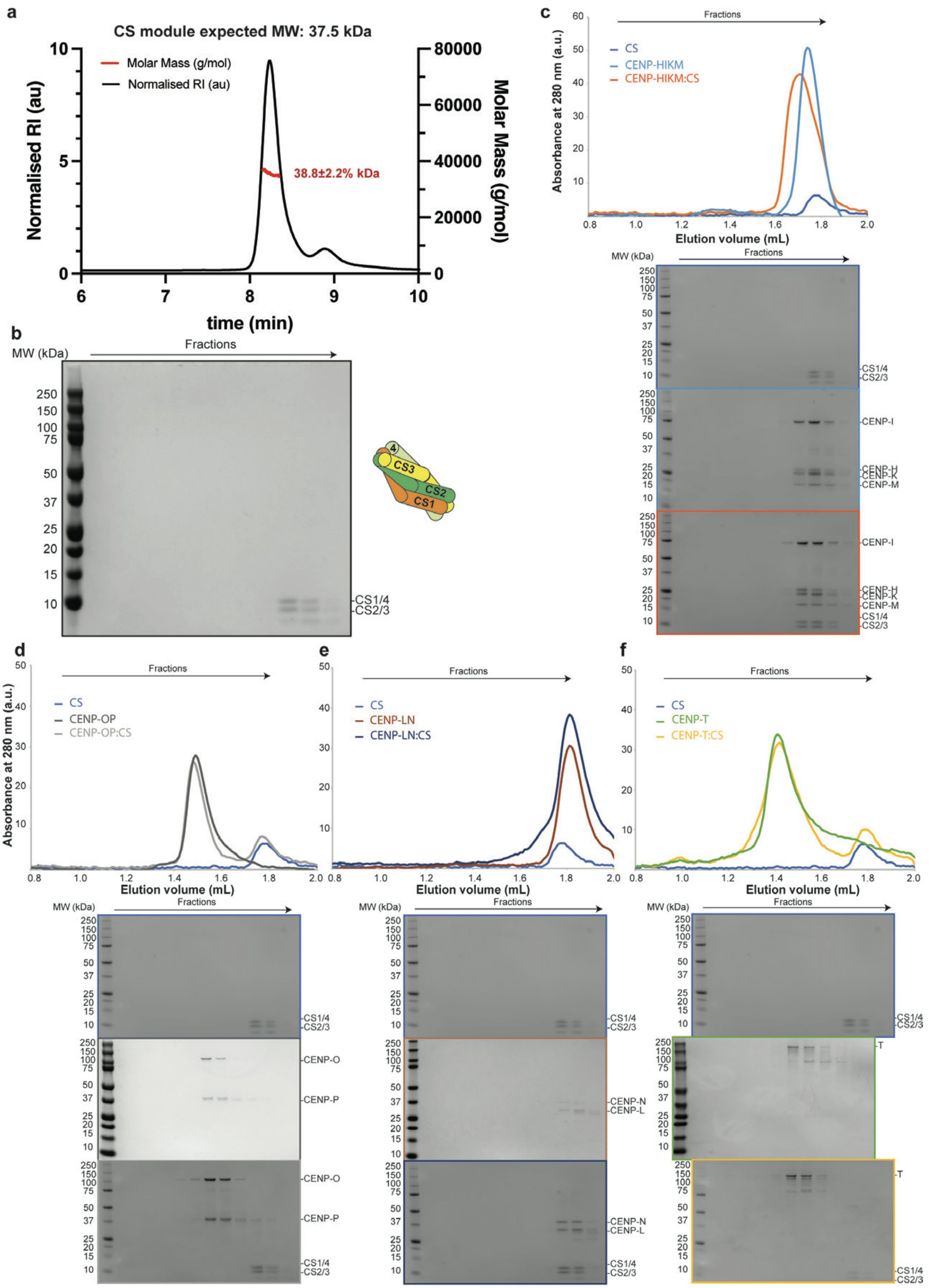
a
CS module expected MW: 37.5 kDa
Molar Mass (g/mol)
Normalised RI (au)
38.8±2.2% kDa
Normalised RI (au)
Molar Mass (g/mol)
time (min)
b
MW (kDa)
Fractions
CS1/4
CS2/3
CS3
CS2
CS1
4
c
Fractions
CS
CENP-HIKM
CENP-HIKM:CS
Absorbance at 280 nm (a.u.)
Elution volume (mL)
MW (kDa)
Fractions
CENP-I
CENP-H
CENP-K
CENP-M
CS1/4
CS2/3
d
Fractions
CS
CENP-OP
CENP-OP:CS
Absorbance at 280 nm (a.u.)
Elution volume (mL)
MW (kDa)
CENP-O
CENP-P
CS1/4
CS2/3
e
Fractions
CS
CENP-LN
CENP-LN:CS
Elution volume (mL)
MW (kDa)
CENP-N
CENP-L
CS1/4
CS2/3
f
Fractions
CS
CENP-T
CENP-T:CS
Elution volume (mL)
MW (kDa)
T
CS1/4
CS2/3

**Extended Data Fig. 5. *Bombyx mori* CS module biochemical characterization.**
**a,** SEC-MALS analysis of the purified *Bombyx mori* CS module plotted with normalized refractive index (RI) and molecular mass on Y-axes against the elution time of complexes on the X-axis. The estimated molecular weight of the CS module is 37.5 kDa. The measured molecular weight of the CS module is 38.8 ± 2.2% kDa with 1.0 ± 3.1% polydispersity (Mw/Mn) value. **b,** SDS-PAGE gel analysis of the *Bombyx mori* CS module eluted from Superose 6 3.2/300 column. **c-f,** Reconstitution of the purified CS module with each component of the $bmCCAN^{CENP}$ complex using Superose 6 3.2/300 column with SDS-PAGE gels for each module and the complexes shown below.

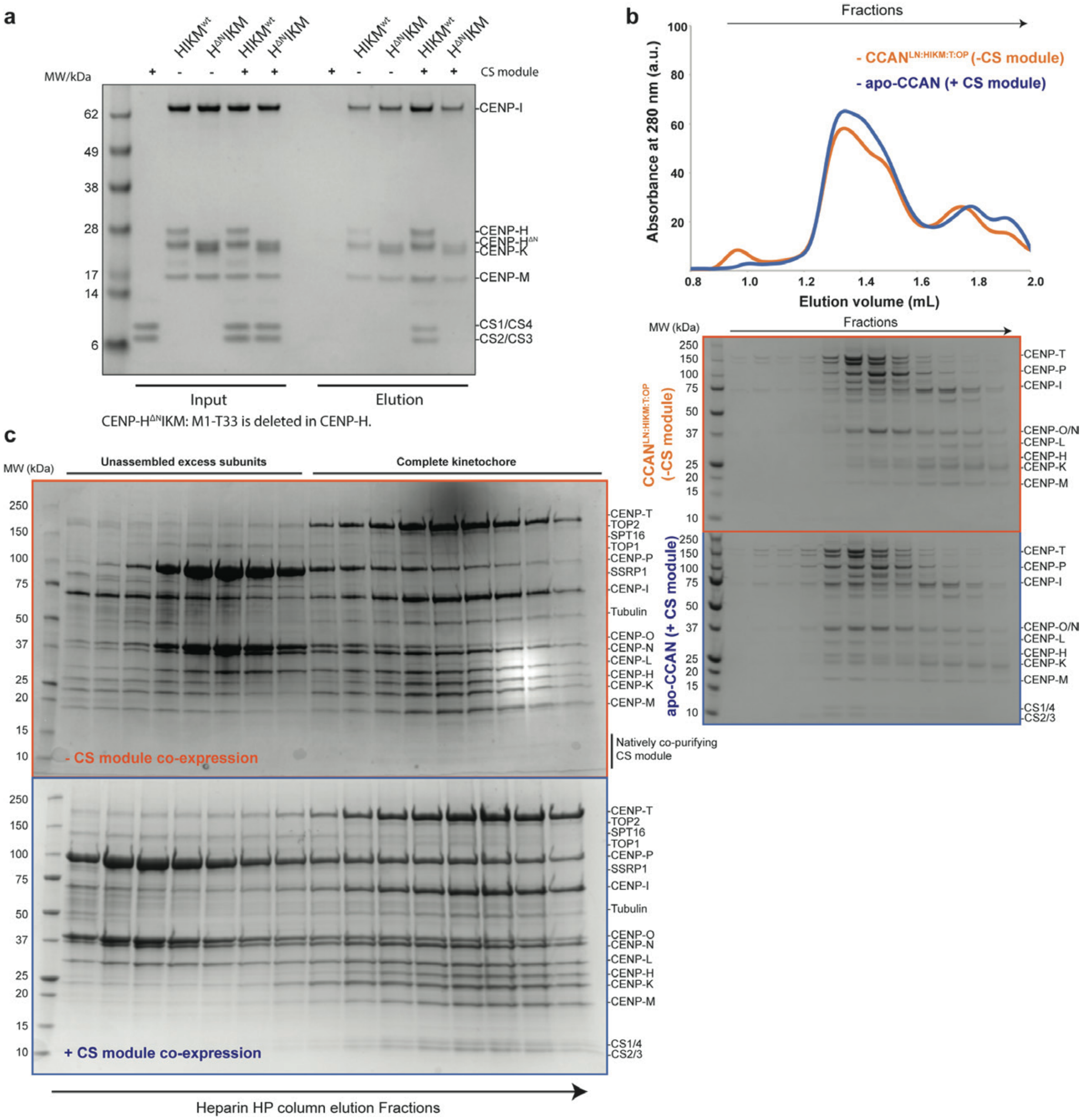


**Extended Data Fig. 6.** ***Bombyx mori*** **CS module biochemical characterization.**
**a,** StrepTactin pull-down assay. BmCENP-HIKM wild-type or bmCENP-H$^{\Delta N}$IKM mutant (where 1-33 amino acids in bmCENP-H protein are deleted) were immobilized on the StrepTactin resin via 2xStrepTag II at the N-terminus of bmCENP-I protein. A recombinantly purified CS module was added to the indicated reactions and assessed for its interaction with the bmCENP-HIKM module. **b,** BmCCAN reconstitution in the presence (apo-bmCCAN, blue line) or absence (bmCCAN$^{\mathrm{CENP,}}$ orange line) of the CS module using Superose 6 3.2/300 column. SDS-PAGE gel analysis of the SEC reconstitution assay. **c,** Heparin column reconstitution, as described in Extended Data Fig. 1c, d, in the absence (orange box and label, repeat of the gel shown in Extended Data Fig. 1d) and presence (blue box and label) of the CS module. The protein bands were identified and confirmed using mass spectrometry analysis (Methods).

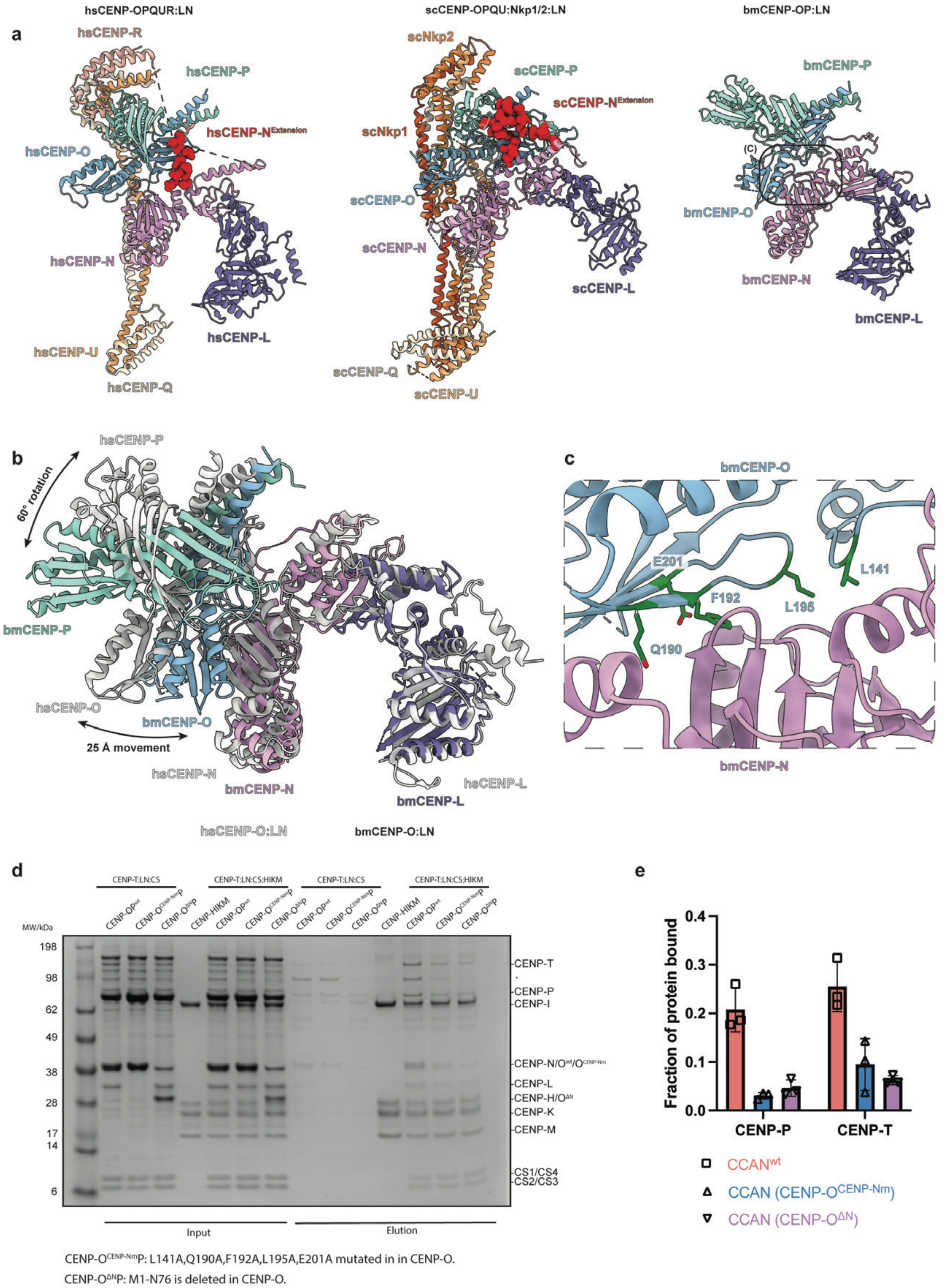
hsCENP-OPQUR:LN
scCENP-OPQU:Nkp1/2:LN
bmCENP-OP:LN
a
hsCENP-R
hsCENP-P
hsCENP-N^Extension
hsCENP-O
hsCENP-N
hsCENP-U
hsCENP-Q
hsCENP-L
scNkp2
scCENP-P
scCENP-N^Extension
scNkp1
scCENP-O
scCENP-N
scCENP-L
scCENP-Q
scCENP-U
bmCENP-P
(C)
bmCENP-O
bmCENP-N
bmCENP-L
b
hsCENP-P
60° rotation
bmCENP-P
hsCENP-O
bmCENP-O
25 Å movement
hsCENP-N
bmCENP-N
bmCENP-L
hsCENP-L
hsCENP-O:LN
bmCENP-O:LN
c
bmCENP-O
E201
L141
F192
L195
Q190
bmCENP-N
d
CENP-T:LN:CS
CENP-T:LN:CS:HIKM
CENP-T:LN:CS
CENP-T:LN:CS:HIKM
CENP-OP^wt
CENP-O^CENP-Nm P
CENP-O^ΔN P
CENP-HIKM
MW/kDa
198
98
62
49
38
28
17
14
6
CENP-T
CENP-P
CENP-I
CENP-N/O^wt/O^CENP-Nm
CENP-L
CENP-H/O^ΔN
CENP-K
CENP-M
CS1/CS4
CS2/CS3
Input
Elution
CENP-O^CENP-Nm P: L141A,Q190A,F192A,L195A,E201A mutated in in CENP-O.
CENP-O^ΔN P: M1-N76 is deleted in CENP-O.
e
Fraction of protein bound
0.4
0.3
0.2
0.1
0.0
CENP-P
CENP-T
CCAN^wt
CCAN (CENP-O^CENP-Nm)
CCAN (CENP-O^ΔN)

**Extended Data Fig. 7. Bombyx mori CS module biochemical characterization.**
**a, C**omparison of hsCENP-OPQUR-LN (PDB ID: 7R5S) [84], scCENP-OPQU-Nkp1-Nkp2-LN (PDB ID: 8OVW) [85], and bmCENP-OP-LN subcomplexes. CENP-$N^{Extension}$ that in human and yeast systems binds CENP-OP module is shown in red as space-filling model, **b,** Alignment of molecular models of insect and human CENP-LN-OP modules on CENP-L protein, with insect structure colored by subunit and human structure colored in grey. **c,** Detailed view of the CENP-O-CENP-N interface highlighting residues involved in the interaction between the two proteins that were mutated in CENP-$O^{Nm}$ mutant (L141A, Q190A, F192A, L195A, E201A). **d,** StrepTactin pull-down assay results where wild-type CENP-O or CENP-O mutants were used for the reconstitution of the bm-apoCCAN complex as indicated in the figure. The assay was repeated in technical triplicates. **e,** Quantification of the apo-bmCCAN StrepTactin pull-down assay shown in **d**. Band intensity for CENP-P and CENP-T was measured in the input and elution reactions in raw gel images to derive the fraction of the protein bound. The data point for each repeat is shown. Column height represents mean value and error bars represent standard deviation.

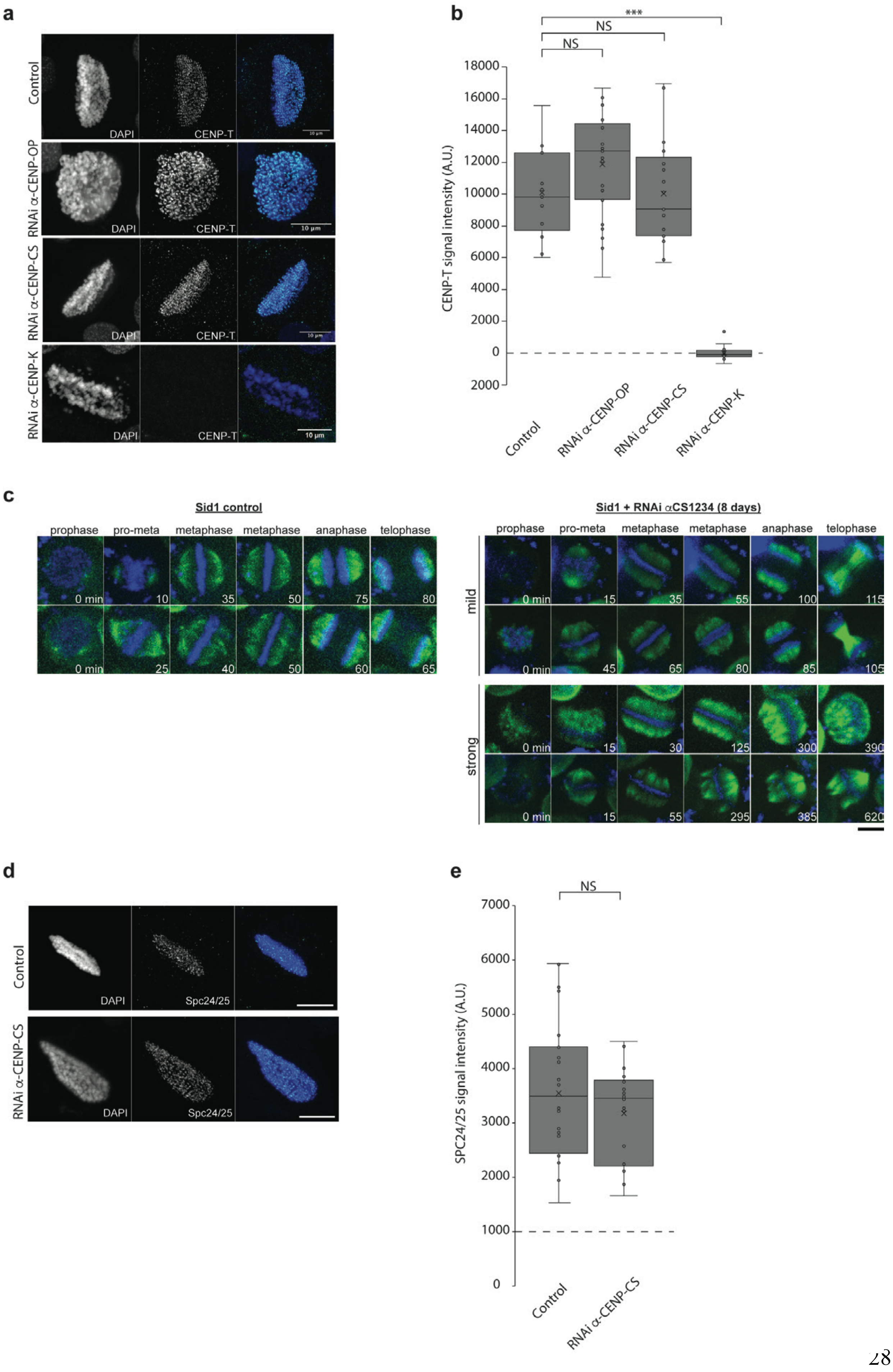
a
Control
RNAi α-CENP-OP
RNAi α-CENP-CS
RNAi α-CENP-K
DAPI
CENP-T
10 μm
b
***
NS
NS
CENP-T signal intensity (A.U.)
18000
16000
14000
12000
10000
8000
6000
4000
2000
0
2000
Control
RNAi α-CENP-OP
RNAi α-CENP-CS
RNAi α-CENP-K
c
Sid1 control
Sid1 + RNAi αCS1234 (8 days)
prophase
pro-meta
metaphase
metaphase
anaphase
telophase
mild
strong
0 min
d
Control
RNAi α-CENP-CS
DAPI
Spc24/25
e
NS
SPC24/25 signal intensity (A.U.)
7000
6000
5000
4000
3000
2000
1000
0
Control
RNAi α-CENP-CS

**Extended Data Fig. 8. BmCENP-T and bmSpc24/25 recruitment is unaffected upon CS or bmCENP-OP depletion.**

**a,** Representative images of mitotic *B. mori* cells showing the levels of endogenous CS with and without depletion of the CS module, bmCENP-OP or bmCENP-K kinetochore components. Scale bar: 10µm. b, Quantification of mean fluorescence intensity for bmCENP-T signals in the control or upon kinetochore depletions. Statistical significance was tested using a two-tailed students's t-test with unequal variance ($p \leq 0.0001$ for four stars, $p \leq 0.001$ for three stars, $p \leq 0.01$ for two stars and $p \leq 0.05$ for one star). **c,** Max-projection time-lapse images of the spindle (green) and chromosome (blue) dynamics for control and CS-depleted cells from mitotic stages prophase through telophase. Time is indicated in minutes. Scale bar, 10 µm. **d,** Representative images of mitotic *B. mori* cells showing the levels of endogenous Spc24/25 with and without depletion of the CS module, Scale bar: 10µm. **e,** Quantification of mean fluorescence intensity for bmSpc24/25 complex signals in the control or upon kinetochore depletions. Statistical significance was tested using a two-tailed students's t-test with unequal variance ($p \leq 0.0001$ for four stars, $p \leq 0.001$ for three stars, $p \leq 0.01$ for two stars and $p \leq 0.05$ for one star).

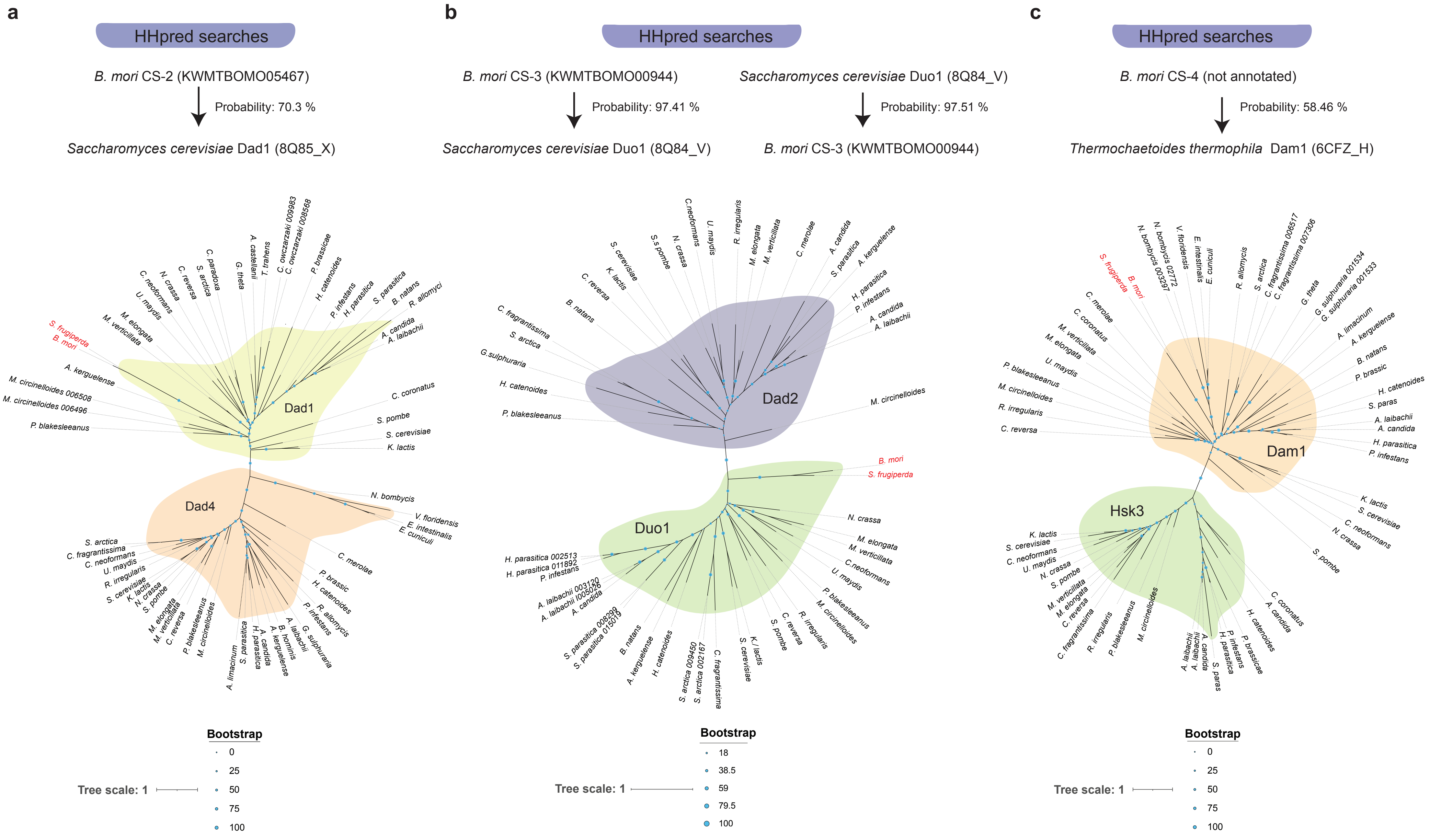
a
HHpred searches
B. mori CS-2 (KWMTBOMO05467)
Probability: 70.3 %
Saccharomyces cerevisiae Dad1 (8Q85_X)
Dad1
Dad4
Bootstrap
0
25
50
75
100
Tree scale: 1
b
HHpred searches
B. mori CS-3 (KWMTBOMO00944)
Probability: 97.41 %
Saccharomyces cerevisiae Duo1 (8Q84_V)
Saccharomyces cerevisiae Duo1 (8Q84_V)
Probability: 97.51 %
B. mori CS-3 (KWMTBOMO00944)
Dad2
Duo1
Bootstrap
18
38.5
59
79.5
100
Tree scale: 1
c
HHpred searches
B. mori CS-4 (not annotated)
Probability: 58.46 %
Thermochaetoides thermophila Dam1 (6CFZ_H)
Dam1
Hsk3
Bootstrap
0
25
50
75
100
Tree scale: 1

**Extended Data Fig. 9. Lepidopteran CS-2, CS-3, and CS-4 proteins are evolutionarily related to the Dam1, Duo1, and Dad1 paralogs of the Dam1/DASH complex.**

**a,** Top: HHpred hits of *B. mori* CS-2 protein against the PDB. Maximum-likelihood phylogeny of *B. mori* and *S. frugiperda* CS-2 proteins with Dad1 and Dad4. **b,** HHpred hits of *B. mori* CS-3 protein against the PDB along with reciprocal search. Maximum-likelihood phylogeny of *B. mori* and *S. frugiperda* CS-3 proteins with Dad2 and Duo1. **c,** HHpred hits of *B. mori* CS-4 protein against the PDB. Maximum-likelihood phylogeny of *B. mori* and *S. frugiperda* CS-4 proteins with Dam1 and Hsk3 paralogs. Dam1/DASH complex components are taken from[79]. Constructed using MAFFT merge and IQ-TREE (Methods). **d,** Multiple sequence alignment between various Dam1 homologs with *B. mori* and *S. frugiperda* CS-4 proteins showing conservation along the N-terminus but not C-terminus of Dam1. Mps1 (red) and Ipl1 (blue) phosphorylation sites are highlighted in *S. cerevisiae* Dam1[79] **e,** Multiple sequence alignment between various Duo1 homologs with *B. mori* and *S. frugiperda* CS3 proteins. Note that the alignments of the lepidopteran CS-3 proteins do not extend to the EB binding site described in *S. cerevisiae* Duo1 [79].

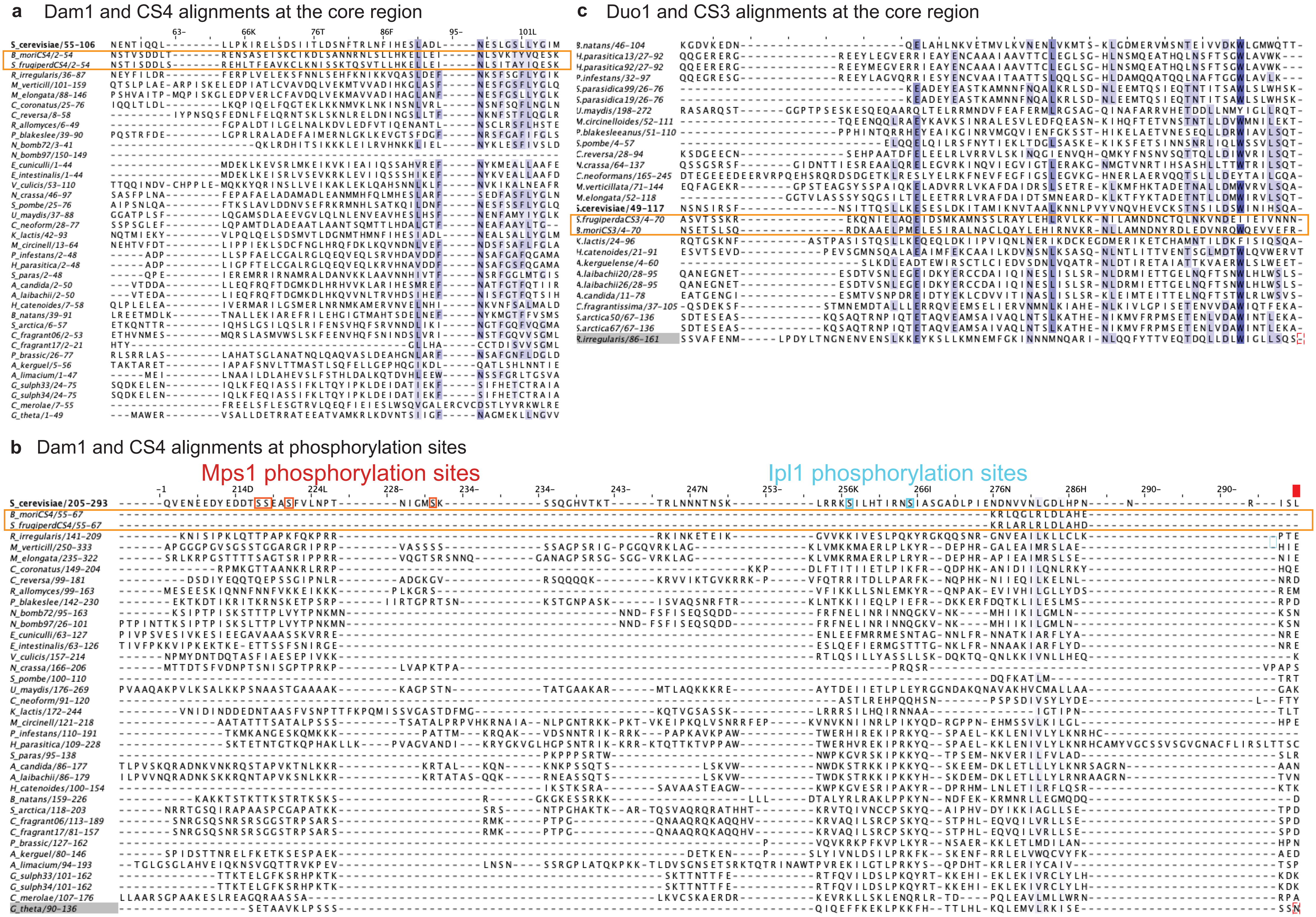
a Dam1 and CS4 alignments at the core region
c Duo1 and CS3 alignments at the core region
b Dam1 and CS4 alignments at phosphorylation sites
Mps1 phosphorylation sites
Ipl1 phosphorylation sites

**Extended Data Fig. 10. Lepidopteran CS-2, CS-3, and CS-4 proteins are evolutionarily related to the Dam1, Duo1, and Dad1 paralogs of the Dam1/DASH complex.**
**a,b,** Multiple sequence alignment between various Dam1 homologs with *B. mori* and *S. frugiperda* CS-4 proteins showing conservation along the N-terminus (the core region) but not the C-terminus (regulatory region) of Dam1. Mps1 (red) and Ipl1 (blue) phosphorylation sites are highlighted in *S. cerevisiae* Dam1 [79] **c,** Multiple sequence alignment between various Duo1 homologs with *B. mori* and *S. frugiperda* CS-3 proteins. Note that the alignments of the lepidopteran CS-3 proteins do not extend to the EB binding site described in *S. cerevisiae* Duo1 [79].

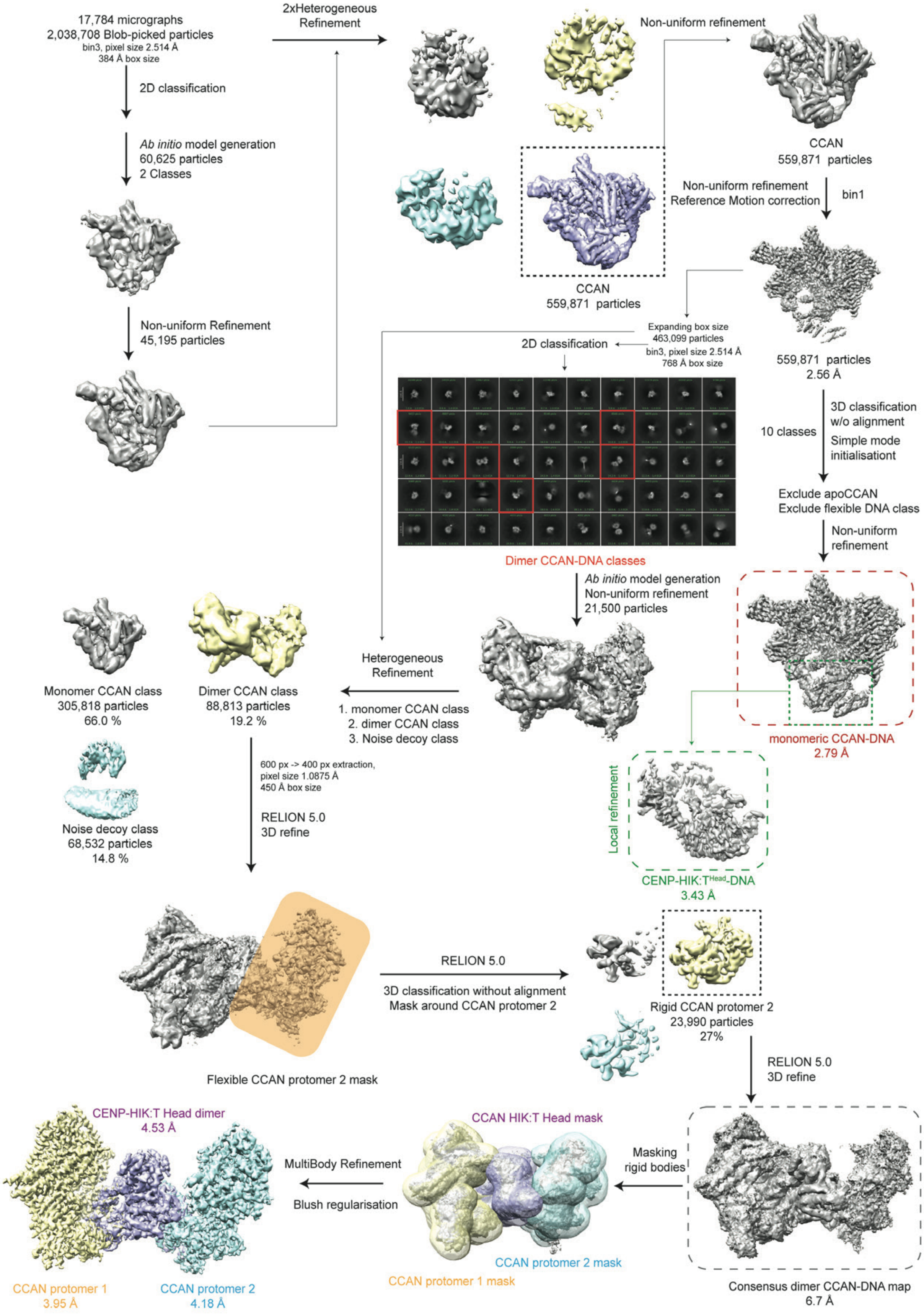
17,784 micrographs
2,038,708 Blob-picked particles
bin3, pixel size 2.514 Å
384 Å box size
2D classification
Ab initio model generation
60,625 particles
2 Classes
Non-uniform Refinement
45,195 particles
2xHeterogeneous Refinement
Non-uniform refinement
CCAN
559,871 particles
CCAN
559,871 particles
Non-uniform refinement
Reference Motion correction
bin1
559,871 particles
2.56 Å
Expanding box size
463,099 particles
bin3, pixel size 2.514 Å
768 Å box size
2D classification
Dimer CCAN-DNA classes
10 classes
3D classification
w/o alignment
Simple mode
initialisationt
Exclude apoCCAN
Exclude flexible DNA class
Non-uniform
refinement
Ab initio model generation
Non-uniform refinement
21,500 particles
Heterogeneous
Refinement
1. monomer CCAN class
2. dimer CCAN class
3. Noise decoy class
Monomer CCAN class
305,818 particles
66.0 %
Dimer CCAN class
88,813 particles
19.2 %
Noise decoy class
68,532 particles
14.8 %
600 px -> 400 px extraction,
pixel size 1.0875 Å
450 Å box size
RELION 5.0
3D refine
monomeric CCAN-DNA
2.79 Å
Local refinement
CENP-HIK:T^Head-DNA
3.43 Å
Flexible CCAN protomer 2 mask
RELION 5.0
3D classification without alignment
Mask around CCAN protomer 2
Rigid CCAN protomer 2
23,990 particles
27%
RELION 5.0
3D refine
Consensus dimer CCAN-DNA map
6.7 Å
Masking
rigid bodies
CCAN HIK:T Head mask
CCAN protomer 2 mask
CCAN protomer 1 mask
MultiBody Refinement
Blush regularisation
CENP-HIK:T Head dimer
4.53 Å
CCAN protomer 1
3.95 Å
CCAN protomer 2
4.18 Å

2

**Extended Data Fig. 11. BmCCAN-DNA complex cryoEM data processing.**
CryoEM data processing initially followed a similar processing pipeline to the apo-bmCCAN, as shown in Extended Data Fig. 2 and described in Methods. To identify dimeric CCAN-DNA complexes, the particle box size was expanded to select dimeric CCAN-DNA classes, which were used for *ab initio* model generation, and the model was further refined. Heterogeneous refinement was used to separate monomeric and dimeric CCAN-DNA classes. Dimeric CCAN-DNA complex was then further processed in RELION 5.0 to identify a class with rigid CCAN protomer 2 using classification without alignment. MultiBody refinement was then used to generate three high-resolution bodies of the CCAN-DNA dimer structure.

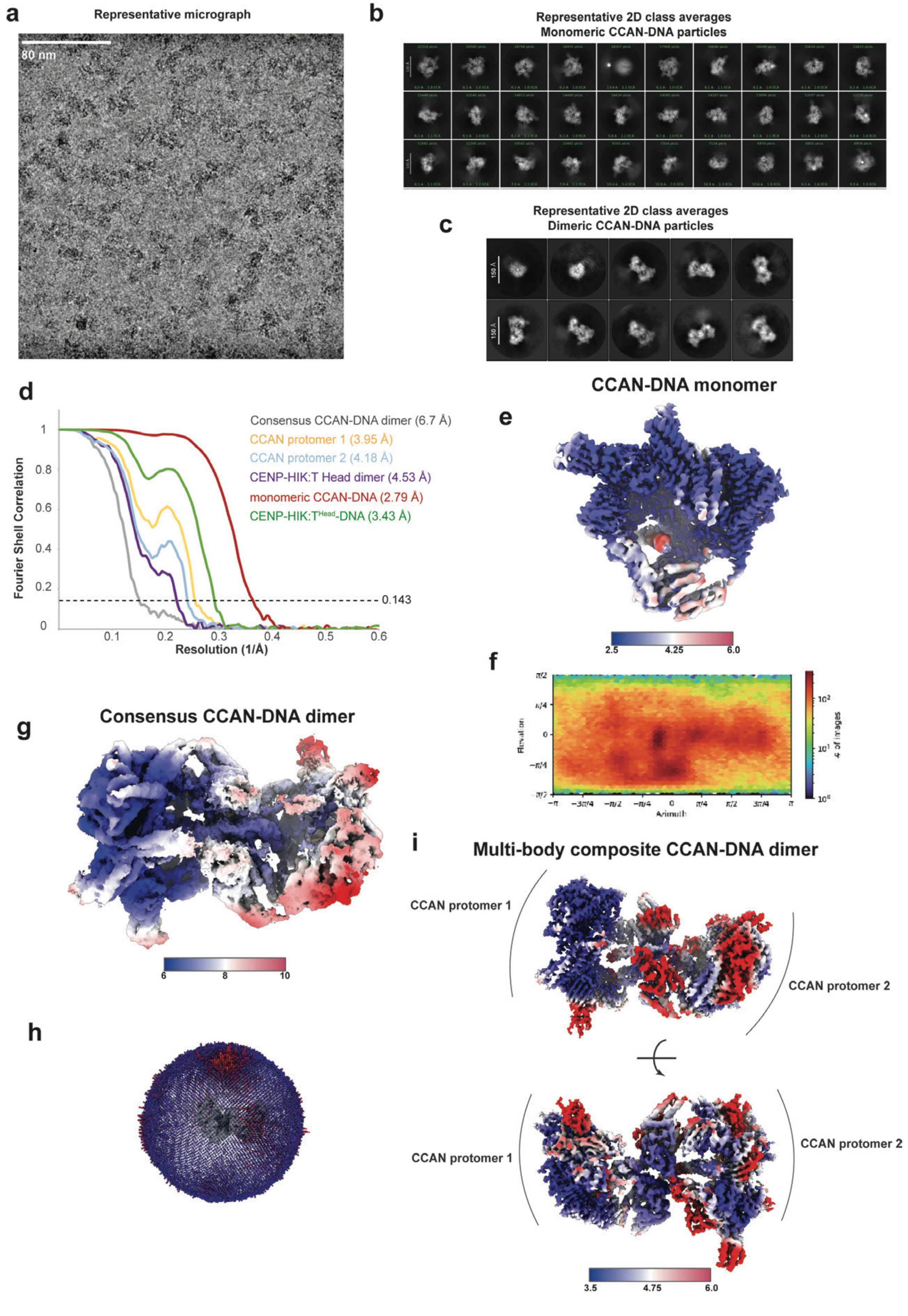
a
Representative micrograph
80 nm
b
Representative 2D class averages
Monomeric CCAN-DNA particles
c
Representative 2D class averages
Dimeric CCAN-DNA particles
150 Å
d
Consensus CCAN-DNA dimer (6.7 Å)
CCAN protomer 1 (3.95 Å)
CCAN protomer 2 (4.18 Å)
CENP-HIK:T Head dimer (4.53 Å)
monomeric CCAN-DNA (2.79 Å)
CENP-HIK:T^Head-DNA (3.43 Å)
Fourier Shell Correlation
Resolution (1/Å)
0.143
CCAN-DNA monomer
e
2.5
4.25
6.0
f
Elevation
Azimuth
### of images
g
Consensus CCAN-DNA dimer
6
8
10
h
i
Multi-body composite CCAN-DNA dimer
CCAN protomer 1
CCAN protomer 2
CCAN protomer 1
CCAN protomer 2
3.5
4.75
6.0

2

**Extended Data Fig. 12. BmCCAN-DNA complex cryoEM data presentation and validation.**

**a,** Representative cryoEM micrograph for the bmCCAN-DNA sample. **b,** Representative 2D class averages of the monomeric bmCCAN-DNA complex and **c,** dimeric CCAN-DNA complex generated using cryoSPARC. **d,** Fourier Shell Correlation (FSC) curves for all reported bmCCAN-DNA cryoEM reconstructions, with the grey dotted line representing 0.143 value. **e,** CryoEM reconstruction of the monomeric bmCCAN-DNA complex colored by resolution, with the resolution range shown below. **f,** Angular distribution of views for the monomeric bmCCAN-DNA complex. **g,** Consensus refinement for the dimeric CCAN-DNA complex colored by resolution, with the resolution range shown below. **h,** Angular distribution of views for the dimeric CCAN-DNA complex. **i,** Composite multi-body map of the dimeric bmCCAN-DNA complex colored by resolution as determined for each individual body.

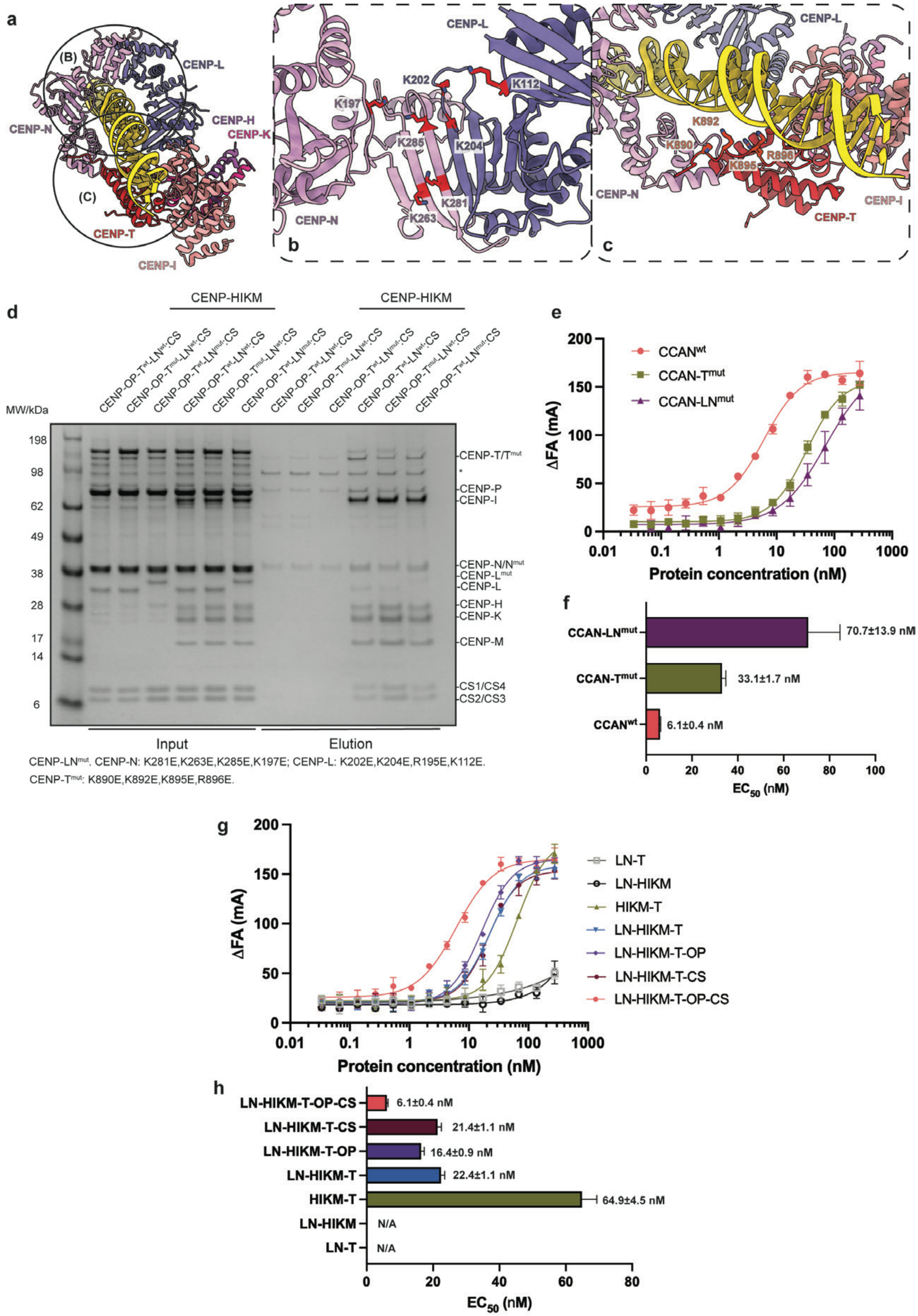

a
CENP-L
CENP-N
CENP-H
CENP-K
CENP-T
CENP-I
(B)
(C)
b
K197
K202
K112
K285
K204
K281
K263
c
K892
K890
K895
R896
d
CENP-HIKM
CENP-HIKM
MW/kDa
Input
Elution
CENP-T/T^mut
CENP-P
CENP-I
CENP-N/N^mut
CENP-L^mut
CENP-L
CENP-H
CENP-K
CENP-M
CS1/CS4
CS2/CS3
CENP-LN^mut: CENP-N: K281E,K263E,K285E,K197E; CENP-L: K202E,K204E,R195E,K112E.
CENP-T^mut: K890E,K892E,K895E,R896E.
e
ΔFA (mA)
Protein concentration (nM)
CCAN^wt
CCAN-T^mut
CCAN-LN^mut
f
70.7±13.9 nM
33.1±1.7 nM
6.1±0.4 nM
EC50 (nM)
g
LN-T
LN-HIKM
HIKM-T
LN-HIKM-T
LN-HIKM-T-OP
LN-HIKM-T-CS
LN-HIKM-T-OP-CS
h
6.1±0.4 nM
21.4±1.1 nM
16.4±0.9 nM
22.4±1.1 nM
64.9±4.5 nM
N/A
N/A


2

**Extended Data Fig. 13. CENP-LN and CENP-T are major contributors to DNA binding.** **a,** A molecular model of the DNA bound to monomeric bmCCAN, with a focus on DNA binding subunits. **b,** A close-up of the bmCENP-LN surface. Residues mutated in CENP-LN$^{mut}$ protein that interact with DNA are highlighted in red and shown as sticks. **c,** A close-up of CENP-T bound to DNA. Residues mutated in CENP-T$^{mut}$ protein that interact with DNA are highlighted in red and shown as sticks. **d,** StrepTactin pull-down assay results with CENP-LN$^{mut}$, CENP-T$^{mut}$ and wild-type proteins showing no defect in bmCCAN assembly in these mutants. **e,** Results of the fluorescence polarization (FP) assay. The assay was repeated at least in technical triplicates with mean value and standard deviation values shown. The best-fit regression analysis curve is fitted through the data points to calculate half-maximal effective concentration ($EC_{50}$) values. **f,** $EC_{50}$ values calculated from FP assay shown in e, are plotted with bar height indicating the mean value and the error bars indicating the standard error of the mean (SEM). $EC_{50}$ with SEM values for each condition are also stated in the figure. **g,** FP assay results where different bmCCAN components were added, as indicated on the figure. The assay was repeated at least in technical triplicates with mean value and standard deviation values shown. **h,** $EC_{50}$ values calculated from the FP assay shown in g, are plotted with bar height indicating the mean value and the error bars indicating the standard error of the mean (SEM).

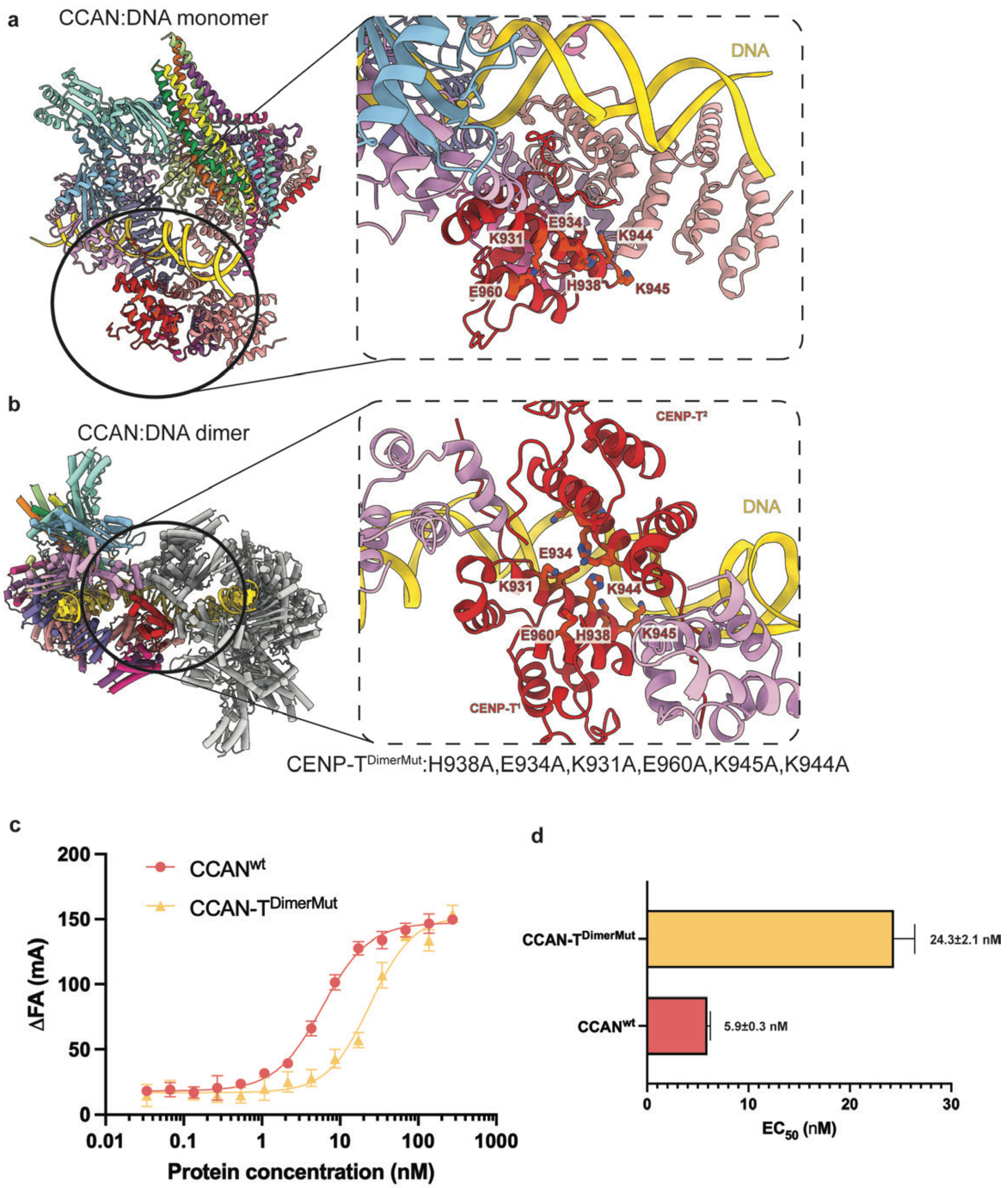


**Extended Data Fig. 14. CENP-T dimerization is important for DNA binding.**
**a,** A molecular model of monomeric bmCCAN-DNA complex, with inset showing residues mutated in the CENP-$T^{DimerMut}$ mutant (K931A, E934A, H938A, K944A, K945A, E960A). These residues are surface-exposed and far removed from the DNA. **b,** A molecular model of dimeric bmCCAN-DNA complex, with the inset showing residues mutated in the CENP-$T^{DimerMut}$ mutant. **c,** Results of the fluorescence polarization (FP) assay that compares wild-type bmCCAN or bmCCAN with CENP-$T^{DimerMut}$ (CCAN-$T^{DimerMut}$). The assay was repeated at least in technical triplicates with mean value and standard deviation values shown. The best-fit regression analysis curve is fitted through the data points to calculate half-maximal effective concentration ($EC_{50}$) values. **d,** $EC_{50}$ values calculated from FP assay shown in c, are plotted with bar height indicating the mean value and the error bars indicating the standard error of the mean (SEM). $EC_{50}$ with SEM values for each condition are also stated in the figure.

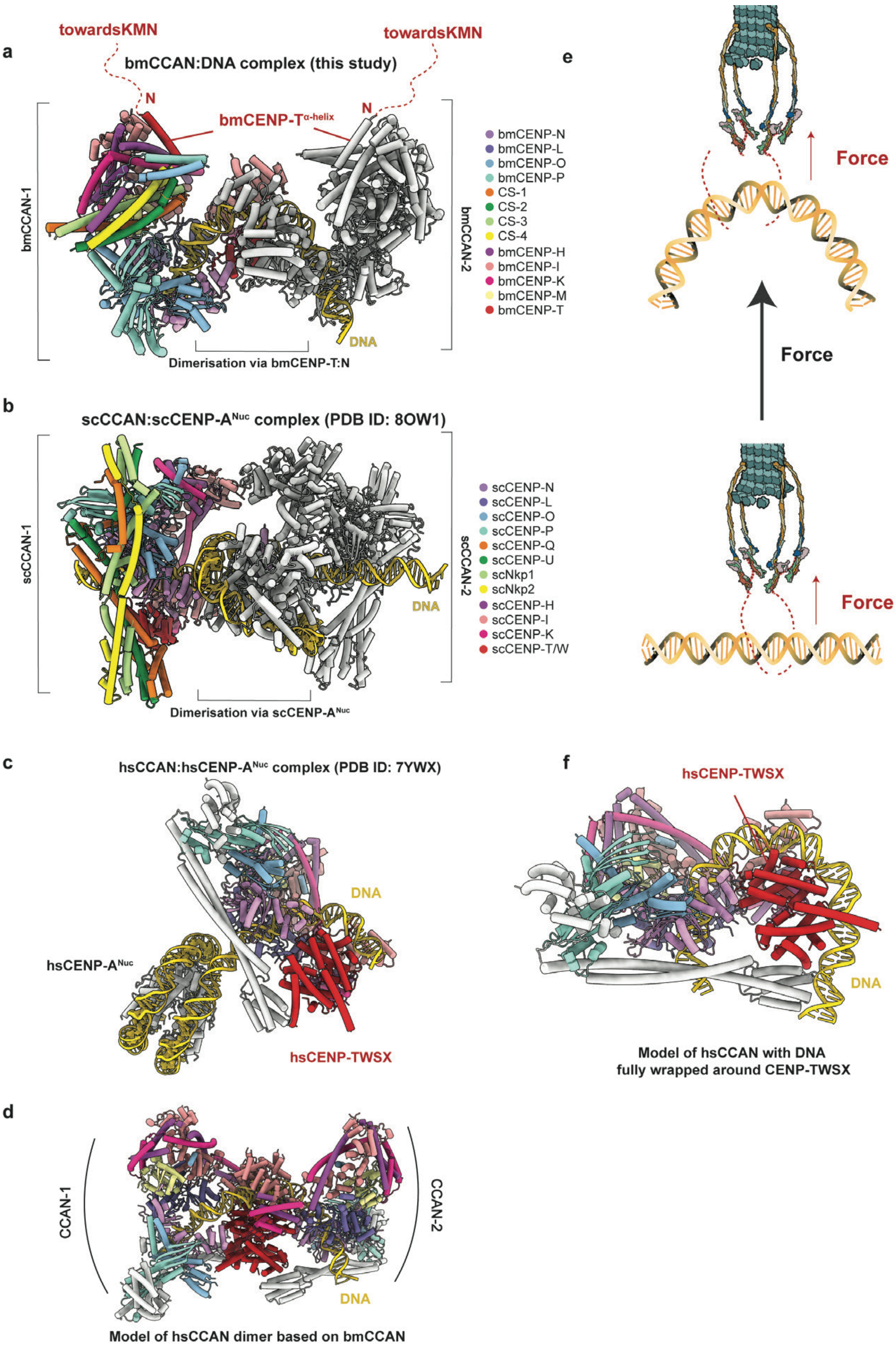
a
bmCCAN:DNA complex (this study)
towardsKMN
towardsKMN
N
N
bmCENP-T$^{\alpha\text{-helix}}$
bmCCAN-1
bmCCAN-2
DNA
Dimerisation via bmCENP-T:N
bmCENP-N
bmCENP-L
bmCENP-O
bmCENP-P
CS-1
CS-2
CS-3
CS-4
bmCENP-H
bmCENP-I
bmCENP-K
bmCENP-M
bmCENP-T
b
scCCAN:scCENP-A$^{Nuc}$ complex (PDB ID: 8OW1)
scCCAN-1
scCCAN-2
DNA
Dimerisation via scCENP-A$^{Nuc}$
scCENP-N
scCENP-L
scCENP-O
scCENP-P
scCENP-Q
scCENP-U
scNkp1
scNkp2
scCENP-H
scCENP-I
scCENP-K
scCENP-T/W
c
hsCCAN:hsCENP-A$^{Nuc}$ complex (PDB ID: 7YWX)
DNA
hsCENP-A$^{Nuc}$
hsCENP-TWSX
d
CCAN-1
CCAN-2
DNA
Model of hsCCAN dimer based on bmCCAN
e
Force
Force
Force
f
hsCENP-TWSX
DNA
Model of hsCCAN with DNA
fully wrapped around CENP-TWSX

**Extended Data Fig. 15. BmCCAN forms an alternative point-centromere-like kinetochore structure.**

**a,** Molecular model of the dimeric bmCCAN-DNA complex determined in this study, emphasizing the bmCENP-$T^{\alpha\text{-helix}}$ and showing that its N-terminus is facing away from the complex. The dotted line represents the N-terminus of bmCENP-T where putative bmKMN binding sites are located. **b,** Aligned molecular model of the complete yeast point-centromere (PDB ID: 8OW1), with only scCCAN and scCENP-$A^{Nuc}$ shown (CBF1 and CBF3 complexes are hidden for clarity). The scCCAN model was aligned against bmCCAN-1 in panel a, and colored using the bmCCAN color code. **c,** Aligned molecular model of the human hsCCAN-CENP-$A^{Nuc}$ complex (PDB ID: 7YWX), with CENP-$A^{Nuc}$ and hsCENP-TWSX modules highlighted. HsCCAN model is colored using bmCCAN color code. **d,** Model of the dimeric human CCAN structure based on the bmCCAN structure. The model was generated by aligning two copies of hsCCAN to bmCCAN-1 and bmCCAN-2, specifically aligning hsCENP-T protein with bmCENP-T in each CCAN protomer. The DNA was preserved from the hsCCAN structure, and it naturally extends in a manner similar to the bmCCAN:DNA complex. **e,** Cartoon schematic of events that are likely to happen when spindle forces are applied to the DNA, suggesting a bending in the DNA due to its low persistence length. The bent DNA resembles the DNA loop that is held by dimeric bmCCAN-DNA complex. **f,** hsCCAN monomer modelled with DNA being completely wrapped around hsCENP-TWSX complex, which also results in hsCCAN holding a DNA loop, similar to bmCCAN structure.

| | bmCENP-LN-HIKM-T EMDB-70567 PDB: 9OKK | bmCENP-LN-HIKM-T-OP EMDB-70561 PDB: 9OKE | apo-bmCCAN EMDB-70568 PDB: 9OKL | bmCCAN-DNA (monomer) EMDB-70560 PDB: 9OKD | bmCCAN-DNA (dimer) EMDB-70558 PDB: 9OKB |
|---|---|---|---|---|---|
| **Data collection and processing** | | | | | |
| Exposures | 19,201 | 19,201 | 19,201 | 24,025 | 24,025 |
| Magnification | 105,000 | 105,000 | 105,000 | 105,000 | 105,000 |
| Voltage (kV) | 300 | 300 | 300 | 300 | 300 |
| Electron exposure (e–/Å$^2$) | 62 | 62 | 62 | 60 | 60 |
| Defocus range (μm) | 1-3 | 1-3 | 1-3 | 1-3 | 1-3 |
| Pixel size (Å) | 0.838 | 0.838 | 0.838 | 0.838 | 0.838 |
| Symmetry imposed | C1 | C1 | C1 | C1 | C1 |
| Initial particle images (no.) | 4,100,926 | 4,100,926 | 4,100,926 | 2,038,708 | 2,038,708 |
| Final particle images (no.) | 237,375 | 328,750 | 265,941 | 241,975 | 23,990 |
| Map resolution (Å) | 3.23 | 3.24 | 3.0 | 2.83 | 2.44 |
| FSC threshold | 0.143 | 0.143 | 0.143 | 0.143 | 0.143 |
| Map resolution range (Å) | 2.6-20 | 2.6-20 | 2.5-20 | 2.83-20 | 2.44-20 |
| **Refinement** | | | | | |
| Initial model used (PDB code) | AlphaFold2, *Ab initio* | AlphaFold2, *Ab initio* | AlphaFold2, *Ab initio* | AlphaFold2, *Ab initio* | AlphaFold2, *Ab initio* |
| Model resolution (Å) | 3.2 | 3.2 | 3.0 | 2.83 | 2.44 |
| FSC threshold | | | | | |
| Model resolution range (Å) | 3.1-20 | 3.1-20 | 2.5-20 | 2.7-20 | 2.2-20 |
| Map sharpening *B* factor (Å$^2$) | - 126.3 | - 133.0 | -92.1 | -30 | -50 |
| Model composition | | | | | |
| Non-hydrogen atoms | 18687 | 28018 | 39509 | 42619 | 84496 |
| Protein residues | 1168 | 1748 | 2569 | 2690 | 5368 |
| Nucleotide | 0 | 0 | 0 | 72 | 116 |
| *B* factors (Å$^2$) | | | | | |
| Protein | 155.63 | 184.96 | 188.73 | 195.75 | 364.03 |
| Nucleotide | N/A | N/A | N/A | 917.56 | 253.81 |
| R.m.s. deviations | | | | | |
| Bond lengths (Å) | 0.003 | 0.004 | 0.002 | 0.003 | 0.002 |
| Bond angles (°) | 0.502 | 0.911 | 0.403 | 0.451 | 0.475 |
| Validation | | | | | |
| MolProbity score | 1.41 | 1.63 | 1.61 | 1.95 | 2.00 |
| Clashscore | 4.54 | 7.95 | 10.71 | 14.62 | 16.86 |
| Poor rotamers (%) | 0.00 | 0.00 | 0.90 | 0.74 | 0.19 |
| Ramachandran plot | | | | | |
| Favored (%) | 96.94 | 96.85 | 97.70 | 95.96 | 96.08 |
| Allowed (%) | 3.06 | 3.03 | 2.22 | 4.01 | 3.83 |
| Disallowed (%) | 0.00 | 0.12 | 0.08 | 0.04 | 0.09 |

**Extended Data Table 1. Cryo-EM data collection, refinement and validation statistics.**

**Supplementary Movie 1. BmCCAN-DNA structure overview.**

**Supplementary Data 1. Mass-spectrometry analysis of the bmCENP-O/P and CS-1/2 pull-down experiments.**

**Supplementary Data 2. Solution mass-spectrometry analysis of the heparin-purified bmCCAN sample.**

**Supplementary References.**


56 Punjani, A., Rubinstein, J. L., Fleet, D. J. & Brubaker, M. A. cryoSPARC: algorithms for rapid unsupervised cryo-EM structure determination. *Nat Methods* **14**, 290-296 (2017). https://doi.org/10.1038/nmeth.4169

57 Adams, P. D. *et al.* PHENIX: building new software for automated crystallographic structure determination. *Acta Crystallogr D Biol Crystallogr* **58**, 1948-1954 (2002). https://doi.org/10.1107/s0907444902016657

58 Burt, A. *et al.* An image processing pipeline for electron cryo-tomography in RELION-5. *FEBS Open Bio* **14**, 1788-1804 (2024). https://doi.org/10.1002/2211-5463.13873

59 Kimanius, D. *et al.* Data-driven regularization lowers the size barrier of cryo-EM structure determination. *Nat Methods* **21**, 1216-1221 (2024). https://doi.org/10.1038/s41592-024-02304-8

60 Jumper, J. *et al.* Highly accurate protein structure prediction with AlphaFold. *Nature* **596**, 583-589 (2021). https://doi.org/10.1038/s41586-021-03819-2

61 Emsley, P., Lohkamp, B., Scott, W. G. & Cowtan, K. Features and development of Coot. *Acta Crystallogr D Biol Crystallogr* **66**, 486-501 (2010). https://doi.org/10.1107/S0907444910007493

62 Jamali, K. *et al.* Automated model building and protein identification in cryo-EM maps. *Nature* **628**, 450-457 (2024). https://doi.org/10.1038/s41586-024-07215-4

63 Remmert, M., Biegert, A., Hauser, A. & Soding, J. HHblits: lightning-fast iterative protein sequence searching by HMM-HMM alignment. *Nat Methods* **9**, 173-175 (2011). https://doi.org/10.1038/nmeth.1818

64 Cortes-Silva, N. *et al.* CenH3-Independent Kinetochore Assembly in Lepidoptera Requires CCAN, Including CENP-T. *Curr Biol* **30**, 561-572 e510 (2020). https://doi.org/10.1016/j.cub.2019.12.014

65 Sissoko, G. B., Tarasovetc, E. V., Marescal, O., Grishchuk, E. L. & Cheeseman, I. M. Higher-order protein assembly controls kinetochore formation. *Nature cell biology* **26**, 45-56 (2024). https://doi.org/10.1038/s41556-023-01313-7

66 Schindelin, J. *et al.* Fiji: an open-source platform for biological-image analysis. *Nat Methods* **9**, 676-682 (2012). https://doi.org/10.1038/nmeth.2019

67 Pettersen, E. F. *et al.* UCSF ChimeraX: Structure visualization for researchers, educators, and developers. *Protein Sci* **30**, 70-82 (2021). https://doi.org/10.1002/pro.3943

68 Mon, H. *et al.* Effective RNA interference in cultured silkworm cells mediated by overexpression of Caenorhabditis elegans SID-1. *RNA Biol* **9**, 40-46 (2012). https://doi.org/10.4161/rna.9.1.18084

69 Poullet, P., Carpentier, S. & Barillot, E. myProMS, a web server for management and validation of mass spectrometry-based proteomic data. *Proteomics* **7**, 2553-2556 (2007). https://doi.org/10.1002/pmic.200600784

70 Perez-Riverol, Y. *et al.* The PRIDE database resources in 2022: a hub for mass spectrometry-based proteomics evidences. *Nucleic Acids Res* **50**, D543-D552 (2022). https://doi.org/10.1093/nar/gkab1038

71 Senaratne, A. P. & Drinnenberg, I. A. All that is old does not wither: Conservation of outer kinetochore proteins across all eukaryotes? *J Cell Biol* **216**, 291-293 (2017). https://doi.org/10.1083/jcb.201701025

72 Bosch Grau, M. *et al.* Tubulin glycylases and glutamylases have distinct functions in stabilization and motility of ependymal cilia. *J Cell Biol* **202**, 441-451 (2013). https://doi.org/10.1083/jcb.201305041

73 Vanpoperinghe, L. *et al.* Live-cell imaging reveals square shape spindles and long mitosis duration in the silkworm holocentric cells. *MicroPubl Biol* **2021** (2021). https://doi.org/10.17912/micropub.biology.000441
74 Skene, P. J. & Henikoff, S. A simple method for generating high-resolution maps of genome-wide protein binding. *Elife* **4**, e09225 (2015). https://doi.org/10.7554/eLife.09225
75 Langmead, B. & Salzberg, S. L. Fast gapped-read alignment with Bowtie 2. *Nat Methods* **9**, 357-359 (2012). https://doi.org/10.1038/nmeth.1923
76 Ramirez, F. *et al.* deepTools2: a next generation web server for deep-sequencing data analysis. *Nucleic Acids Res* **44**, W160-165 (2016). https://doi.org/10.1093/nar/gkw257
77 Robinson, J. T. *et al.* Integrative genomics viewer. *Nat Biotechnol* **29**, 24-26 (2011). https://doi.org/10.1038/nbt.1754
78 Soding, J., Biegert, A. & Lupas, A. N. The HHpred interactive server for protein homology detection and structure prediction. *Nucleic Acids Res* **33**, W244-248 (2005). https://doi.org/10.1093/nar/gki408
79 van Rooijen, L. E., Tromer, E. C., van Hooff, J. J. E., Kops, G. & Snel, B. Increased Sampling and Intracomplex Homologies Favor Vertical Over Horizontal Inheritance of the Dam1 Complex. *Genome Biol Evol* **15** (2023). https://doi.org/10.1093/gbe/evad017
80 Katoh, K. & Standley, D. M. MAFFT multiple sequence alignment software version 7: improvements in performance and usability. *Mol Biol Evol* **30**, 772-780 (2013). https://doi.org/10.1093/molbev/mst010
81 Minh, B. Q. *et al.* IQ-TREE 2: New Models and Efficient Methods for Phylogenetic Inference in the Genomic Era. *Mol Biol Evol* **37**, 1530-1534 (2020). https://doi.org/10.1093/molbev/msaa015
82 Kalyaanamoorthy, S., Minh, B. Q., Wong, T. K. F., von Haeseler, A. & Jermiin, L. S. ModelFinder: fast model selection for accurate phylogenetic estimates. *Nat Methods* **14**, 587-589 (2017). https://doi.org/10.1038/nmeth.4285
83 Letunic, I. & Bork, P. Interactive Tree of Life (iTOL) v6: recent updates to the phylogenetic tree display and annotation tool. *Nucleic Acids Res* **52**, W78-W82 (2024). https://doi.org/10.1093/nar/gkae268
84 Yatskevich, S. *et al.* Structure of the human inner kinetochore bound to a centromeric CENP-A nucleosome. *Science* **376**, 844-852 (2022). https://doi.org/10.1126/science.abn3810
85 Dendooven, T. *et al.* Cryo-EM structure of the complete inner kinetochore of the budding yeast point centromere. *Sci Adv* **9**, eadg7480 (2023). https://doi.org/10.1126/sciadv.adg7480
